# Cross-species transcriptomic and epigenomic analysis reveals key regulators of injury response and neuronal regeneration in vertebrate retinas

**DOI:** 10.1101/717876

**Authors:** Thanh Hoang, Jie Wang, Patrick Boyd, Fang Wang, Clayton Santiago, Lizhi Jiang, Manuela Lahne, Sooyeon Yoo, Levi J. Todd, Cristian Saez, Casey Keuthan, Isabella Palazzo, Natalie Squires, Warren A. Campbell, Meng Jia, Fatemeh Rajaii, Trisha Parayil, Vickie Trinh, Dong Won Kim, Guohua Wang, John Ash, Andy J. Fischer, David R. Hyde, Jiang Qian, Seth Blackshaw

**Affiliations:** Solomon H. Snyder Department of Neuroscience, Johns Hopkins University School of Medicine, Baltimore, MD, 21205, USA; Department of Ophthalmology, Johns Hopkins University School of Medicine, Baltimore, MD, 21205, USA; Department of Neurology, Johns Hopkins University School of Medicine, Baltimore, MD, 21205, USA; Center for Human Systems Biology, Johns Hopkins University School of Medicine, Baltimore, MD, 21205, USA; Institute for Cell Engineering, Johns Hopkins University School of Medicine, Baltimore, MD, 21205, USA; Kavli Neuroscience Discovery Institute, Johns Hopkins University School of Medicine, Baltimore, MD, 21205, USA; Department of Biological Sciences, University of Notre Dame, Notre Dame, IN, 46556 USA; Center for Stem Cells and Regenerative Medicine, University of Notre Dame, Notre Dame, IN, 46556 USA; Center for Zebrafish Research, University of Notre Dame, Notre Dame, IN, 46556 USA; Department of Neuroscience, Ohio State University Wexner Medical Center, Columbus, OH, 43210 USA; Department of Ophthalmology, University of Florida School of Medicine, Gainesville, FL, 32610 USA

## Abstract

Injury induces retinal Müller glia of cold-blooded, but not mammalian, vertebrates to regenerate neurons. To identify gene regulatory networks that control neuronal reprogramming in retinal glia, we comprehensively profiled injury-dependent changes in gene expression and chromatin accessibility in Müller glia from zebrafish, chick and mice using bulk RNA-Seq and ATAC-Seq, as well as single-cell RNA-Seq. Cross-species integrative analysis of these data, together with functional validation, identified evolutionarily conserved and species-specific gene networks controlling glial quiescence, gliosis and neurogenesis. In zebrafish and chick, transition from the resting state to gliosis is essential for initiation of retinal regeneration, while in mice a dedicated network suppresses neurogenic competence and restores quiescence. Selective disruption of NFI family transcription factors, which maintain and restore quiescence, enables Müller glia to proliferate and generate neurons in adult mice following retinal injury. These findings may aid in the design of cell-based therapies aimed at restoring retinal neurons lost to degenerative disease.

**Summary sentence:** This study identifies gene regulatory networks controlling proliferative and neurogenic competence in retinal Müller glia.

## Introduction

The degeneration of retinal neurons is the key pathological feature of blinding diseases such as macular degeneration, retinitis pigmentosa and glaucoma. While there are therapies that can slow the progression of vision loss, there is currently no effective way to restore retinal neurons lost to disease. Although mammals lack the ability to regenerate retinal neurons, retinal glia in many other species retain some degree of injury-induced neurogenic competence. In zebrafish, loss of retinal neurons robustly stimulates regeneration of all neuronal subtypes even into adulthood (*1, 2*). This results from Müller glia (MG) reprogramming and reentering the cell cycle, producing neurogenic Müller glia-derived progenitor cells (MGPC). In contrast, although MG of post-hatch chicks proliferate robustly in response to injury, they retain limited neurogenic competence, and only give rise to small numbers of inner retinal neurons (*3, 4*). In mice and humans, MG do not even proliferate robustly in response to neuronal injury, and altogether lack neurogenic competence.

Studies in zebrafish identified a number of genes that are rapidly induced in MG following injury and regulate neurogenic competence. Multiple extrinsic signaling pathways, including Wnt, BMP, FGF, TNFα, and cytokines, were found to regulate MGPC formation (*5–14*). These induce neurogenic factor genes, such as *ascl1a* (*15*) and the RNA binding protein *lin28a* (*16*), which are necessary for MG reprogramming. Most of these signaling pathways are activated in response to injury in zebrafish, chick and mouse, but they fail to induce *Ascl1* or *Lin28a* in mammals. Interestingly, forced expression of *Ascl1* in mouse MG, in combination with neuronal injury and histone deacetylase inhibitor treatment, can induce direct conversion of MG to inner retinal cells that show neuronal morphology and respond to light (*17, 18*). Conversely, inhibition of the Hippo pathway stimulates MG proliferation in mice following injury, but does not induce neurogenic competence (*5, 19*). This suggests that key molecular components of MG reprogramming remain to be identified. A more complete understanding of the genetic and epigenetic barriers to induction of proliferative and neurogenic competence in mammalian MG can potentially inform and improve approaches for cell-based therapies aimed at treating retinal dystrophies.

To comprehensively identify transcriptional and epigenetic regulators of neurogenic competence in MG, we profiled mRNA levels and chromatin accessibility in zebrafish, chick and mouse in response to multiple neuronal injury models, as well as injury-independent treatment with exogenous factors that induce MG reprogramming (Fig. 1A). By performing both bulk and single-cell RNA-Seq (scRNA-Seq) analysis of MG, in combination with ATAC-Seq profiling, we reconstructed both evolutionarily conserved and species-specific gene regulatory networks controlling glial quiescence, gliosis, proliferation and neurogenic competence (Fig. 1B). Our findings highlight key similarities and differences in the response of MG to injury in species with dramatically different regenerative abilities, and identify mechanisms that actively suppress the formation of MGPCs in mammals.

**Fig. 1.**
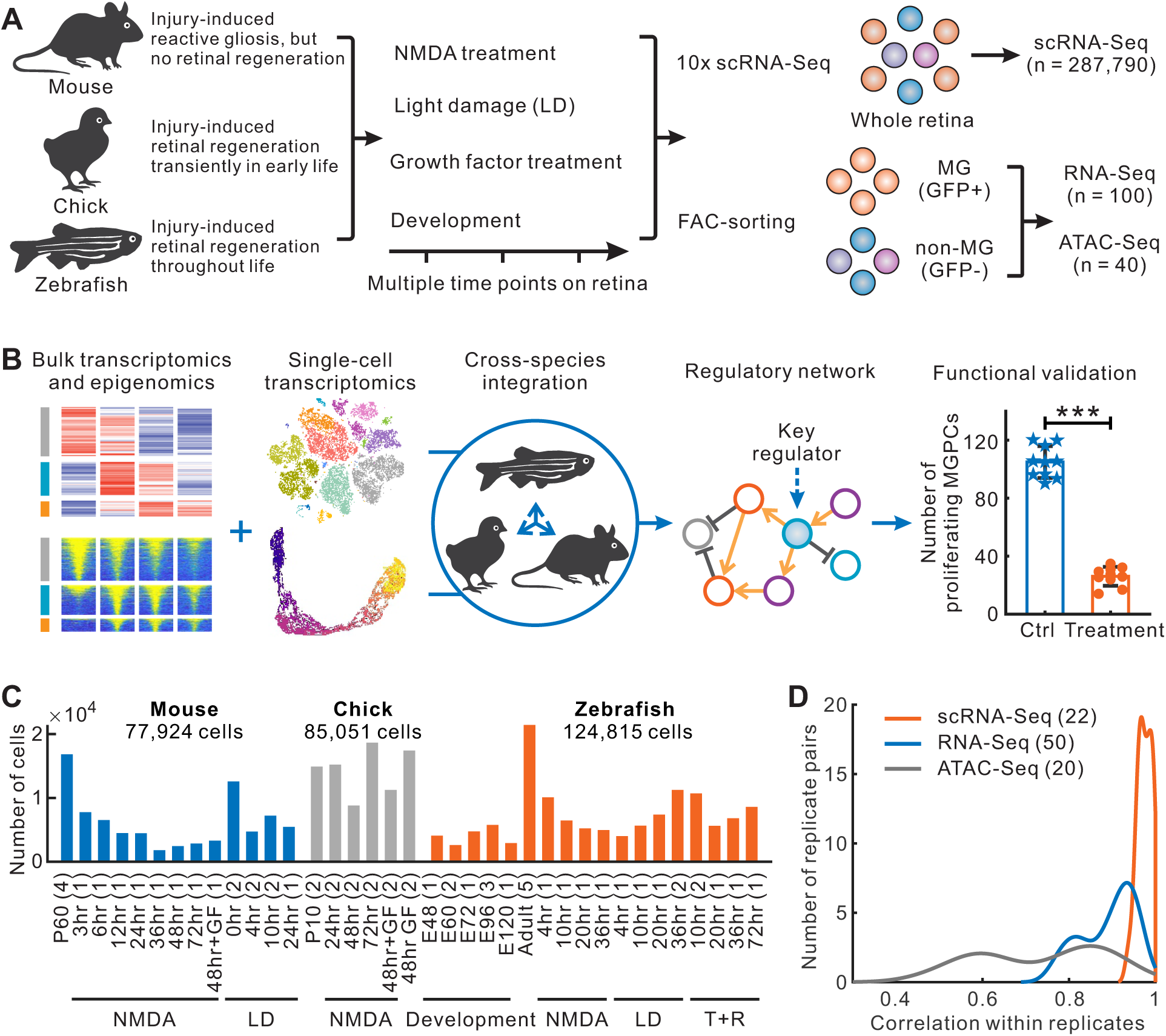
Overview of study. **(A)** Schematics of the project. We performed single-cell RNA-Seq (scRNA-Seq) for the whole retina, bulk RNA-Seq and ATAC-Seq for FAC-sorting (fluorescence activated cell sorting) Müller glia (MG)/non-MG from multiple treatments and timepoints in mouse, chick and zebrafish. **(B)** Schematics of data analysis and functional validation. For bulk RNA-Seq and ATAC-Seq data, we identified differentially expressed genes and differentially accessible regions. For scRNA-Seq, we performed cell clustering and MG trajectory to identify species-specific and evolutionarily conserved changes in gene expression. After integrative analysis, we reconstructed the regulatory network to identify key regulators for further validation. MGPCs, Müller glia-derived progenitor cells. **(C)** Number of single cells from scRNA-Seq. The number in brackets indicates the number of sequenced libraries. LD, light damage; GF, growth factor (insulin+FGF2); T+R, TNFα and RO4929097 treatment. **(D)** Distribution of the correlations for all replicates examined. The number in brackets of the legend indicates the number of replicates analyzed.

## Results

### Comprehensive profiling of MG from zebrafish, chick and mouse retinas

To identify gene regulatory networks that control MG reprogramming, we used three different well-characterized stimuli. These are light damage (outer retinal injury), NMDA treatment (inner retinal injury), and treatment with exogenous factors that induces MG reprogramming in an injury-independent manner. In zebrafish, injury-independent reprogramming was assessed using TNFα and a gamma-secretase inhibitor (T+R) (*12*). In chick, we tested the effects of FGF2 and insulin, which was previously shown to induce MG reprogramming independent of injury (*20*). In the mouse, we also tested FGF2 and insulin treatment, which induces limited MG proliferation, but not neurogenic competence (*21*).

For bulk RNA-Seq and ATAC-Seq, in zebrafish, we used the reporter line Tg[*gfap:EGFP*]nt11 (*22*) and cell sorting to enrich MG cells, purifying 9.8% of total input retinal cells (Fig. S1A). In mouse, we performed intraperitoneal injection of tamoxifen at postnatal day (P) 21 to induce MG-specific Sun1-GFP expression in *GlastCreERT2;CAG-lsl-Sun1-GFP* mice (*23*), purifying 3.5% of total input cells (Fig. S1A). Based on RT-qPCR, we had an overall enrichment of 30-fold of canonical MG marker genes such as *rlbp1a/Rlbp1* and *Glul* in the GFP-positive (GFP+) fraction in both species (Fig. S1B). Bulk RNA-Seq samples were generated from both the GFP+ MG and GFP-neuronal fractions in zebrafish and mouse, while ATAC-Seq samples were generated from the GFP+ cells. Each sample had a minimum of two biological replicates. In both species, we analyzed retinas with a time course following either inner retinal injury induced by NMDA excitotoxicity or outer retinal injury induced by light damage. In total, we generated 100 bulk RNA-Seq libraries and 40 bulk ATAC-Seq libraries (Fig. 1A, Table S1, Supplementary Data 1).

In parallel, we conducted scRNA-Seq analysis of whole retinas at the same time points as used for the bulk RNA-Seq. We profiled retinas following either NMDA or light damage in zebrafish and mouse, as well as following NMDA damage in P10 chick. To distinguish injury-responsive genes from injury-independent reprogramming genes, we profiled zebrafish retinas at multiple time points following T+R treatment, and a single time point in chick and mouse retinas following FGF2+insulin treatment. In addition, we profiled zebrafish retinas at multiple timepoints during development. In total, we obtained 287,790 single cells across all these conditions (Fig. 1C). We observed high concordance between experimental replicates for bulk RNA-Seq, scRNA-Seq and bulk ATAC-Seq (Fig. 1D, Fig. S1C-E).

### RNA-Seq analysis reveals species-specific MG response to retinal injury

We first compared bulk RNA-Seq data from GFP+ and GFP-samples to identify a core set of MG-enriched genes (Fig. S2A). Reads from both zebrafish and mouse showed high mappability (Fig. S2B), and hierarchical cluster analysis revealed that biological replicate samples were highly similar (Fig. S2C). Comparative analysis of GFP+ and GFP-samples confirmed MG enrichment of most MG-specific genes previously described in the literature (Fig. S2D, E, Supplementary Data 2). Many known markers specific to other retinal cell types were enriched in the GFP-samples.

Principal component analysis (PCA) of bulk RNA-Seq data from zebrafish and mouse revealed common species-specific patterns of transcriptional changes in MG in response to both NMDA and light damages (Fig. 2A, B). Both forms of injury induced rapid and similar changes in each species. In mouse, the most dramatic transcriptional changes occurred rapidly at early time points, within 3 and through 12 hrs following injury. At later time points following both NMDA and light damage, MG progressively reverted to a transcriptional state that closely resembles resting, uninjured MG (Fig. 2A). However, in zebrafish, both NMDA and light damage induced MG to adopt a transcriptional state that progressively diverged from the resting state (Fig. 2B). Extensive overlap in expression was observed between genes differentially regulated by light damage and NMDA treatment in MG at all time points in both zebrafish and mouse (Fig. S3A, B, Supplementary Data 3, 4).

**Fig. 2.**
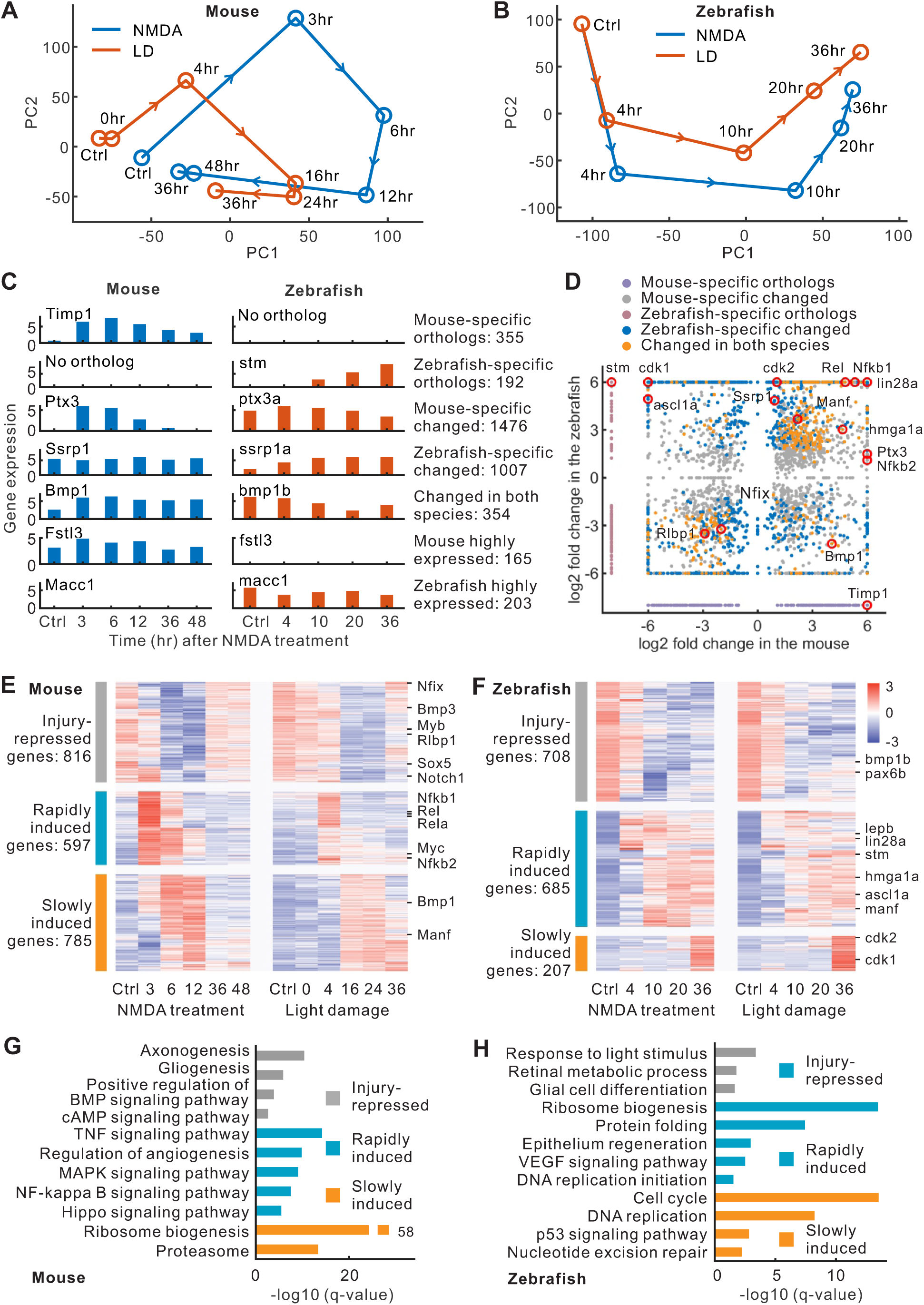
Dynamic bulk transcriptomes in mouse and zebrafish MG. **(A, B)** Principal component (PC) analysis of MG expression from retinal injury models in mouse and zebrafish. **(C)** Examples of genes showing species-specific changes or differences in baseline expression. **(D)** Comparison of correlated and differentially expressed genes (cDEG) between mouse and zebrafish. The maximal fold-change was used in treatment vs. control. **(E, F)** Heatmap of cDEGs in mouse and zebrafish. Scaled expression of GFP+ samples was shown in the color bar. **(G, H)** Most highly enriched functions of cDEGs in zebrafish and mouse.

We hypothesized that some genes specifically expressed in zebrafish might promote MGPC formation, while other genes specifically expressed in mouse might repress MGPC formation. We predicted that these genes would fall into four broad categories. The first category are injury-regulated genes that lack an ortholog in the other species. We identified 355 mouse-specific injury-regulated genes, and 192 zebrafish-specific genes in this category (Fig. 2C, D, Fig. S3C, Supplementary Data 5). The second category consists of genes for which orthologs are present in both species, but which show injury-regulated expression in only one species. These include 1476 genes in mouse and 1007 genes in zebrafish. The third category consists of 354 genes that have different patterns of injury-induced changes in the two species. The final category consists of genes that do not show injury-regulated changes in expression, but show large differences in baseline expression in the two species, and may thus serve to promote or inhibit competence to form MGPCs. We observed 165 mouse genes and 203 zebrafish genes in this final category.

These differentially expressed genes (DEGs) showed three broad patterns of injury-induced regulation in both zebrafish and mouse (Fig. 2E, F). A large subset of genes showed high and selective expression in resting MG, which were then rapidly repressed following injury. The two remaining categories showed rapid and slow injury-dependent induction, respectively. Gene ontology (GO) analysis of DEGs showed major species-specific differences in the functions of these genes. For instance, mouse MG genes that were rapidly induced after injury were enriched for the TNF, MAPK, Nfkb and Hippo signaling pathways (Fig. 2G). In contrast, in zebrafish MG, rapidly induced genes were enriched for ribosome biogenesis (Fig. 2H). Ribosome biogenesis was also enriched among slowly induced genes in mouse MG, while in zebrafish, slowly induced genes were enriched in cell cycle-related functions. These results are consistent with the fact that zebrafish MG begin to express *pcna* at 25 hrs, and undergo cell division at 36 hrs, following light damage (*24–26*).

### ScRNA-Seq analysis identifies resting and reprogrammed MG

We next performed scRNA-Seq analysis of zebrafish, chick and mouse retinas. We first analyzed the unique molecular identifiers (UMI) and genes in each sample. On average, we observed ∼1000-1500 genes and ∼2500-3250 UMI per cell. Some consistent differences were seen across species and treatment conditions. For instance, injured retinas expressed higher numbers of genes and UMI (Fig. S4A). T-stochastic nearest neighbor embedding (tSNE) analysis readily separated retinal cell types into distinct clusters (Fig. 3A-F, Supplementary Data 6), and no clear batch effects were observed among different samples (Fig. S4B, S5A, S5C, S5G, S6A, S6B).

**Fig. 3.**
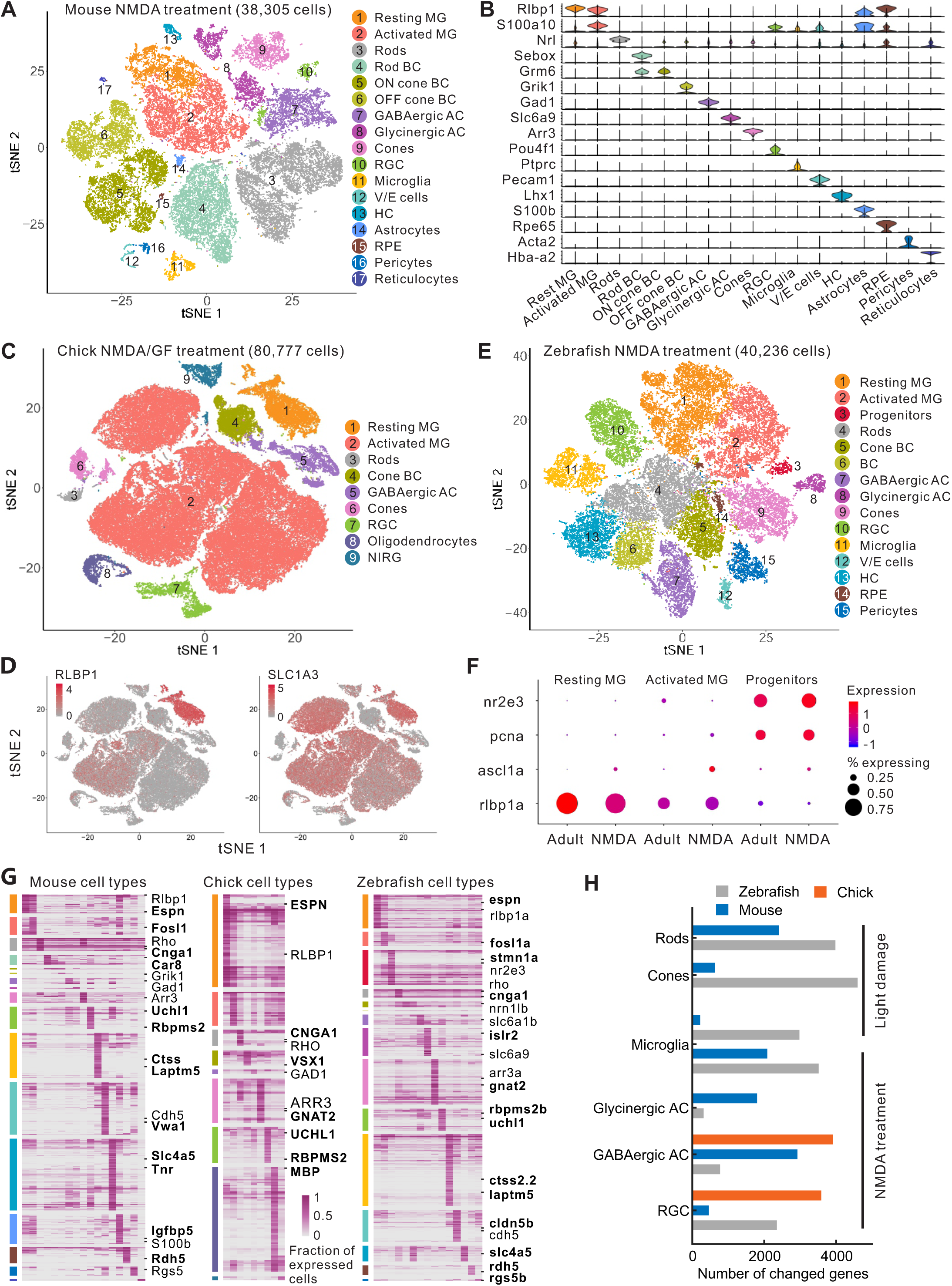
Single-cell transcriptomes from mouse, chick and zebrafish retinas. **(A)** tSNE view of mouse retinal cells after NMDA injury. **(B)** Violin plot showing the expression of known markers in mouse retinal cells. **(C)** tSNE view of chick retinal cells. **(D)** Single-cell expression of MG marker genes in the chick. **(E)** tSNE view of zebrafish retinal cells. **(F)** Gene expression of MG and rod progenitors in adult control and NMDA-treated samples from zebrafish. **(G)** Known and new (in bold) cell-type specific marker genes in all three species. Each column represents an individual cell type. **(H)** Number of pseudo-temporally changed genes in the retinal cells primarily targeted by each injury model in mouse, chick and zebrafish. MG, Müller glia cells; BC, bipolar cells; AC, amacrine cells; RGC, retinal ganglion cells; V/E cells, vascular/endothelial cells; HC, horizontal cells; RPE, retinal pigment epithelium; NIRG, non-astrocytic inner retinal glial cells.

Using well-established marker genes (Supplementary Data 7), along with MG-enriched genes identified in our bulk RNA-Seq analysis (Fig. S2D, S2E, Supplementary Data 2), we successfully annotated all major retinal cell types in each species (Fig. 3A-F, Fig. S4C, S5, S6A, S6B). While all major cell types were detected in each of the different treatments, the efficiency of capture of individual cell types varied depending on the injury models. For instance, more activated MG and microglia were captured following both injury and T+R treatment relative to control, while fewer photoreceptor cells were captured following light damage relative to NMDA and T+R treatment (Fig. S6C, S6D, S7A).

In mouse, we readily distinguished rod bipolars from cone ON and OFF bipolar cells, and GABAergic from glycinergic amacrine cells (Fig. 3A, B, Fig. S5G). Small subpopulations of non-neuroretinal cells, including microglia, vascular/endothelial cells, astrocytes, pericytes and RPE cells were also detected. In chick, we detected all major neuronal subtypes along with two glial cell types -- oligodendrocytes and non-astrocytic inner retinal glia (NIRG), which are found specifically in chick (*27, 28*) (Fig. 3C, Fig. S5B). In zebrafish, we observed a molecularly distinct subset of progenitor cells in the uninjured retina, which strongly expresses *nr2e3* (Fig. 3E, F, Fig. S5D), which likely correspond to rod progenitors derived from MG in the central retina during persistent neurogenesis (*29*).

In all three species, we readily distinguished resting and activated MG cells following each treatment. Many MG-enriched genes showed evolutionarily conserved patterns of expression. For instance, *Rlbp1/RLBP1/rlbp1a* all showed a high and selective expression in resting MG, and reduced expression following injury (Fig. 3B, D, F). Other genes such as *Gfap/GFAP/gfap* showed species-specific, injury-dependent expression patterns (Fig. S5F). In mouse, *Gfap* was selectively induced following injury, while it was expressed in both resting and activated MG in zebrafish, and barely detected in chick MG. In all three species, MG were over-represented relative to their actual abundance in the intact retina, representing 25.8%, 38.2%, and 16.7% of total retinal cells in zebrafish, chick and mouse, respectively (Fig. S6C, S6D, S7A). This likely reflects bias in cell survival and capture efficiency (*30, 31*). Of note, we did not observe any clear evidence for molecular heterogeneity of MG cells in the untreated control retina from any of the three species.

While cell type-specific marker genes have been identified for major retinal cell types in mammals, they are not as well annotated in zebrafish and chick. Comparison of cell type-specific transcriptomes allowed us to identify new evolutionarily conserved and species-specific marker genes for each of the common cell types in all three species (Fig. 3G, Supplementary Data 8, 9). For instance, we identified a new marker gene *Espn*/*ESPN*/*espn* that was selectively expressed in MG of all three species (Fig. 3G).

In parallel, we investigated injury-regulated genes in non-MG cells. In analyzing bulk RNA-Seq data, we identified several hundred genes which were changed in the MG-depleted fraction in zebrafish and mouse following both NMDA treatment and light damage (Supplementary Data 3). For the scRNA-Seq data, we focused our analysis only on cell types most directly affected by the injury: rod and cone photoreceptors in light damage, amacrine and ganglion cells in NMDA damage, and microglia in both models. We observed large numbers of pseudo-temporally changed genes (PCGs) in all cell types and identified some genes induced in multiple cell types (Fig. 3H, Fig. S7B-D, Supplementary Data 10). These include the cytokines *c7b* and *cxcl18b*, which were induced in both neurons and microglia in zebrafish, and the neuropeptide *Cartpt/cart3*, which was induced in neurons and microglia in both zebrafish and mice. These factors may serve as general signals of cellular injury, and may contribute to injury-mediated activation of MG.

### Trajectory analysis reveals dramatic differences in proliferative and neurogenic competence in MG across species

To profile dynamic injury-induced changes in gene expression in MG, we conducted trajectory analysis on resting and activated MG from our scRNA-Seq data. Consistent with bulk RNA-Seq data (Fig. 2A), mouse MG showed rapid injury-induced changes in gene expression after both light damage and NMDA treatment, followed by a gradual reversion to a transcriptional state that closely resembles resting MG (Fig. 4A, B, Fig. S8, Supplementary Data 11). In contrast, zebrafish MG showed progressive and consistent changes in gene expression following both light damage and NMDA injury (Fig. 4C, Fig. S9), again closely resembling what were seen using bulk RNA-Seq (Fig. 2B). In zebrafish, T+R treatment produced similar changes in gene expression (Fig. S9). In zebrafish, a small side branch was also observed in both damage models and T+R treated retinas, which may represent a transient reactive state (Fig. 4C, Fig. S9). This small side branch was enriched for genes associated with gliosis, such as *manf*, while the major branch showed increased expression of neurogenesis and proliferation-related genes, such as *pcna* (Fig. S9B, C). RNA velocity analysis (*32*) indicates that, following NMDA treatment, MG passed from a resting to a gliotic state, and from there either transitioned to a proliferative state or reverted to a resting state (Fig. 4C).

**Fig. 4.**
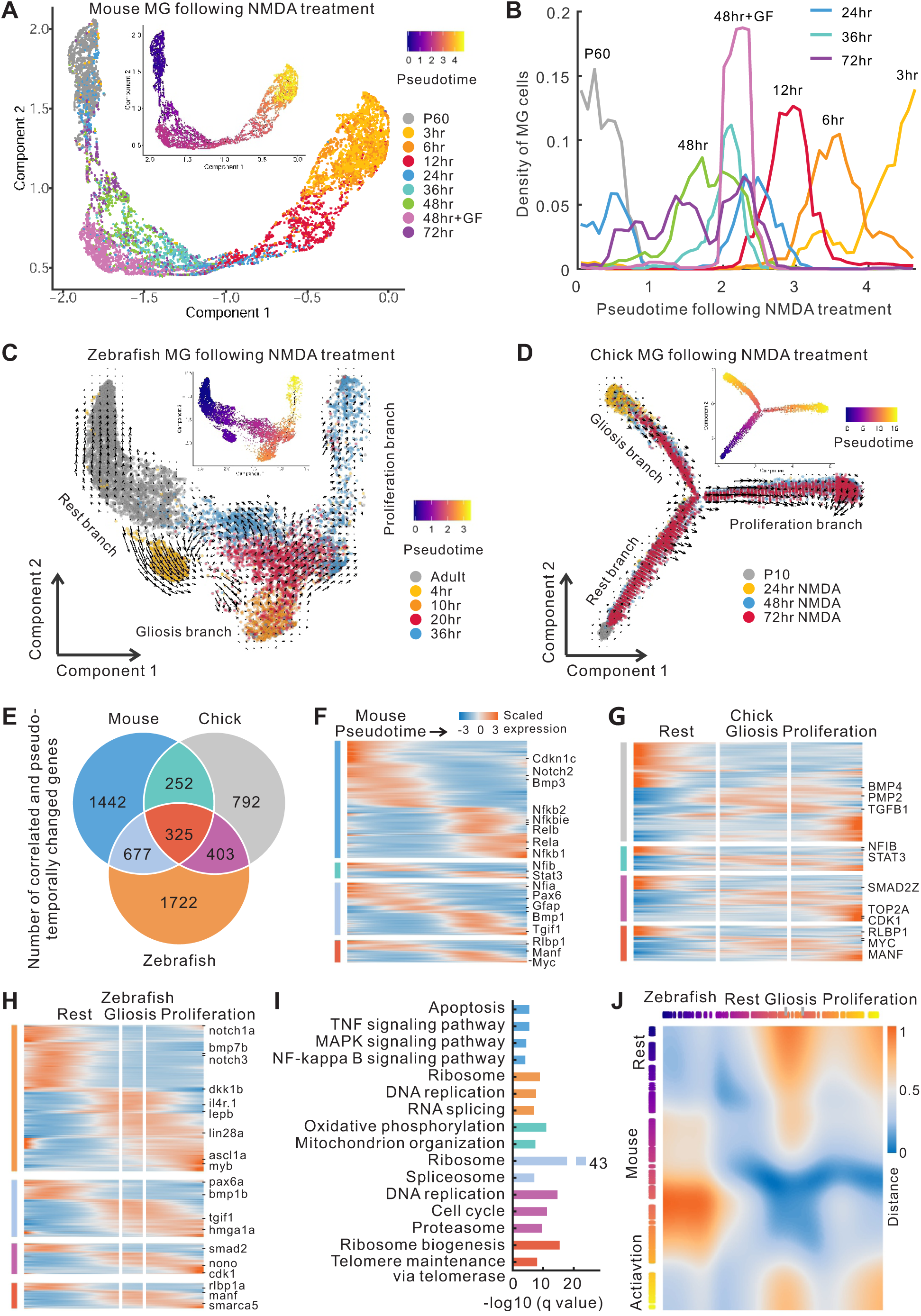
Trajectory and pseudotime analysis of mouse, chick and zebrafish MG. **(A)**Trajectory of mouse MG following NMDA treatment. Inserted plot shows the pseudotime of MG measured by scRNA-Seq. GF, growth factor (insulin+FGF2) treatment. **(B)** The density of mouse MG across the pseudotime. **(C, D)** Trajectory of zebrafish (**C**) and chick (**D**) MG. Arrows indicate the directions of MG transition identified through RNA velocity analysis. **(E)** Venn diagram of the number of correlated and pseudo-temporally changed genes (cPCGs) in three species. **(F-H)** Heatmap of cPCGs from three species. Color bars on the left correspond to the categories of cPCGs in Fig. 4E. **(I)** Top enriched functions of cPCGs. Colors represent the categories of species-specific or shared cPCGs corresponding to Fig. 4E. **(J)** Alignment of zebrafish and mouse MG trajectories. The trajectory is colored to indicate pseudotime. The color of the heatmap represents the cell dissimilarity (distance).

In contrast, chick MG exhibited a clearly branched trajectory (Fig. 4D, Fig. S10). One branch was enriched for cells profiled at 24 hours following NMDA treatment, while the second was enriched for cells from later time points (48 and 72 hours). Analysis of gene expression indicated that, while both branches show reduced expression of genes specific to resting MG such as *RLBP1,* the branch that predominantly contains cells from 24 hours was enriched in genes associated with reactive gliosis, such as *MANF* and *WNT6* (Fig. S10C-D, Supplementary Data 11). The other branch, in contrast, was enriched for genes associated with proliferation (*CDK1*) and neurogenesis (*ASCL1*) (*33*). MG from undamaged retinas treated with growth factors (FGF2 + insulin) occupied branches that were enriched for genes associated with reactive gliosis and proliferation, with very few having a transcriptional profile that resembled resting MG (Fig. 4D, Fig. S10A, E). RNA velocity analysis indicates that by 24hrs after injury, MG in the gliotic branch were transitioning back towards the branch point, and like in zebrafish, from that point either reprogrammed to a proliferative state or reverted to a resting state (Fig. 4D).

Collectively, these findings indicate that chick and zebrafish MG undergo reprogramming by passing through a gliotic state prior to generating MGPCs. To more comprehensively identify genes that are markers of the gliotic stage, we directly compared cells at this gliotic stage to resting MG in all three species in both injury and non-injury models, resulting in an inventory of both evolutionarily conserved and species-specific molecular markers of gliosis (Fig. S8H-I, S9H and S10F; Supplementary Data 12). Meanwhile, we also identified branch-specific genes in zebrafish and chick (Fig. S10G, Supplementary Data 13, 14).

By comparing PCGs detected using scRNA-Seq in different injury and treatment models, we next sought to identify core sets of evolutionarily conserved and species-specific patterns of gene expression in MG. We observed high overlap in both zebrafish and mouse between PCGs in light damage and NMDA injury (Fig. S11A, 11B, Supplementary Data 15, 16, 17). T+R treatment overlapped extensively with both injury models in zebrafish, as did NMDA and FGF2+insulin treatment in chick. We observed examples of evolutionarily conserved changes in gene expression, such as the decreased expression of *Rlbp1*/*RLBP1*/*rlbp1a* and the increased expression of *Manf*/*MANF*/*manf*, and species-specific changes such as selective induction of *Rela/b* and *Nfkb1/2* expression in mouse MG (Fig. 4E-H, Fig. S11C). Gene Ontology (GO) analysis likewise identified functional categories enriched in evolutionarily conserved PCGs, such as ribosome biogenesis, and in species-specific PCGs, such as DNA replication and cell cycle regulators, which were enriched in zebrafish and chick, but not in mice (Fig. 4I).

Finally, we calculated the dissimilarity of global gene expression profiles in MG at different stages following injury across species. We observed that resting MG from mouse and zebrafish were similar to each other, as were gliotic MG (Fig. 4J). In mice, we observed that a subset of gliotic MG resembled the proliferative branch in the zebrafish trajectory, which is consistent with observations that mouse MG transiently upregulate cell cycle regulators such as *Ccnd1* and *Cdk4* following injury (*5, 19*). In zebrafish, MG in the resting, gliosis and proliferative branches were close to chick MG from corresponding branches (Fig. S11D).

### MG reprogramming in zebrafish reverses the gene expression program associated with gliogenesis

To determine whether the formation of MGPCs from resting MG represented a simple reversal of gene expression changes observed during MG differentiation from multipotent retinal progenitor cells (RPCs), we next obtained scRNA-Seq data from developing zebrafish retinas, and integrated this with our analysis of injury-induced changes in gene expression in MG. Our recent scRNA-Seq study of developing mouse retina identified two transcriptionally and developmentally distinct RPC subtypes, which were designated primary and neurogenic RPCs (*30*). Using marker genes identified from this and other studies (Supplementary Data 7), we identified primary and neurogenic RPCs in the embryonic zebrafish retina (Fig. 5A, Fig. S12, Supplementary Data 18). We next aggregated all RPCs and MG cells, and performed trajectory analysis. While NMDA injury clearly induced many MG to shift to a state which closely resembled that of multipotent RPCs, we also observed a separate branch that was specific to injured MG (Fig. 5B, C). This injury-associated branch was most clearly observed in NMDA treatment (Fig. S13), but was also seen in light damage (Fig. S14) and T+R treatment (Fig. S15). This branch was primarily populated by cells profiled at early time points following injury or T+R treatment (Fig. S13B, S14E, S15E). This injury-associated branch was selectively enriched for gliosis-associated genes such as *manf* and *lepb*, as well as cytokine receptors such as *il4r.1*, while the main branch was enriched for genes that regulate proliferation and neurogenic competence, such as *ascl1a*, *hmgn2* and *cdk1* (Fig. S13C-F, S14F, S15F, Supplementary Data 19).

**Fig. 5.**
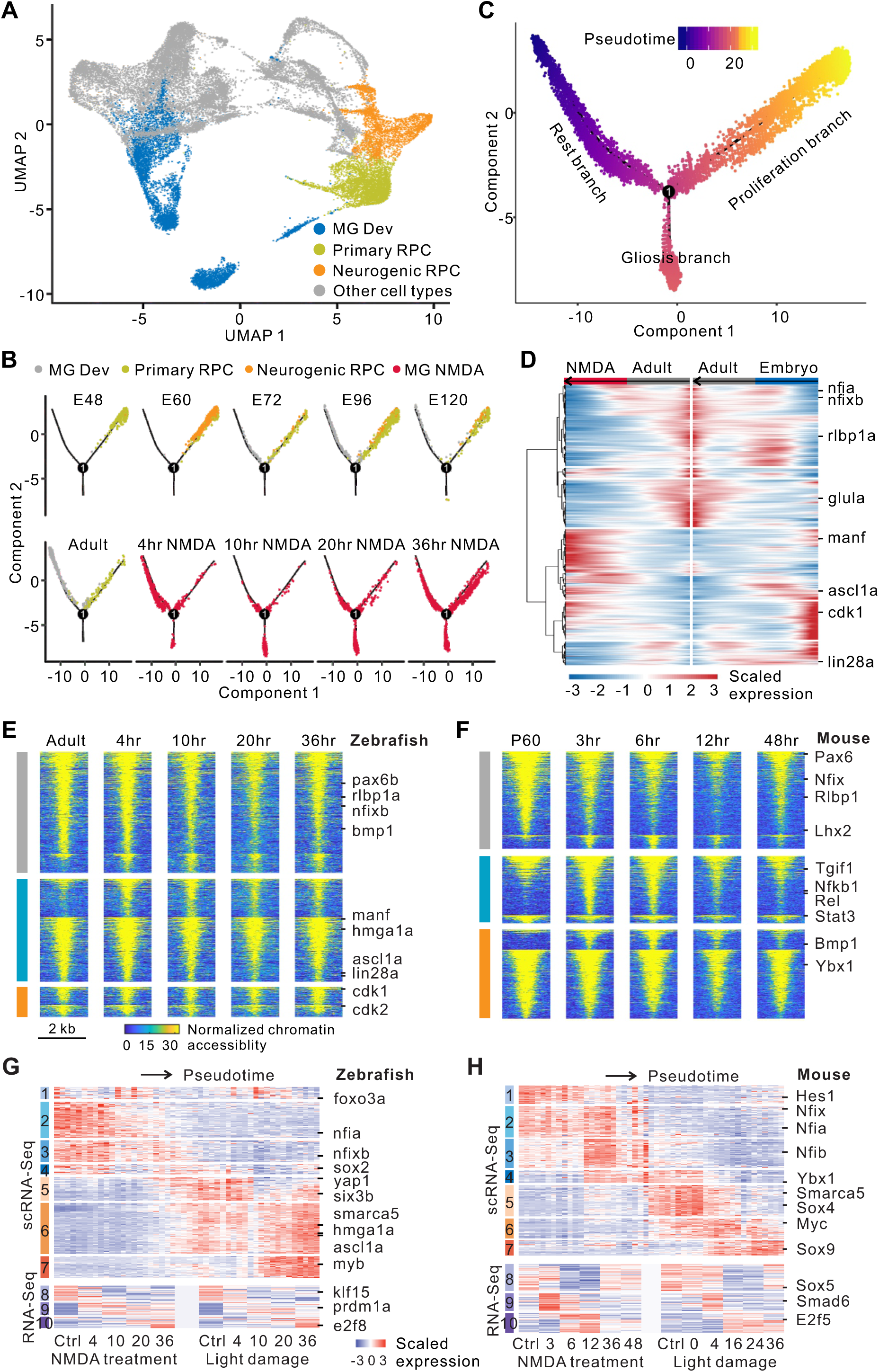
Analysis of developing zebrafish retinas and ATAC-Seq data. **(A)**UMAP view of retinal cell types from multiple timepoints (48 hrs to 120 hrs) during zebrafish development using scRNA-Seq. **(B)** Trajectory of RPC/MG from zebrafish development and NMDA treatment. **(C)** Pseudotime of zebrafish RPC/MG. **(D)** Heatmap of single-cell expression of pseudo-temporally changed genes shared by developing MG and NMDA-treated MG. Scaled expression profile for each gene is shown in the heatmap. The column represents single cells ordered by pseudotime. **(E, F)** Changes in chromatin accessibility of correlated differentially expressed genes (cDEGs) following NMDA treatment in zebrafish (**E**) and mouse (**F**). Each line indicates a correlated and differentially accessible region (cDAR) which is associated with injury-repressed (grey), rapidly induced (cyan) and slowly induced (orange) cDEGs from Fig. 2E, F. In each group, cDARs are separated by their positive and negative correlations with gene expression, and then ranked by the intensity of chromatin accessibility. **(G, H)** Ten modules are derived from differentially or highly expressed genes in zebrafish (**G**) and mouse (**H**). In each module (color bars on the left), example TFs are shown on the right.

Extensive overlap was observed between genes that showed dynamic expression during development and following NMDA treatment (Fig. S13A), light damage (Fig. S14A), and T+R treatment (Fig. S15A). For overlapping genes, we found that a great majority showed strongly negative correlation, with genes highly expressed early in development showing low expression in resting MG and vice versa, as expected (Fig. 5D). Taken together, these data suggest that, after first passing through a gliotic stage, MG undergo reprogramming by reversing the normal temporal pattern of gene expression associated with gliogenesis.

### Identification of gene regulatory networks that control MG states

We next analyzed ATAC-Seq data obtained from zebrafish and mouse MG. We observed an expected size distribution of ATAC-Seq inserts in NMDA-treated and light-damaged samples in both zebrafish and mouse (Fig. S16A). A comparison of differentially accessible chromatin regions (DARs) from NMDA and light-damaged samples revealed extensive overlap between these two injury conditions in both mouse and zebrafish (Fig. S16B, C, Supplementary Data 20-23). As expected, a majority of DARs are from candidate regulatory regions (Fig. S16D).

Overall, there was a strong correlation of chromatin accessibility and gene expression (Fig. S16E, F). We observed many species-specific and evolutionarily conserved dynamic injury-induced changes in chromatin accessibility. For instance, while *ascl1a* and *lin28a* were both devoid of open chromatin regions in unstimulated zebrafish and mouse MG, a rapid change in chromatin accessibility in response to injury was observed at the promoters of both genes in zebrafish but not in mouse (Fig. 5E, 5F, Fig. S17A). Likewise, we observed a rapid and transient change in the promoter accessibility of genes that were selectively expressed at early time points of retinal injury in mouse but not in zebrafish MG, such as *Nfkb1, Nfkb2, Rel* and *Stat3* (Fig. 5E, 5F, Fig. S17A).

We next developed a computational method to construct an integrated model of the gene regulatory networks that control response to injury in MG (Fig. S17B). We term this algorithm Integrated Regulatory Network Analysis (**IReNA**). To this end, we integrated bulk and single-cell RNA-Seq data from all the damage/treatment models in both zebrafish and mouse to obtain a comprehensive list of DEGs/PCGs at each time point, and to identify transcription factors (TFs) expressed in MG (Supplementary Data 24). In parallel, we analyzed ATAC-Seq data to identify differential footprints for MG-expressed TFs, and their target motifs associated with DEGs (*34*). We then inferred regulatory relationships between these TFs and their target genes, and used K-means clustering to separate target genes into different regulatory modules (see Methods). Based on these inferred regulatory relationships, we then constructed an integrated gene regulatory network (GRN) for each module, and compared these GRNs between zebrafish and mouse.

Integrating bulk and single-cell RNA-Seq data, we clustered 4190 and 3942 injury-regulated genes in zebrafish and mouse, respectively, into 10 gene modules each (Fig. 5G, H, Fig. S17C, D, Supplementary Data 25). Based on differential footprinting analysis, we identified 26,083 and 30,177 regulatory relationships among 192 and 212 MG-expressed TFs in zebrafish and mouse, respectively (Supplementary Data 26). We then calculated the coefficient correlation between the expression of TFs and their target genes to infer whether individual TFs activate (a positive correlation) or repress (a negative correlation) expression of their target genes. From 192 zebrafish and 212 mouse MG-expressed TFs, we separately identified 97 and 156 TFs that were predicted to activate or repress the overrepresented genes in any one of the 10 gene modules (Fig. 5G, H, Supplementary Data 27). We further constructed two regulatory networks consisting of these TFs in zebrafish and mouse (Fig. S18, S19, Supplementary Data 28). We observed that some TFs regulated target genes in the same modules, such as the regulations of *nfia* in zebrafish and *Sox9* in mouse (Fig. S18, S19, S20A). Other TFs primarily target genes from different modules. For instance, in zebrafish, *myb* from the proliferation-related module activates the module associated with neurogenesis (Fig. S18, S20A). In mouse, *Nfib* represses genes in a separate module related to gliosis (Fig. S19, S20A).

To obtain insight into regulatory relationships between modules, we calculated the number of inferred positive and negative regulatory relationships between each pair of modules, and determined whether these were statistically overrepresented relative to the whole network. An intramodular regulatory network was then generated for both zebrafish and mouse (Fig. 6A, B). In zebrafish, we observed multiple TF modules that were most active in resting MG (Fig. 6A, Fig. S18). TFs in these resting modules (module 1, 2, 3, 4 and 8) are predicted to both positively regulate expression of TFs in their own module, and activate expression of TFs in other modules active in resting MG. This quiescence-associated network is in turn predicted to repress expression of TFs found in a set of modules associated with gliosis (Fig. 6A), which also includes a number of genes associated with neurogenic competence. TFs found in this second group (module 5, 6 and 9) include *ascl1a*, *tgif1* and *six3b* -- all of which have been known to be necessary for generating MGPCs in zebrafish (*15, 35*). These TFs in the gliosis network in turn positively induce their own expression, and negatively regulate TFs in the quiescence-associated network. Directly downstream of the gliosis network lies a third set of TF modules (module 7 and 10), which regulate genes associated with DNA replication and cell cycle progression. These TF modules include both TFs known to regulate cell cycle entry, such as *myca* and *e2f3,* and TFs that promote neurogenesis, such as *olig2* and *foxn4* (Fig. 6A, Fig. S18) (*36–38*). In zebrafish MG, we conclude that a bistable regulatory relationship between mutually opposed quiescence and gliosis networks controls the transition from a resting to an activated state. Once MG have transitioned to a gliotic state, this then rapidly leads to either a return to quiescence or the acquisition of neurogenic and proliferative competence, and full conversion to MGPCs (Fig. 6C).

**Fig. 6.**
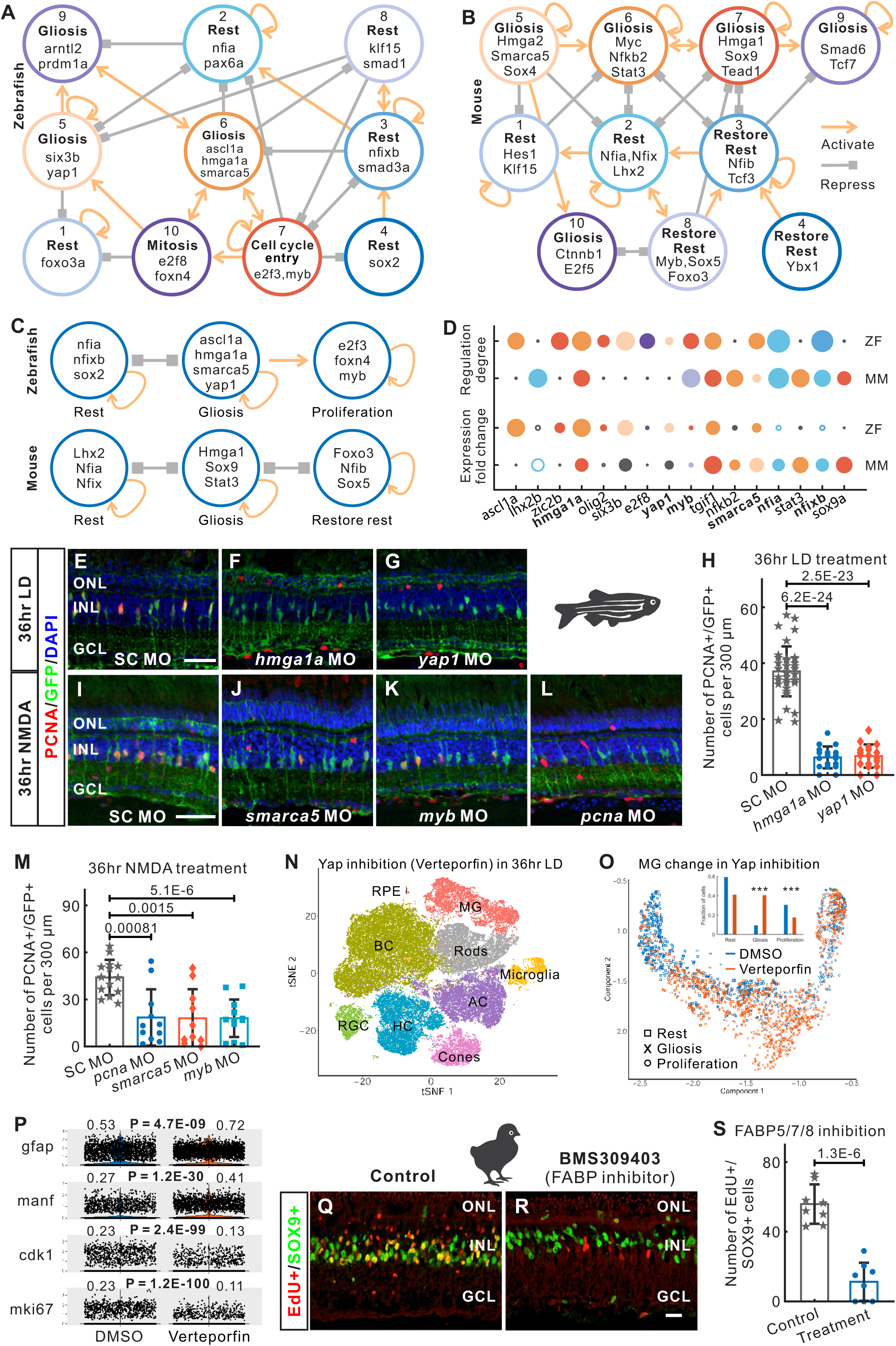
MG regulatory network and functional validation of key regulators in zebrafish and chick. **(A, B)** Intramodular regulatory network in zebrafish (**A**) and mouse (**B**). Colored circles represent the modules. The connection indicates a significant regulation among modules. **(C)** Super regulatory network in zebrafish and mouse. **(D)** Comparison of TF features between zebrafish (ZF) and mouse (MM). TFs are ordered by the difference of expression fold changes between zebrafish and mouse. Colors represent TF modules in **A**, **B**. Black circle indicates that the TF is not enriched in corresponding species. Circle size indicates fold change of gene expression, or the number of TF regulations. Hollow circles indicate decreased expression in treatment vs. control. **(E-H)** In zebrafish, morpholino-mediated knockdown of *hmga1a* and *yap1* inhibited MG proliferation after 36hr light damage. **(I-M)** Knockdown of *smarca5* and *myb* significantly decreased MG proliferation after NMDA damage, as compared with the control morpholino. *Pcna* morpholinos were used as a positive control. **(N)** t-SNE view of retinal cells from control (DMSO) and Yap inhibition (verteporfin) in 36hr light damage. **(O)** Distribution of DMSO- and verteporfin-treated MG in the trajectory. Inserted figure shows the fraction of DMSO- and verteporfin-treated MG in three branches. Fisher’s exact test was used. **(P)** Single-cell expression of gliosis-related and proliferation-related genes in DMSO- and verteporfin-treated MG. The fraction of expressed cells and p-value (Fisher’s exact test) are shown. **(Q-S)** In chick, pharmacological inhibition of FABP5/7/8 using BMS309403 significantly inhibited MGPC formation 3 days after NMDA injury. ONL, outer nuclear layer; INL, inner nuclear layer; GCL, ganglion cell layer. Scale bar, 20 μm. Error bar in bar-plot represents the standard deviation.

In mouse, we likewise observed three interconnected networks of TF modules (Fig. 6B, C). Like in zebrafish, the first network consists of TFs in cross-activating modules in resting MG (module 1 and 2) (Fig. 6B, Fig. S19). TFs in this network include *Lhx2*, which has been known to negatively regulate reactive gliosis in mouse MG (*39*), and Notch pathway genes such as *Hes1*. This quiescence-associated network is predicted to repress genes in a separate gliosis-associated network (module 5, 6, 7 and 9). These in turn are predicted to repress the quiescence-associated network, establishing a bistable, cross-repressive regulatory relationship like that seen in the zebrafish (Fig. 6A-C). TFs in the gliosis-associated network included Nfkb pathway components (*Rela, Relb, Nfkb1, Nfkb2*), and multiple TFs that were previously shown to promote glial activation (*Stat3*, *Sox9*) (*40, 41*), along with the Hippo pathway target *Tead1* (Fig. 6B).

Interestingly, TFs associated with cell cycle entry (*Myc*) and even neurogenesis (*Sox4*) (*42*), are also included in this gliosis-associated network in the mouse. Expression of these genes, however, peaks relatively early following injury and drops rapidly, as eventually do all gliosis-associated genes (Fig. 5H). An indication as to how this reversion to a resting state occurs in mouse comes from the presence of a third network of TF modules (module 3, 4 and 8) that is not found in the zebrafish (Fig. 6B). This network is also enriched for TFs expressed in resting MG, e.g., *Nfib and Sox5*, which showed reduced expression immediately following injury, but elevated expression at later time points (Fig. 5H). These TF modules are separated from those that are predicted to maintain MG in a resting state, and appear to mediate restoration of quiescence following activation (Fig. 6B). Taken together, this implies that the transition between rest, gliosis, and restoration of the resting state in the mouse are mediated by three separate bistable, auto-activating transcriptional regulatory networks (Fig. 6C). Furthermore, they imply that MG transition through a gliotic state in order to reprogram into MGPCs, and that dedicated transcriptional regulatory networks exist to maintain and restore a resting state in mouse MG.

### Functional validation of candidate genes that regulate proliferative and neurogenic competence in MG

We next tested whether loss of function of key TFs in the regulatory network affects MGPC formation. To select such TFs, we integrated the regulatory networks and gene expression datasets, and compared them across species (Fig. 6D, Supplementary Data 29). We examined the degree of connectivity of each TF within the regulatory networks. This value, which is defined as the number of edges in the regulatory networks, reflects the relative importance of the TF in that network (Fig. S18, S19, S20B). Similarly, we also assessed the fold-change of gene expression during the course of activation and reprogramming, and integrated these results to select top candidates.

We then directly tested whether individual TFs that were predicted to promote gliosis were required for MG reprogramming and proliferation. The non-histone HMG family DNA binding protein gene *hmga1a/Hmga1* was induced in reactive MG in zebrafish and mouse (Fig. 6A-D, Fig. S21A). We used morpholinos to knockdown the expression of *hmga1a* to determine whether it was necessary for MGPC formation in zebrafish. Retinas electroporated with the *hmga1a* morpholino exhibited a significant reduction in the number of proliferating MG 36 hours following light damage, relative to retinas electroporated with the Standard Control morphant (Fig. 6E, F, H). Even after 72 hours of light damage, the *hmga1a* morphant retina possessed significantly fewer PCNA-positive MGPCs than the Standard Control morphant retina (Fig. S21B-D). ScRNA-Seq analysis of *hmga1a* morphant retinas showed decreased expression in a larger number of markers specific to the proliferative state relative to Standard Control morphants, and a corresponding increase in markers specific to the resting and gliotic stages (Fig. S21I-M, Supplementary Data 30).

We next examined the potential role of *yap1*, which was also induced in the gliotic stage (Fig. 6A, C, Fig. S21A). YAP was recently shown to be required for MG to exit quiescence in both *Xenopus* and mice (*5, 19*). We found that morpholino-mediated knockdown of Yap1 protein in the light-damaged zebrafish retina significantly reduced the number of proliferating MG relative to the Standard Control morphant (Fig. 6E, G, H). Furthermore, injection of the drug verteporfin, which inhibits the interaction of Yap1 and TEAD in the nucleus (*43*), nearly eliminated PCNA-positive MG in the light-damaged retina relative to the DMSO control in a dose-dependent manner (Fig. S21E-H). ScRNA-Seq analysis of verteporfin-treated retinas indicated a substantial reduction in the fraction of proliferative MG, and a corresponding increase in the fraction of gliotic MG (Fig. 6N-P, Fig. S21N-P). We also tested the role of *smarca5*, a SWI/SNF-related matrix-associated actin-dependent regulator of chromatin, that was also expressed in gliotic MG in the zebrafish and mouse (Fig. 6A-D, Fig. S21A). We used morpholinos to knockdown the expression of Smarca5 and found that the *smarca5* morpholino likewise significantly reduced the number of proliferating MG relative to the Standard Control morphants after NMDA injury (Fig. 6I, J, M).

To test the potential role of proliferation-related genes, we used morpholinos to knockdown the expression of Myb, a transcription factor that was predominantly expressed in proliferating zebrafish cells (Fig. 6A, C, Fig. S20A). Knockdown of Myb significantly reduced the number of proliferating MG relative to the Standard Control morphant in the NMDA-damaged retina (Fig. 6I, K, M). For each TF targeted in zebrafish, MG proliferation was reduced to a comparable extent to that seen with morphants targeting proliferating cell nuclear antigen (*pcna*) gene, which served as a positive control (Fig. 6E-M).

Although a core set of evolutionarily conserved genes were upregulated in reactive gliosis (Supplementary Data 12), many genes showed species-specific induction during gliosis. In chick but not in zebrafish or mouse, for instance, we identified the lipid binding proteins *FABP5*, *FABP7* and *FABP8* (also known as *PMP2*) as highly expressed in gliotic MG (Fig. S22A). The expression of FABP8 protein was further confirmed by immunohistochemistry (Fig. S22B, C). To globally inhibit the function of these *FABP* family genes, we treated chick retina with the FABP5/7/8 inhibitor BMS309403 (*44*) and measured NMDA-induced formation of MGPCs. We observed a significant decrease in the number of MGPCs compared with the DMSO-injected control retina (Fig. 6Q-S).

Finally, to test the functional role of TFs predicted to maintain and/or restore MG quiescence, we selected the NFI family transcription factors *Nfia*, *Nfib* and *Nfix*, which were expressed in resting MG, down-regulated rapidly after injury but elevated at later timepoints in mouse (Fig. S22D). To this end, we examined the phenotype of tamoxifen-fed *GlastCreER;CAG-lsl-Sun1-GFP;Nfia/b/x^lox/lox^* mice, in which NFI family members were selectively deleted from mature MG (Fig. S22E, F). ScRNA-Seq analysis indicates that *Nfia/b/x*-deficient MG were clearly distinct from the control (Fig. S23A, B). *Nfia/b/x* loss of function, as expected, led to a reduction in expression of genes selectively expressed in resting MG, such as *Glul*, *Rlbp1, Aqp4* and *Apoe*. *Nfia/b/x*- deficient MG also upregulated cell cycle regulators such as *Ccnd1* and *Ccnd3*, along with the neurogenic bHLH factor *Ascl1* (Fig. S23C).

We next tested whether NMDA or a combination of NMDA and growth factors (FGF2 and insulin) could induce *Nfia/b/x*-deficient MG to proliferate and generate neurons. We found that *Nfia/b/x*-deficient MG proliferated at substantially higher levels than wildtype MG in response to NMDA treatment, both with and without growth factors. EdU+/GFP+ cells were found in both inner and outer nuclear layers 3 days after injury (Fig. 7A, Fig. S23D, E). Furthermore, a substantial number of the MG-derived GFP+ cells expressed either the photoreceptor and bipolar cell marker Crx and amacrine/ganglion/horizontal cell markers HuC/D and NeuN at day 14 after treatment (Fig. 7B, Fig. S23F, G).

**Fig. 7.**
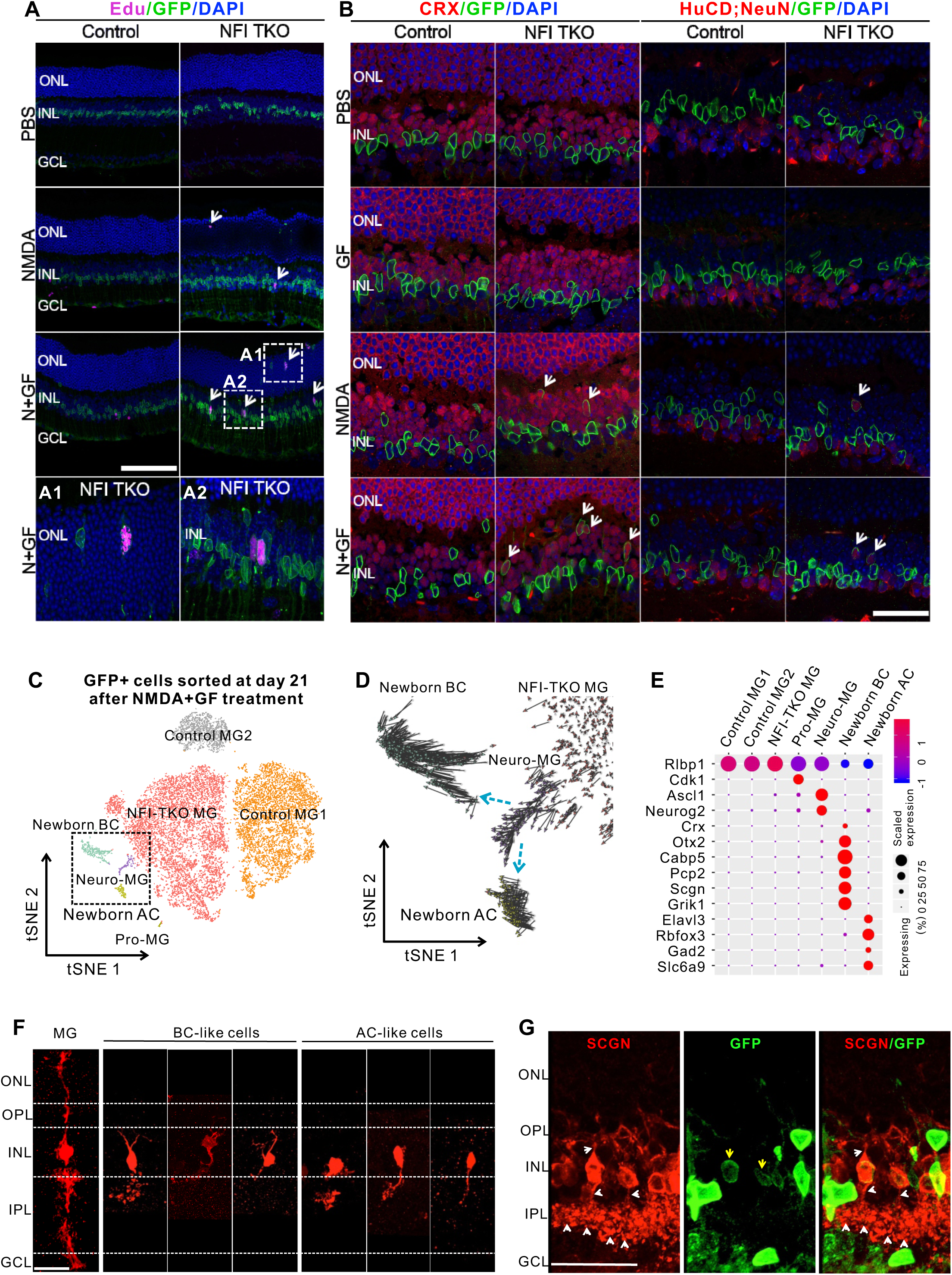
Loss of *Nfia/b/x* enables MG to proliferate and generate neurons following injury in mouse. **(A)**Adult control and MG-specific *Nfia/b/x* knockout mice were intravitreally injected with either PBS, NMDA, or NMDA with growth factor (N+GF). Treatment of NMDA with and without growth factor significantly increased MG proliferation in both ONL and INL layers (**A1, A2**) in the MG-specific *Nfia/b/x* knockout retina as compared with the control at day 3 after injection. **(B)** Retinal samples were collected at day 14 after injection and immunostained for retinal neuronal markers. *Nfia/b/x-*deficient MG generated CRX+ neurons (white arrows) in response to NMDA injury. Combination of NMDA and GF treatment significantly increased the number of MG-derived CRX+ neurons in the MG-specific *Nfia/b/x* knockout retina. NMDA injury led to the generation of HuCD+NeuN+ neurons (white arrows) from MG cells in the MG-specific *Nfia/b/x* knockout retina. GF treatment further enhanced NMDA-induced generation of HuCD+NeuN+ neurons. **(C)** tSNE-view of GFP+ cells flow-sorted from control and MG-specific *Nfia/b/x* knockout retina at day 21 after N+GF treatment show different types of MG-derived cells, including neurogenic MG (Neuro-MG), proliferative MG (Pro-MG), newborn bipolar cells (BCs) and newborn amacrine cells (ACs). **(D)** RNA velocity analysis indicates that neuro-MG give rise to newborn BC and AC cells. RNA velocity was shown for cells in the dotted box from **C**. Primary arrows are shown in blue dotted arrows. **(E)** Single-cell expression of known maker genes, including a MG-specific marker *Rlbp1*, neurogenic marker *Ascl1*, proliferative marker *Cdk1*, bipolar cell markers and amacrine cell markers. (**F**) Morphological characterizations of MG-derived neurons in *NFIa/b/x*-deficient retinas using AAV9-pCAG-Flex-TdTomato. Adult mice were intravitreally injected with AAVs and treated with Tamoxifen diet for 3 weeks. Neuronal morphology was examined at Day 14 after N+GF treatment. GFP/TdTomato+ cells in *NFIa/b/x*-deficient retinas showed morphological features resembling bipolar cells and amacrine cells. (**G**) Characterization of MG-derived bipolar cells from *NFIa/b/x*-deficient retinas at 2 months after N+GF treatment using Secretagonin (SCGN) protein marker. The newborn bipolar cells send processes to the OPL and IPL retinal layers (white arrows). MG, Muller glia; BC, bipolar cell; AC, amacrine cell; ONL, outer nuclear layer; OPL, outer plexiform layer; INL, inner nuclear layer; IPL, inner plexiform layer; GCL, ganglion cell layer; N+GF, NMDA, FGF2 and insulin. Scale bar, 20 ì m. Error bar in bar-plot represents the standard deviation.

To more fully characterize the identity of neurons generated from *Nfia/b/x*-deficient MG, we next performed scRNA-Seq on GFP-positive MG and MG-derived cells FACS-sorted from *Nfia/b/x*-deficient MG and control retina 21 days after treatment with NMDA and growth factors (Fig. S23H, I). Our analysis showed that *Nfia/b/x*-deficient MG produced several distinct cell populations, including proliferative MG, neurogenic MG, newborn bipolar cells and newborn amacrine cells, after treatment with NMDA and growth factors (Fig. 7C). RNA velocity analysis indicated that neurogenic MG directly gave rise to newborn bipolar and amacrine cells (Fig. 7D). These newborn neurons expressed markers of multiple bipolar and amacrine subtypes (Fig.7E, Fig. S23J, Supplementary Data 31). Interestingly, the neurogenic MG did not express cell cycle regulators, indicating that these newborn neurons may have arisen from *Nfia/b/x*-deficient MG through a process of direct transdifferentiation. This was confirmed by immunohistochemical data which showed that newborn neurons rarely co-labeled with EdU (data not shown). Conversely, proliferative MG from *Nfia/b/x*-deficient MG did not express neurogenic markers, implying that they represent a population of symmetrically dividing, self-renewing MG. By comparison with native retinal cell types, we found that newborn bipolar and amacrine cells derived from *Nfia/b/x*-deficient MG closely resembled native bipolar and amacrine cells (Fig. S23K). Morphological analysis using AAV9-Flex-TdTomato showed that subsets of MG-derived cells resembled bipolar and amacrine cells, 14 days after injury (Fig. 7F). These MG-derived bipolar cells extend processes to both the OPL and IPL retinal layers (Fig. 7G). More complete characterizations of these neurons await further study. Taken together, these data clearly demonstrate that separate populations of bipolar and amacrine cells were generated from the *Nfia/b/x*-deficient MG, and that NFI factors act to repress proliferative and neurogenic competence in mouse.

## Discussion

This study is the first effort to comprehensively profile changes in gene expression and chromatin accessibility that underlie cellular reprogramming in species with dramatically different regenerative abilities. Retinal MG of zebrafish have a high capacity to proliferate and give rise to new neurons, while those of post-hatch chick retain high proliferative but limited neurogenic competence. Mouse MG, in contrast, lack substantial proliferative competence and do not generate neurons. Using both bulk and single-cell RNA-Seq, along with ATAC-Seq analysis, we have identified both evolutionarily conserved and species-specific transcriptomic and epigenomic events that occur during MG development, following outer and inner retinal injury, as well as following growth factor treatments that induce MG reprogramming in zebrafish and chick. Integration of these datasets has allowed identification of gene regulatory networks that are strong candidates for both promoting and restricting neurogenic and proliferative competence in MG. These datasets will serve as a starting point for future studies that functionally characterize these candidate genes.

This study identifies several key events that appear to be critical for regulating MG reprogramming. First, we observed that MG in all three species pass through a state resembling reactive gliosis. Reactive gliosis occurs following all forms of retinal injury, but differs considerably in its duration. It is a transient state in zebrafish, and a much more prolonged state in chick. In contrast, mouse MG arrest in gliosis, before returning to quiescence. Most reactive zebrafish and chick MG pass through this gliotic state to become MGPCs, which express high levels of neurogenic bHLH genes *Ascl1* in zebrafish, and much lower levels in chick, in line with their more limited neurogenic competence. Reactive gliosis is a critical step in the formation of MGPCs, as loss of function of genes that are highly enriched in reactive MG such as *hmga1a* in zebrafish and *FABP* family genes in chick blocks the formation of MGPCs. Previous studies in zebrafish have likewise identified multiple genes that show enriched expression in reactive MG -- such as *six3b*, *sox2*, *lepb* and others (*25, 35, 45*) -- to also be essential for MGPC formation.

Second, in both mouse and zebrafish, we identified dedicated gene regulatory networks that maintain or restore expression of genes specific to quiescence of MG. Notably, although reactive mouse MG transiently express cell cycle regulatory genes such as *Ccnd1, Ccnd2* and *Myc*, as well as the neurogenic *Sox4*, they return to quiescence rapidly, and have dedicated a transcriptional regulatory network that mediates this process. NFI family genes, which are members of this network, act to maintain expression of genes specific to quiescent MG and to restrict proliferative and neurogenic competence in mouse MG. NFI family genes show evolutionarily conserved expression patterns, with injury repressing expression in MG in all three species examined. Furthermore, selective deletion of NFI genes in mouse MG, in combination with NMDA treatment, resulted in both MG proliferation and formation of MG-derived neurons that closely resemble retinal bipolar and amacrine cells. Taken together, this suggests that NFI family genes act to maintain MG quiescence, and prevent expression of genes associated with transition to gliosis and/or MGPC status, and suppress neurogenesis from MGPCs. Other data support this proposed role for NFI factors. Our group has recently shown that *Nfia/b/x* act in late-stage retinal progenitors to drive gliogenesis and restrict neurogenic and proliferative competence, with *Nfia/b/x* loss of function mutants leading to persistence of proliferative progenitors and generation of excess rod photoreceptors (*30*). Furthermore, the observation that *Ascl1* overexpression induces direct transdifferentiation of MG to inner retinal neurons but not photoreceptors may result from the strong persistent expression of *Nfia/b/x* that is seen in these transformed cells (*17, 46*).

Ever since the development of the Yamanaka protocol for generation of iPSCs (*47*), the great majority of work aimed at induced dedifferentiation or cellular reprogramming has focused on gain of function approaches, in which TFs associated with the desired fate are overexpressed. However, with the exception of the iPSC generation, the effectiveness of these approaches has been limited, in large part because genes specific to the cell of origin are not efficiently repressed. This study highlights the importance of downregulating these gene regulatory networks in the context of endogenous cellular reprogramming, and identifies potential targets for improving regenerative therapies that rely on directed reprogramming of endogenous cells.

## Materials and Methods

### Animals

#### Fish maintenance

All zebrafish (*Danio rerio*) lines, including AB, *albino*; Tg[*gfap:GFP]mi2001* (*48*) and *albino*; Tg[*gfap*:EGFP]*nt11* (*22*) were maintained in the Center for Zebrafish Research at the University of Notre Dame Freimann Life Science Center. The adult zebrafish used in these studies were 6 to 12 months old (3-5cm in length). Fish were maintained under a light and dark cycle of 14 hours light and 10 hours of dark at 28.5°C. All experimental protocols were approved by the animal use committee at the University of Notre Dame and are in compliance with the ARVO statement for the use of animals in vision research.

#### Chick maintenance

The use of animals in these experiments was in accordance with the guidelines established by the National Institutes of Health and the Ohio State University. Newly hatched wild type leghorn chickens (*Gallus gallus domesticus*) were obtained from Meyer Hatchery (Polk, Ohio). Postnatal chicks were kept on a cycle of 12 hours light and 12 hours dark (lights on at 8:00 AM). Chicks were housed in a stainless steel brooder at about 25°C and received water and Purina^tm^ chick starter *ad libitum*.

Fertilized eggs were obtained from the Michigan State, Department of Animal Science Eggs were incubated between 36.6°C and 37.8°C and embryos were staged according to guidelines set by Hamburger and Hamilton (*49*).

#### Mice maintenance

CD1 mice were purchased from Charles River Laboratories (Wilmington, MA). *GLASTCreERT2* and *Sun1-sGFP* transgenic mice were provided by Jeremy Nathans (The Johns Hopkins University School of Medicine). *GLASTCreERT2*; *Sun1-sGFP* were generated by breeding and subsequent backcrossing. To induce *Cre* recombination, *GLASTCreERT2*; *Sun1-sGFP* mice at ∼2 weeks of age were intraperitoneally injected with Tamoxifen in corn oil for 2 consecutive days at 1mg/dose. *Nfia^lox/lox^*; *Nfib^lox/lox^*, and *Nfix^lox/lox^* mice were crossed to *GLASTCreERT2*; *Sun1-sGFP* mice. To generate MG-specific loss of function mutants of *Nfia/b/x* genes, 3 week-old *GLASTCreERT2*; *Nfia/bx^lox/lox^*;*Sun1-sGFP* mice were fed with tamoxifen diet for 3 weeks, followed by 2 weeks with normal diet. All mice were housed in a climate-controlled pathogen free facility on a 14 h-10 h light/dark cycle (07:00 lights on – 19:00 lights off). All experimental procedures were pre-approved by the Institutional Animal Care and Use Committee (IACUC) of the Johns Hopkins University School of Medicine.

#### Retinal damage paradigms in fish

To induce death of rods and cones, adult *albino*; Tg[*gfap*:EGFP]*nt11* fish were dark adapted for 14 days, then transferred to clear polycarbonate tanks placed between four fluorescent bulbs (20,000 lux) and water temperature maintained at 32°C for up to 72 hours (*50, 51*). Fish were euthanized by anesthetic overdose of 0.2% 2-phenoxyethanol in system water.

To induce amacrine and ganglion cell death, we injected N-methyl-D-aspartic acid (NMDA) (Sigma-Aldrich; St. Louis, MO) intravitreally (*52*). Adult Tg[*gfap:GFP]mi2001* zebrafish were anesthetized in 0.1% 2-phenoxyethanol. A sapphire blade was used to make an incision in the posterior cornea and 0.5 μl of 100 mM NMDA was injected into the intravitreal space of the eye using a 33 gauge Hamilton syringe. The fish were revived and placed in a 32°C dark incubator for up to 36 hours. Fish were euthanized by anesthetic overdose of 0.2% 2-phenoxyethanol in system water.

#### TNFα and RO4929097-induced proliferation

Recombinant TNFα was expressed in *E. coli* and purified as previously described (*53*). TNFα was diluted to a concentration of 1.5 mg/ml in water prior to injection. Undamaged adult Tg[*gfap:GFP]mi2001* fish were anesthetized in 0.1% 2-phenoxyethanol, cut in the posterior cornea with a sapphire blade, and intravitreally injected with 0.5 μl of TNFα using a 33 gauge Hamilton syringe every 12 hours. The fish were also intraperitoneally injected with 50 μl 0.75 mM RO4929097 (Selleckchem; Houston, TX) using a 30 gauge bevelled needle every 12 hours. The fish were revived and placed in a 32°C dark incubator for up to 72 hours before being euthanized by anesthetic overdose of 0.2% 2-phenoxyethanol in system water.

#### Intraocular injections in chicks

Chickens were anesthetized via inhalation of 2.5% isoflurane in oxygen and intraocular injections performed as described previously (*54*). For all experiments, the right eyes of chicks were injected with the “test” compound and the contra-lateral left eyes were injected with vehicle as a control. Compounds were injected in 20 μl sterile saline with 0.05 mg/ml bovine serum albumin added as a carrier. Compounds used in these studies included NMDA (38.5 or 154 g/dose; Sigma-Aldrich), FGF2 (200 ng/dose; R&D systems), insulin (1μg/dose; Sigma-Aldrich), and BMS309430 (500 ng/dose; Tocris). Two μg EdU was injected to label proliferating cells. Injection paradigms are included in each figure.

#### Mouse N-methyl-D-aspartate (NMDA) and light damage

For NMDA damage, adult mice, either CD1 or *GLASTCreERT2*; *Sun1-sGFP* mice at ∼2 months of age, were anesthetized with isoflurane inhalation. A puncture was made just behind the limbus with a 30G needle. Two microliters of 100mM NMDA in PBS was intravitreally injected using a syringe with a 33G blunt-ended needle. Mice were sacrificed, and retinas were collected at the indicated time points.

For NMDA and growth factor treatment of NFI TKO mice, adult *GLASTCreERT2*; *Nfia/bx^lox/lox^*;*Sun1-sGFP* and *GLASTCreERT2*;*Sun1-sGFP* control mice, both in CD1 background, were intravitreally injected with 2 μl of 100mM NMDA or 2 μl of 100mM NMDA with 100ng/μl FGF2 and 1μg/μl Insulin (N+GF treatment), followed by 2 μl of 500ng/μl EdU at 24 and 48hr after the first injection. For the N+GF treatment group, 2 μl of 100ng/μl FGF2 and 1μg/μl Insulin was co-injected with each EdU injection.

Contralateral eyes were injected with PBS and EdU as the controls. Retina were collected at the indicated timepoints for immunohistochemistry and fluorescence-activated cell sorting (FACS).

For morphological examination of MG and MG-derived neurons, *GLASTCreERT2*; *Nfia/bx^lox/lox^*;*Sun1-sGFP* and *GLASTCreERT2*;*Sun1-sGFP* control mice were intravitreally injected with 1 μl of AAV9-pCAG-Flex-Tdtomato (Addgene #28306) at 1×10¹³ vg/mL and treated with Tamoxifen diet for 3 weeks. Morphology of GFP/TdTomato+ MG and MG-derived neurons was examined at day 14 after N+GF treatment.

Light damage was performed as previously described (*40*). In brief, the mice were reared in cyclic 12-hour low light/12-hour dark conditions at the University of Florida animal housing facility. Prior to light damage, mice were placed in a modified cage equipped with dimmable white light LED strips. Light intensity was measured using a light meter (Thermo Fisher Scientific Inc., Waltham, MA) and set to 2000 lux. Animals were subjected to the damaging light for 4 hours (6PM-10PM) and were moved back to low light conditions to recover. All animals were kept in ventilated racks for the duration of the experiment and the lighting equipment was approved by Animal Care Services.

#### Zebrafish retinal dissociation, cell sorting and methanol fixation

Zebrafish were euthanized in 0.2% 2-phenoxyethanol and retinas dissected and placed in Leibowitz medium (Thermo Fisher Scientific) prior to dissociation. Retinas were subsequently placed in 20U/ml papain (10 retinas per 1ml) (Worthington), and incubated at 28°C for 30 minutes with gentle agitation. Cells were then pelletized and resuspended in PBS containing 0.1mg/ml leupeptin (Sigma-Aldrich) and 10U/ml DNaseI (Roche). Cells were then filtered through a 70 μm filter (Miltenyi Biotec), and kept on ice until sorting and incubated with 5 μg/ml propidium iodide (PI; Life Technologies).

Cells were transferred into a 5ml FACS tube and loaded into a FACS AriaII. Gating was set using dissociated retinas from *albino* zebrafish both unstained and PI stained and unstained Tg[*gfap*:EGFP]*nt11* zebrafish. GFP+ and GFP-cells were sorted directly into Trizol LS (Life Technologies), and immediately flash frozen.

For single-cell RNA-Seq dissociated cells were methanol fixed (*55*). Dissociated cells were pelleted and resuspended in D-PBS (Thermo Fisher Scientific). This was repeated for a total of 3 washes, with samples kept on ice throughout. Cells were then resuspended in 100 μl D-PBS in cryovials and 900 μl-20°C methanol added dropwise while slowly vortexing. Cells were left on ice for 15 minutes and then placed directly in −80°C.

#### Mouse retinal cell dissociation and fluorescence-activated cell sorting (FACS)

Mice were euthanized, and eye globes were removed and kept in ice-cold 1x PBS. The neural retinas were dissected, and cells were dissociated using Papain Dissociation System (LK003150, Worthington). Briefly, retinas were incubated in pre-equilibrated papain/DnaseI mix (20 units/ml papain and 100 units/ml DnaseI, 200ul per retina) in a 37^0^C water bath. Retinas were mixed by inverting the tubes every 5 min and gently triturated after the first 10 min at every 5 min with 1ml pipette tips. After 25 min of enzyme incubation, undigested retinal tissues were removed by running through a 50um cell filter. Dissociated cell mixtures were treated with DnaseI and subjected to density gradient centrifugation to remove the cell debris by following the manufacturer’s instructions. For FACS sorting experiments, dissociated cell pellets were resuspended in ice-cold PBS containing 2% heat-inactivated FBS and 5 units/ml DnaseI).

For scRNA-Seq experiments, cell pellets were rinsed twice without disturbing the cell pellet to completely remove any trace of DnaseI. Dissociated cells were then re-suspended with ice-cold PBS containing 0.04% BSA and 0.5U/μll RNase inhibitor to obtain a final cell concentration of 0.5E10+6 ∼ 1.5E10+6 cells/ml. Dissociated cells were filtered through a 50um filter, and cell count and viability were assessed by Trypan blue staining.

FACS experiments were performed using Sony SH800S Cell Sorter. Retinal cells from non-transgenic mice were used to set the gating threshold for GFP-positive cells. Cells were flow-sorted into GFP-positive and GFP-negative fractions into ice-cold PBS containing a final concentration of 10% heat-activated FBS. For ATAC-Seq, cells were kept on ice until use. For bulk RNA-Seq, cells were centrifuged at 500xg for 5min, resuspended in 700μl QIAzol lysis reagent (miRNAeasy Mini Kit) and stored at −80°C until RNA extraction.

#### Immunohistochemistry and imaging in mice

Mouse eye globes were collected and fixed in 4% PFA in PBS for 2hr at room temperature. Eyes were washed with PBS, and retina were carefully dissected and put in 30% sucrose overnight at 4°C. Retina were then embedded, cryosectioned at 20 um thickness and stored at −80°C. Sections were dried at 37°C for 20 min and washed 3 x 5 min with PBS prior to incubating with a blocking buffer (0.2% Triton, 5% horse serum in PBS) for 2 hr at room temperature. For CRX, HuC/D and NeuN immunostaining, retinal sections were treated with 100 mM sodium citrate pH 6.0 at −80°C for 30 min for antigen retrieval before the blocking step. Sections were then incubated with primary antibodies at indicated concentrations in the blocking buffer (Table S2) overnight at 4°C. Sections were washed 3x 10min in PBST (0.1% Triton in PBS) and incubated with secondary antibodies in the blocking buffer for 2hr at room temperature. Sections were counterstained with DAPI, washed 3 x 10 min in PBST and mounted in Vectashield with DAPI. EdU staining was performed by following the manufacturer instructions. Images were acquired by using confocal Zeiss LSM 700. EdU+/GFP+ cells were counted and averaged from > 6 random whole sections for each retina. CRX+/GFP+ and HuCD;NeuN+/GFP+ cells were counted from 4 images taken across > 4 random sections per retina. Fraction of CRX+/GFP+ and HuCD;NeuN+/GFP+ cells were calculated from total GFP+ cells. Each data point in the quantification bar graphs was calculated from individual retina. For morphological characterization of neurons, 10-20 z-stacked confocal images (0.15-0.35 μm per image) were taken for GFP/TdTomato+ cells and GFP/SCNG+ cells at day 14 and 2 months after N+GF treatment.

#### Injection and electroporation of morpholinos in zebrafish

Morpholino mediated knockdown was performed as previously described (*56, 57*). Morpholinos (Gene Tools) were reconstituted in 100 μl nuclease free water to yield a 3 mM solution. Dark-adapted *albino*; Tg[*gfap*:EGFP]*nt11* fish were anesthetized in 0.1% 2-phenoxyethanol and corneas were removed from the eye using tweezers, 0.5 μl of morpholino solution was intravitreally injected into the eye. Platinum plate electrode tweezers (Protech International Inc.) were used to deliver two 50 ms pulses (75 V with a 1 s pause between pulses) to the eye using a CUY21 Square Wave Electroporator (Protech International Inc.). Fish were then allowed to recover, and then either anaesthetized and intravitreally injected with NMDA and placed in a 32°C dark incubator for 36 hours or they were placed in constant intense light for either 36 or 72 hours. After either the NMDA- or light-damage, fish were euthanized by anesthetic overdose of 0.2% 2-phenoxyethanol in system water. The following previously validated morpholinos were used in this study: Standard Control: 5’-CCTCTTACCTCAGTTACAATTTATA-3’ (Gene Tools),

*pcna*: 5’-TGAACCAGACGTGCCTCAAACATTG-3’ (*56*),

*myb:* 5’-GCCGCCTCGCCATCCCGCTGTTCG-3’ (*58*),

*smarca5*: 5’-CTTCTTCCCGCTGCTGCTCCATGCT-3’ (*59*),

*hmga1a*: 5’-CTGTGTCCTTGCCAGAATCACTCAT-3’ (*60*),

*yap1*: 5’-CTCTTCTTTCTATCCAACAGAAACC-3’ (*61*).

#### Verteporfin injection in zebrafish

Verteporfin was resuspended at 3.48 mM in 100% DMSO and then diluted with water to the desired working concentration (either 12.5 or 25 μM). Dark-adapted *albino*; Tg[*gfap*:EGFP]*nt11* fish were anesthetized in 0.1% 2-phenoxyethanol and 0.5 μl of either verteporfin or DMSO was intravitreally injected into the eye and the fish were exposed to constant light damage. Every 12 hours, the intravitreal injections were repeated until 36 hours, when the fish were euthanized and retinal sections were analyzed by immunohistochemistry and confocal microscopy.

#### Immunohistochemistry and imaging in zebrafish

Immunohistochemistry was performed as described previously (*12, 53*). Fish were euthanized in 0.2% 2-phenoxyethanol. Eyes were collected and fixed in 9:1 ethanolic formaldehyde. After fixation, eyes were rehydrated through an ethanol gradient (90%, 80%, 70%, 50% v/v ethanol in water), then washed for 15 minutes in 5% sucrose in PBS, and incubated overnight in 30% sucrose in PBS at 4°C. Eyes were then transferred into a 2:1 mixture of tissue freezing media (TFM) (Triangle Biomedical Sciences; Durham, NC) and 30% sucrose in PBS and incubated at 4°C overnight. Eyes were embedded in 100% TFM and frozen at −80°C. 14 μm cryosections were prepared and stored at −80°C.

Sections were dried at 50°C for 20 minutes and the sections were surrounded using a liquid PAP pen (Ted Pella; Redding, CA). Sections were then rehydrated in PBS for 20 minutes prior to blocking for 1 hour (1% Tween-20, 1% Triton-X, 2% normal goat serum, 2% DMSO in PBS). Primary antibodies (Table S2) were diluted in blocking solution and slides incubated in primary antibody solution at room temperature overnight. Sections were washed 3 x 10 min in PBST (1x PBS with 0.05% Tween-20), and then incubated for 1 hour at room temperature with secondary antibodies in blocking solution at 1:1000, and counterstained with DAPI at 1:1000. Sections were washed 3 x 10 min in PBST, and No 1.5 coverslips (VWR) were mounted using Prolong Gold (Life Technologies). Confocal imaging was performed using a Nikon A1R laser scanning confocal microscope. Images were obtained from the center of the dorsal retina using only sections containing the optic stalk. Quantification was performed using 6.5 μm z-stacks, and cell counts normalized to 300 μm. Data was statistically analyzed using one-way ANOVA with Tukey’s post-hoc test if more than two groups were compared, otherwise using Student’s t-test with unequal variances.

#### Immunohistochemistry and imaging in chick

Tissues were fixed, sectioned, and immunolabeled as described previously (*11, 62*). Working dilutions and sources of antibodies were listed in Table S2. None of the observed labeling was due to non-specific labeling of secondary antibodies or autofluorescence because sections labeled with secondary antibodies alone were devoid of fluorescence. Secondary antibodies were diluted to 1:1000 in PBS plus 0.2% Triton X-100. Draq5 (10 uM; Thermofisher) was added to the secondary antibody solution to label nuclei.

For EdU-labeling, immunolabeled sections were fixed in 4% formaldehyde in PBS for 5 minutes at room temperature, washed for 5 minutes with PBS, permeabilized with 0.5% Triton X-100 in PBS for 1 minute at room temperature, and washed twice for 5 minutes in PBS. Sections were incubated for 30 minutes at room temperature in 2M Tris, 50 mM CuSO_4_, Alexa Fluor 568 Azide (Thermo Fisher Scientific), and 0.5M ascorbic acid in dH_2_O.

Digital photomicrographs were obtained using Leica DM5000B microscopes equipped with epifluorescence and Leica DC500 digital camera. Confocal images were obtained using a Leica SP8 imaging. Images were optimized for color, brightness and contrast, multiple channels overlaid and figures constructed by using Adobe Photoshop. Cell counts were performed on representative images. To avoid the possibility of region-specific differences within the retina, cell counts were consistently made from the same region of the retina for each data set.

#### RNA extraction and RT-qPCR

RNA was extracted from both GFP-positive and GFP-negative cell fractions using miRNAeasy Mini Kit (#217004, Qiagen). To test for MG enrichment of sorted samples, expression of some MG markers was measured by RT-qPCR. Briefly, RNA samples were first reverse-transcribed into cDNA using random primers and Superscript IV reverse transcriptase (#18091050, ThermoFisher) according to the manufacturer’s instructions. The qPCR assays were performed on the cDNA using GoTaq Green Master Mix (#M7122, Promega) using a StepOnePlus Real-time instrument (ThermoFisher). Intron-spanning primers were designed to specifically quantify targeted mRNA transcripts. Glyceraldehyde 3-phosphate dehydrogenase (*Gapdh*) expression was used as the endogenous control (Table S3). PCR specificity was monitored by determining the product melting temperature and by checking for the presence of a single DNA band on agarose gel analysis of the qRT-PCR products

#### Bulk RNA-Seq

Flow-sorted RNA samples were sent to the Deep Sequencing and Microarray Core (Johns Hopkins University) for library preparation and sequencing. Briefly, ribosomal RNA was depleted, and total RNA was captured from the RNA samples using Illumina TruSeq Stranded RNA LT kit Ribo-Zero^TM^ Gold (# 15032619, Illumina). Around 8-10 libraries were pooled and sequenced for paired-end 75 cycles using the NextSeq 500 system with ∼400-500 million reads per run, resulting in ∼45-55 million reads per library.

#### ScRNA-Seq

For fresh samples (NMDA mouse damage, chick, zebrafish development and some zebrafish damaged samples), cells were resuspended with ice-cold PBS containing 0.04% BSA and 0.5U/UL RNase inhibitor. For methanol-fixed samples (mouse LD damage and most of zebrafish damage samples), fixed cells were processed by following 10x genomic protocol for methanol-fixed cells. Briefly, methanol-fixed cells were centrifuged at 3000xg for 10 min at 4^0^C and washed 3 times in an iced-cold re-suspension buffer containing PBS, 1% BSA and 0.5U/UL RNAse inhibitors. Dissociated cells were resuspended in resuspension buffer and filtered through a 50ul cell filter. Cells (∼10k) were loaded into a 10x Genomics Chromium Single Cell system (10x Genomics, CA, USA) using v2 chemistry following the manufacturer’s instructions. Libraries were pooled and sequenced on Illumina NextSeq with ∼200 million reads per library. Sequencing results were processed through the Cell Ranger 2.1.1 pipeline (10x Genomics) with *--expect cell* =10,000 parameter.

#### ATAC-Seq

Sorted cells (∼50k-75k) from GFP-positive samples were used for ATAC library preparation. In our experiences, there are two main factors that are critical for a successful ATAC-Seq library preparation. First, cell nuclei need to be intact during nuclei isolation process by optimizing IGEPAL concentration and minimal pipetting. Second, enzyme concentration and tagmentation time need to be optimized depending on cell types. Nuclear isolation and all centrifugation steps were carried out on ice in a 4^0^C cold room. Briefly, cells were spun down at 500xg for 5 min. Cells were rinsed with ice-cold PBS without disturbing cell pellet and centrifuged again at 500xg for 5 min. Cells were lysed by adding 50UL of ice-cold cell lysis buffer (10 mM Tris.Cl pH 7.4, 10 mM NaCl, 3 mM MgCl_2_) containing 0.03% IGEPAL and protease inhibitors (1 tablet per 7ml of lysis buffer) and mixing 3 times by pipetting. Cells were then immediately spun down at 500xg for 10 min and washed with 150UL of ice-cold lysis buffer without IGEPAL and protease inhibitors. For tagmentation, cell nuclei were incubated with 2.5 UL enzyme in 50UL total volume at 37^0^C in a thermocycler (Illumina Nextera DNA library prep kit, #FC1211030). DNA was cleaned up using MinElute PCR purification kit (#28006, Qiagen) and eluted in 10UL of EB buffer.

ATAC-Seq DNA was amplified, and the number of PCR cycles were calculated by following previously described protocol (*63*). PCR products (10UL) were run on a 1.5% agarose gel for expected DNA pattern. PCR products were then cleaned up by double-sized selection by using Ampure beads (Agencout AMPure XP, Beckman Coulter, #A63880) to remove large and small DNA fragments. This was performed by using 1:0.5 and 1:1.6 ratio of sample to Ampure beads (v/v). Completed ATAC-Seq libraries (8–10) were then analyzed by Fragment Bioanalyzer and sequenced for paired-end 75 cycles using the NextSeq 500 system with ∼400-500 million reads per run.

#### Bulk RNA-Seq data analysis

Using NextSeq 500 with a paired-end read of 100bp (RNA-Seq), we measured gene expression profiles of 50 GFP+ (GFP-positive) and 50 GFP- (GFP-negative) samples (Table S1, Supplementary Data 1). Raw data from mouse and zebrafish were separately mapped to the GRCm38/mm10 and GRCz10/danRer10 genome assembly using STAR (*64*). Raw counts of genes were further used to calculate FPKM (Fragments Per Kilobase Of Exon Per Million) and identify differentially expressed genes through EdgeR (*65*).

To identify GFP+ (Müller glial cells, MG) or GFP- (non-Müller glial cells, non-MG) specific genes, we performed t-test of paired GFP+/GFP- samples from NMDA treatment and light damage models. Genes with fold change > 2 and FDR (false discovery rate) < 0.01 were chosen as GFP+ or GFP-specific expressed genes (Supplementary Data 2). Furthermore, we identified species-highly expressed genes from GFP+ specific expressed genes. In each species and each GFP+ sample, we calculated the rank of gene expression. According to the rank of gene expression, we performed Wilcox rank test between two species to identify species-enriched genes in GFP+ samples.

We compared each time point of treated samples with control samples to identify temporally changed genes in GFP+ samples. If fold change > 1.5 and FDR < 0.05, the gene was defined as significantly differentially expressed across different time points of treatment (Supplementary Data 3). To identify model-independent response genes that are consistently changed between NMDA treatment and light damage model, we calculated the Pearson’s correlation coefficient between models in each species. Meanwhile, we shuffled the expression for each gene and then calculated the correlation as the control. The correlated differentially expressed genes (cDEGs) are defined as the genes that are significantly differentially expressed in both models and the correlation of expression patterns between the models > 0.5 (Supplementary Data 4).

To compare differentially or highly expressed genes between two species, we identified orthologs between mouse and zebrafish based on vertebrate ortholog information from MGI database (http://www.informatics.jax.org/). We then grouped genes into six categories: (i) mouse gene mapping to zero ortholog in zebrafish (1 vs. 0), (ii) zebrafish gene mapping to zero ortholog in mouse (0 vs. 1), (iii) one mouse/zebrafish gene mapping to one ortholog in the other species (1 vs.1), (iv) one mouse gene mapping to multiple zebrafish orthologs (1 vs. N), (v) multiple mouse genes mapping to one zebrafish ortholog (N vs. 1), (vi) multiple mouse/zebrafish genes mapping to multiple orthologs in the other species (N vs. N). If multiple orthologs were mapped, we chose the most significantly differentially expressed genes. If there is no significantly differential expression, we select the most correlated and expressed genes between two models. To identify genes that could potentially promote retinal regeneration from bulk RNA-seq data, we divided differentially or highly expressed genes into seven categories, including mouse-specific changed genes without orthologs in the zebrafish, zebrafish-specific changed genes without orthologs in the mouse, mouse-specific changed genes with orthologs, zebrafish-specific changed genes with orthologs, genes changed in species but with different expression patterns, mouse highly expressed genes, and zebrafish highly expressed genes (Supplementary Data 5).

In addition, we performed K-means of GFP+ samples to separate cDEGs into three groups, including injury-repressed genes, rapidly induced genes and slowly induced genes. Function enrichment analysis on genes was conducted in R package ‘clusterProfiler’ (*66*). Hierarchical clustering of all RNA-Seq samples, and principal component analysis (PCA) were also processed in R platform.

#### Mapping and quality control of single-cell RNA-Seq data

We analyzed 19, 12, and 27 single-cell RNA-Seq libraries from mouse, chick, and zebrafish, respectively (scRNA-Seq, Supplementary Data 1). Mouse, chick and zebrafish raw reads from scRNA-Seq were separately mapped to GRCm38/mm10, Gallus_gallus-5.0/galGal5 and GRCz10/danRer10 genome using STAR (*67*). The gene-cell matrices were generated using Cell Ranger from 10X Genomics. We further calculated the number of genes and the number of UMIs (Unique molecular identifiers) for each cell. In the chick, we removed low-quality cells which have < 200 genes or < 500 UMIs. In mouse and zebrafish, cells with > 200 genes and > 1000 UMIs were included for further analysis. Meanwhile, for each sample we also removed the outlier cells which have > 2-fold of average number of genes, or > 2-fold of average number of UMIs across all cells in each library.

#### Single-cell clustering

For all single cells from each treatment, we first identified variable genes which have 0.1∼8 expression level and > 1 dispersion. Based on the variable genes, we performed multiple alignment and clustering of single cells across different time points of treatment using Seurat (*68*). Single-cell clusters were identified through k-nearest neighbors and a shared nearest neighbor modularity optimization. The single-cell clustering was visualized through t-SNE (*69*). In analysis, top 5000 variable genes were used in mouse NMDA and light damage treatment, and zebrafish T+R treatment. In the zebrafish NMDA and light damage, we used top 8000 and 6000 variable genes. In the chick, top 4000 variable genes were used. No obvious batch effects were observed after performing multiple alignments of single cells.

#### Identification of cell doublets in single-cell RNA sequencing

In order to identify cell doublets in our scRNA-Seq data, we developed a new algorithm. First, we performed single-cell clustering and identified marker genes for each cluster (i.e. cluster-specific genes). For each cluster marker, we then used AUC (area under the receiver operating characteristic curve of gene expression) to calculate the gene power (*GP_i, j_*) (*68*), which indicates the capability of marker *j* to distinguish cluster *i* and the other clusters. For each cell and each cluster, we next calculated summed powers (*SP*):

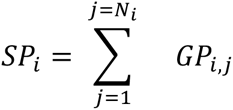

where *N_i_* is the number of markers in cluster *i*. Meanwhile, we calculated the threshold of *SP* in each cluster indicating whether the cell expresses markers in that cluster. The threshold was obtained by performing statistics on *SP* of all cells for each cluster.

Based on *SP* of the cell and threshold, we determined cell doublets. If a cell simultaneously expresses markers from more than one cluster, the cell is regarded as a doublet. In each scRNA-Seq clustering, we identified 5∼10% cell doublets. Cell doublets were removed for further analysis.

#### Identification of cell types and markers for each cluster

We used known specific marker genes to annotate cell type for each cluster of cells (Supplementary Data 7). Most known marker genes were reported in the mouse. For the chick and zebrafish, we identified orthologs of known mouse marker genes, and used those markers to determine cell types of clusters. Meanwhile, we identified novel markers of each cluster (or cell type) in each injury model and species by setting > 0.3 gene power (Supplementary Data 8). By comparing these marker genes across species, we identified species-specific or conserved markers for each cell type (Supplementary Data 9).

#### Trajectory and pseudo-time analysis of single cells

We constructed the trajectory of MG in response to retinal injury and performed pseudo-time analysis using Monocle (*70*). Variable genes were used to construct the trajectory of MG. Pseudo-time of cells was calculated by setting unstimulated control cells as the root. Furthermore, we identified significantly temporally changed genes along pseudotime (PCGs), which were defined as differentially expressed as a function of pseudo-time (Supplementary Data 15). We used the following criteria to identify PCGs: fraction of expressed cells > 0.01, single-cell expression difference > 0.1 and q-value < 0.001. Single-cell expression difference was calculated:

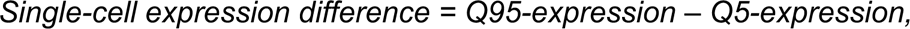

where Q95-expression and Q5-expression separately represent 0.95 and 0.05 quantile of log-transformed single-cell expression values across all bins of pseudo-time. We grouped cells across pseudo-time into 50 bins with equal pseudo-time intervals. Average single-cell expression profiles were obtained by averaging all cells within each bin (bin-derived expression).

To identify model-independent PCGs, we also used bin-derived expression to calculate the single-cell expression correlation between two treatment models. In each species, PCGs were further selected according to single-cell expression correlation between models. The cutoff of single-cell expression correlation for analysis was set to 0.2 based on the distribution of the correlation of shuffled single-cell expression (Supplementary Data 16, 17).

In the trajectories of chick and zebrafish MG, we found three obvious branches. Using monocle, we used single-cell expression as a function of branch state to identify branch-specific expressed genes (BEGs). BEGs were defined as the difference of proportion of expressed cells between one branch and the other two branches > 0.1 and q-value < 0.001 (Supplementary Data 13, 14).

We aligned MG trajectories from two species through cellAlign (*71*). In alignment, we separated MG cells based on branch state and then sorted MG using pseudo-time. Correlated PCGs shared by two species were used in trajectory alignment.

#### RNA velocity analysis

To minimize batch effect, we chose branch-specific or cell type-specific genes for performing RNA velocity (*32*). In zebrafish NDMA treatment, we selected top 50 specifically expressed genes for each branch according to the differential proportion of expressed cells. Among the 150 genes, we selected 92 genes that satisfy the criteria, the number of cells detected spliced UMIs (S) > 10, and the number of cells detected unspliced UMIs (U) is 1 < U < 300. Other parameters were set to the default.

In chick, top 60 specifically expressed genes were selected for each branch. The criteria for further selection are S > 10 and 1 < U < 400. In total, 107 genes were used for RNA velocity in chick. For flow-sorted scRNA-Seq samples at 21 days after NFI TKO and NMDA+GF treatment in mouse, top 50 specifically expressed genes were chosen for each cell type. We performed RNA velocity using 185 genes after setting S > 10 and 1 < U < 2000.

#### Comparative analysis of MG from zebrafish development and injury models

We first used Seurat to identify cell subpopulations for zebrafish development (Supplementary Data 18). Single-cell clustering was visualized through UMAP (*72*). Based on known marker genes of retinal cell types (Supplementary Data 7), we annotated cell types for each cell subpopulation. We then selected 12,680 primary RPC (retinal progenitor cells), neurogenic RPC cells and MG to construct a trajectory and identify pseudo-temporally changed genes (PCGs) during MG development (Supplementary Data 19). To compare each injury model with retinal development in the zebrafish, we combined 12,680 developmental cells with those MG cells in NMDA treatment, light damage and T+R treatment. Meanwhile, we identified the overlapped PCGs between retinal development and injury model. Using overlapped PCGs, we reconstructed MG trajectory for each injury model combined with cells obtained in retinal development. This analysis was performed using Monocle software (*70*).We then identified branch-specific expressed genes (BEGs) through a similar method performed as described above.

#### Mapping and normalization of ATAC-Seq

We measured 20 zebrafish GFP+ and 20 mouse GFP+ samples using ATAC (Assay for Transposase-Accessible Chromatin) sequencing technology (Table S1, Supplementary Data 1). After removing adaptors using cutadapt (*73*), 50bp paired-end ATAC-Seq reads from zebrafish and mouse were separately aligned to GRCz10/danRer10 and GRCm38/mm10 reference genome using Bowtie2 with default parameters (*74*). We performed a similar analysis as our previous publication on ATAC-Seq analysis (*75*). Briefly, we filtered reads from chromosome M and Y, and included high mapping-quality reads (MAPQ score > 10) for further analysis. Duplicate reads were removed using Picard tools MarkDuplicates (http://broadinstitute.github.io/picard/). ATAC-Seq peak regions were called using MACS2 with parameters --nomodel --shift - 100 --extsize 200 (*76*). The blacklisted regions in mouse were excluded from peak regions (https://www.encodeproject.org/annotations/ENCSR636HFF/). We further merged the ATAC-Seq peaks from each sample to obtain a union set of peaks. In total we identified 248,761 peaks in zebrafish and 119,718 peaks in mouse (Supplementary Data 20). We counted the raw fragments (*C_R_*) for each peak using HTSeq (*77*). Normalized fragments (C*_N_*) were calculated as:

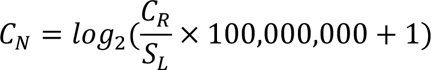

where *S_L_* is the sequencing library size of the sample.

#### ATAC-Seq footprints and differentially accessible regions

In open chromatin regions, we identified the footprints using DNase2TF (Supplementary Data 21). Meanwhile, we identified differentially accessible regions (DARs). To identify DARs, we included 150,148 peaks in zebrafish which have > 6 normalized fragments, and 104,442 peaks in mouse which have > 5 normalized fragments on average across all samples. EdgeR was used to identify DARs in comparison of each time point of treatment with the corresponding peaks from control ATAC-Seq samples (*65*). In the zebrafish, we chose fold change > 1.5 and FDR < 0.05 as the criteria of DARs. In the mouse, the criteria of DARs is fold change > 2 and FDR < 0.01. Totally, we identified 56,939 DARs in the zebrafish and 77,447 DARs in the mouse (Supplementary Data 22).

We calculated the correlation of chromatin accessibility between NMDA treatment and light damage models. Correlated peaks were identified if correlation >0.5. The cutoff was determined based on the correlation distribution of shuffled chromatin accessibility. From all DARs, we identified 24,515 correlated DARs in the zebrafish and 52,297 correlated DARs in the mouse (Supplementary Data 23).

#### Reconstructing an integrated regulatory network

In order to reconstruct the regulatory network by integrating bulk RNA-Seq, scRNA-Seq and ATAC-Seq data to, we developed a new method named ***I***ntegrated ***Re***gulatory ***N***etwork ***A***nalysis (IReNA) (Fig. S16a).

First, we identified the relevant transcription factors (TFs) in the system following the criteria: (a) The TFs were highly expressed: expression level > 5 FPKM for bulk RNA-Seq, or > 0.1 fraction of expressed cells for scRNA-Seq; (b) Correlated expression patterns between NMDA and light damage model: correlation > 0.5 in bulk RNA-Seq, or > 0.2 in scRNA-Seq; (c) Differentially expressed in treatment vs. control (identified from above RNA-Seq and scRNA-Seq analysis) or highly expressed in MG. We identified 192 candidate TFs in the zebrafish and 212 candidate TFs in the mouse (Supplementary Data 27). Corresponding binding motifs of the TFs were extracted from TRANSFAC 2018.3 version.

Next, we clustered all differentially or highly expressed genes as well as candidate TFs into ten modules through K-means clustering in each species. Seven modules were from scRNA-Seq and three modules were only from bulk RNA-Seq.

For each gene in the modules, we used FIMO to identify DNA motifs within its footprints in the correlated DARs (P value < 1.0E-04) (*78*). Meanwhile, we calculated footprint occupancy score (FOS) for each footprint in each ATAC-Seq sample. The FOS was calculated as previously described (*75*),

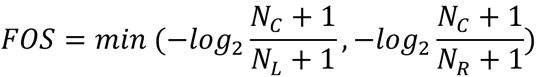

where *N*_C_ indicates the counts of ATAC-Seq inserts in the central region of the TF motif. The size of the central region is equal to the length of TF motif. *N*_L_ and *N*_R_ are separately 1/3 of the counts of ATAC-Seq inserts in the left and right flanking regions of the TF motif, as 3 times of the size of the central region were chosen as the size of the flanking region. To further refine regulatory relationship between TFs and targets, we included binding motifs with FOS > 1 in any one of ATAC-Seq samples. We obtained regulatory relationships between genes and TFs.

Finally, we used expression correlation to determine active (positive correlation) or repressive (negative correlation) regulations of TFs to target genes. If both TF and target gene of one regulatory pair are RNA-Seq specific, the correlation was calculated using bulk RNA-Seq data, otherwise single-cell expression correlation was used. For expression correlation calculating from bulk RNA-Seq data, > 0.5 and < −0.5 were selected to active and repressive regulations, respectively. For single-cell correlation, > 0.5 and < −0.5 were selected to active and repressive regulations, respectively. We identified 26,083 regulatory relationships in the zebrafish and 30,177 relationships in the mouse (Supplementary Data 26).

#### Enrichment of TFs regulating each gene module

For each gene module, we calculated the probability of the TF regulating module:

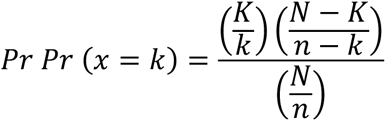

where *N* is the total number of regulations, *K* is the number of all regulations targeting module A, *n* is the number of regulations from the TF, *k* is the number of regulations from the TF to module A, and 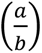 is a binomial coefficient. We separately calculated p-value for active and repressive regulations. FDR was further calculated based on p-value.

Using FDR < 10E-6 in the zebrafish and FDR < 10E-5 in the mouse, we separately identified 97 zebrafish TFs and mouse 157 TFs which regulate at least one gene module (Supplementary Data 27). Furthermore, we extracted a regulatory network consisting of these TFs from the regulatory network of candidate genes (Fig. S18 for zebrafish, Fig. S19 for mouse, Supplementary Data 28).

#### Constructing an intramodular regulatory network

To determine the regulatory relationships between two modules, we calculated the significance of regulations from module A to module B using hypergeometric test:

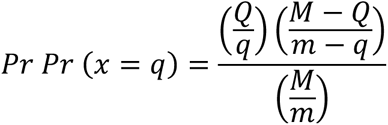

where *M* is the total number of regulations, *Q* is the number of all regulations from module A, *m* is the number of all regulations targeting module B, *q* is the number of regulations from module A to module B, and 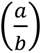 is a binomial coefficient. We assessed the significance of active and repressive regulations, respectively. We set FDR < 0.02 in zebrafish and FDR < 0.001 in mouse as the cutoff to obtain significant regulations between modules (Figure 6d, e).

#### Features of the TFs in regulatory network

We calculated the regulation degrees of each enriched TF in the regulatory network. The regulation degrees include total degree (number of regulations connecting with TFs), active indegree (number of regulations activated by TFs), repressive indegree (number of regulations repressed by TFs), active outdegree (number of regulations activating TFs), repressive outdegree (number of regulations repressing TFs). For the regulatory network of candidate genes as well as the regulatory network of enriched TFs, we separately calculated regulation degrees (*D_R_*). Normalized regulation degrees (*D_N_*) were calculated as:

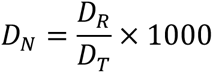

where *D_T_* is the total number of regulations in the gene regulatory network or the network of enriched TFs. We also included other features of the TFs, such as modules regulated by the TFs. Bulk and single-cell expression of the TFs is shown in Supplementary Data 29 as well.

#### Mapping of new single cells to the original trajectory

To map new single cells to the original trajectory, we first calculated Person’s correlation coefficient between new cells and original cells. To calculate the expression correlation, we used top variable genes which were used for constructing the trajectory. For each new cell, the most correlated cell from the original trajectory was identified. Correspondingly, the location and state of the original cell were assigned to the new cell. Based on this, we obtained the distribution of new cells in the original trajectory (Figure 6o).

## Supporting information

Summary of all supplemental tables and datasets

Supplementary Data 1. Information on all samples and libraries measured by RNA-Seq, ATAC-Seq and scRNA-Seq

Supplementary Data 2. Gene expression data for all RNA-Seq samples, and list of GFP+ or GFP- specific expressed genes

Supplementary Data 3. List of differentially expressed genes following NMDA treatment and light damage in mouse and zebrafish

Supplementary Data 4. List of correlated and differentially expressed genes in mouse and zebrafish MG

Supplementary Data 5. List of genes showing species-specific changes in expression pattern or level in MG

Supplementary Data 6. tSNE clustering of retinal cells in in mouse, chicken and zebrafish

Supplementary Data 7. List of known specific marker genes for retinal cell types

Supplementary Data 8. Marker genes of retinal cell types in each model from mouse, chicken and zebrafish

Supplementary Data 9. Species-conserved marker genes of each retinal cell type

Supplementary Data 10. List of pseudo-temporally changed genes in mouse, chicken and zebrafish nonMG measured by scRNA-Seq

Supplementary Data 11. Trajectory and pseudotime of mouse, chick and zebrafish MG

Supplementary Data 12. List of differentially expressed genes in gliotic MG vs resting MG

Supplementary Data 13. List of branch-specific expressed genes in zebrafish NMDA, LD and T+R treatment

Supplementary Data 14. List of branch-specific genes in chick MG

Supplementary Data 15. List of pseudo-temporally changed genes in mouse, chicken and zebrafish MG

Supplementary Data 16. List of correlated and pseudo-temporally changed genes in mouse, chicken and zebrafish MG

Supplementary Data 17. List of genes showing species-specific changes in single-cell expression pattern or level in mouse, chick and zebrafish MG

Supplementary Data 18. UMAP clustering of retinal cells in zebrafish development

Supplementary Data 19. List of pseudo-temporally changed genes in zebrafish MG development

Supplementary Data 20. Counts of open chromatin regions in zebrafish and mouse MG identified by ATAC-Seq

Supplementary Data 21. Transcription factor footprints detected in open chromatin regions of zebrafish and mouse MG

Supplementary Data 22. Differentially accessible regions in zebrafish and mouse

Supplementary Data 23. Correlated and differentially accessible regions in zebrafish and mouse

Supplementary Data 24. List of correlated and differentially expressed genes identified by RNA-seq and scRNA-seq data analysis

Supplementary Data 25. Gene list of ten modules in zebrafish and mouse MG

Supplementary Data 26. Regulatory network of regulators and genes

Supplementary Data 27. Significance of candidate regulators regulating gene modules

Supplementary Data 28. Regulatory network of enriched regulators in zebrafish and mouse

Supplementary Data 29. Features of enriched regulators in zebrafish and mouse

Supplementary Data 30. List of differentially expressed genes in MG following hmga1a knockdown or yap1 inhibition

Supplementary Data 31. List of cluster markers from flow-sorted 21 days after NFI TKO and NMDA+GF treatment in mouse

Table S1. Statistics of bulk RNA-Seq and ATAC-Seq data

Table S2. List of antibodies used in the study.

Table S3. List of qPCR primers used in the study

## Supplementary Tables

**Table S1: Statistics of bulk RNA-Seq and ATAC-Seq data.**

**Table S2. List of antibodies used in the study.**

**Table S3. List of qPCR primers used in the study.**

## Supplementary Information

**Supplementary Data 1. Information on all samples and libraries measured by RNA-Seq, ATAC-Seq and scRNA-Seq.** Tab 1 (RNAseq): Information of RNA-Seq samples. Tab 2 (ATACseq): Information of ATAC-Seq samples. Tab 3 (scRNAseq): Information of scRNA-Seq samples.

**Supplementary Data 2. Gene expression data for all RNA-Seq samples, and list of GFP+ or GFP-specific expressed genes.** Tab 1 (Mouse): List of genes specifically expressed in GFP+ and GFP-samples in mouse. Tab 2 (Zebrafish): List of genes specifically expressed in GFP+ and GFP-samples in zebrafish. FPKM (Fragments Per Kilobase Of Exon Per Million) values of gene expression are given starting from column G.

**Supplementary Data 3. List of differentially expressed genes (DEGs) following NMDA treatment and light damage in mouse and zebrafish.** Tab 1 (Mouse.MG): List of differentially expressed genes in MG in mouse NMDA and light damage (LD); Tab 2 (Zebrafish.MG): List of differentially expressed genes in MG in zebrafish NMDA and light damage; Tab 3 (Mouse.nonMG): List of differentially expressed genes in non-MG in mouse NMDA and light damage (LD); Tab 4 (Zebrafish.nonMG): List of differentially expressed genes in non-MG in zebrafish NMDA and light damage. Statistics (log.fold.change, P.value and FDR) of each time point of treatment vs. control are given. FPKM values of gene expression are given in the tables starting from column AC.

**Supplementary Data 4. List of correlated and differentially expressed genes (cDEGs) in mouse and zebrafish.** Tab 1 (Mouse): List of cDEGs in mouse. Tab 2 (Zebrafish): List of cDEGs in zebrafish. FPKM values of gene expression are given starting from column E.

**Supplementary Data 5. List of genes showing species-specific changes in expression pattern or level.** Tab 1 (Differentially.expressed.genes): List of genes showing species-specific changes in expression pattern; FPKM values of gene expression in GFP+ samples are given starting from column F. Tab 2 (Highly.expressed.genes): List of genes showing species-specific changes in baseline expression. Ratio of FPKM in GFP+ vs. corresponding GFP-samples are given starting from column F.

**Supplementary Data 6. tSNE clustering of retinal cells in mouse, chicken and zebrafish.** Tab 1 (Mouse.NMDA): tSNE clustering of retinal cells in mouse NMDA treatment. Tab 2 (Mouse.LD): tSNE clustering of retinal cells in mouse light damage (LD). Tab 3 (Chick.NMDA.GF): tSNE clustering of retinal cells in chick NMDA or growth factor (GF) treatment. Tab 4 (Zebrafish.NMDA): tSNE clustering of retinal cells in zebrafish NMDA treatment. Tab 5 (Zebrafish.LD): tSNE clustering of retinal cells in zebrafish light damage. Tab 6 (Zebrafish.T+R): tSNE clustering of retinal cells in zebrafish TNFa+RO4929097 (T+R) treatment.

**Supplementary Data 7. List of known specific marker genes for retinal cell types.** Tab 1 (Mouse): Known specific marker genes for mouse retinal cell types. Tab 2 (Chick): Known specific marker genes for chick retinal cell types. Tab 3 (Zebrafish): Known specific marker genes for zebrafish retinal cell types. Tab 4 (Abbreviation.of.cell.types): Full name of retinal cell types in abbreviation.

**Supplementary Data 8. Specific marker genes of retinal cell types from mouse, chicken and zebrafish.** Tab 1 (Mouse): Specific marker genes of retinal cell types from mouse NMDA treatment and light damage (LD). Tab 2 (Chick): Specific marker genes of retinal cell types from chick NMDA/growth factor (GF) treatment. Tab 3 (Zebrafish): Specific marker genes of retinal cell types from zebrafish NMDA treatment, light damage (LD) and TNFα+RO4929097 (T+R) treatment.

**Supplementary Data 9. Species-conserved marker genes of each retinal cell type.** Tab 1 (Shared.by.mouse.and.chick): Conserved marker genes of retinal cell types shared by mouse and chick. Tab 2 (Shared.by.mouse.and.zebrafish): Conserved marker genes of retinal cell types shared by mouse and zebrafish. Tab 3 (Shared.by.chick.and.zebrafish): Conserved marker genes of retinal cell types shared by chick and zebrafish. Tab 3 (Shared.by.three.species): Conserved marker genes of retinal cell types shared by mouse, chick and zebrafish.

**Supplementary Data 10. List of pseudo-temporally changed genes in mouse, chicken and zebrafish non-MG measured by scRNA-Seq.** List of genes which are changed during pseudotime of NMDA treatment or light damage (LD) in non-muller glia cells (GABAergic AC, Glycinergic AC, RGC, cones, rods and microglia) from mouse, zebrafish and chick.

**Supplementary Data 11. Trajectory and pseudotime of mouse, chick and zebrafish MG.** Tab 1 (Mouse). Trajectory and pseudotime of mouse MG. Tab 2 (Chick): Trajectory and pseudotime of chick MG. Tab 3 (Zebrafish): Trajectory and pseudotime of zebrafish MG.

**Supplementary Data 12. List of differentially expressed genes (DEGs) in gliotic MG vs resting MG.** Tab 1 (Mouse): List of DEGs in activated MG vs resting MG in mouse; Tab 2 (Chick): List of DEGs in gliotic MG vs. resting MG in chick; Tab 3 (Zebrafish): List of DEGs in gliotic MG vs. resting MG in zebrafish; Tab4 (Gliosis.in.mm.ch.zf): Comparison of DEGs shared by all models in each species of mouse, chick and zebrafish. In chick and zebrafish, resting MG and gliotic MG are from the rest branch and gliosis branch, respectively.

**Supplementary Data 13. List of branch-specific expressed genes (BEGs) in zebrafish MG following NMDA, light damage (LD) and TNFα+RO4929097 (T+R) treatment.** Tab 1 (NMDA): List of BEGs in zebrafish MG following NMDA treatment. Tab 2 (LD): List of BEGs in zebrafish MG following light damage. Tab 3 (T+R): List of BEGs in zebrafish MG following T+R treatment. Differential proportion of expressed cells in the current branch was calculated by subtracting the proportion of expressed cells in the other branches.

**Supplementary Data 14. List of branch-specific genes in chick MG.** Differential proportion of expressed cells in the current branch was calculated by subtracting the proportion of expressed cells in the other branches.

**Supplementary Data 15. List of pseudo-temporally changed genes (PCGs) in mouse, chicken and zebrafish MG.** Tab 1 (Mouse): List of PCGs in mouse MG. Tab 2 (Chick): List of PCGs in chick MG. Tab 3 (Zebrafish): List of PCGs in zebrafish MG. Normalized expression values are given in Tab 1 starting from column K, in Tab 2 starting from column K, in Tab 3 starting from column P.

**Supplementary Data 16. List of correlated and pseudo-temporally changed genes (cPCGs) in mouse, chicken and zebrafish MG.** Tab 1 (Mouse): List of cPCGs in mouse MG. Tab 2 (Chick): List of cPCGs in chick MG. Tab 3 (Zebrafish): List of cPCGs in zebrafish MG. Normalized expression values are given in Tab 1 starting from column K, in Tab 2 starting from column K, in Tab 3 starting from column P.

**Supplementary Data 17. List of genes showing species-specific changes in single-cell expression pattern or level in mouse, chick and zebrafish MG.** Tab 1 (Pseudo-temporally.changed.genes): List of genes showing species-specific changes in single-cell expression pattern. Tab 2 (Highly.expressed.genes): List of genes showing species-specific changes in single-cell baseline expression. Normalized expression values are given starting from column AC.

**Supplementary Data 18. UMAP clustering of retinal cells in zebrafish development.** Cell type and UMAP coordinates (UMAP.1, UMAP.2 and UMAP.3) of each retinal cell in zebrafish development are shown.

**Supplementary Data 19. List of pseudo-temporally changed genes in zebrafish MG development.** Normalized expression values are given starting from column G.

**Supplementary Data 20. Counts of open chromatin regions (OCRs) in zebrafish and mouse MG identified by ATAC-Seq analysis.** Tab 1 (Zebrafish): Counts of OCRs in zebrafish MG. Tab 2 (Mouse): Counts of OCRs in mouse MG. Raw counts of each peak are given starting from column H.

**Supplementary Data 21. Transcription factor footprints detected in open chromatin regions (OCRs) of zebrafish and mouse MG.** Tab 1 (Zebrafish): Transcription factor footprints detected in OCRs of zebrafish MG. Tab 2 (Mouse): Transcription factor footprints detected in OCRs of mouse MG. The column log10.p.value represents the significance of TF footprint as predicted by Dnase2tf.

**Supplementary Data 22. Differentially accessible chromatin regions (DARs) in zebrafish and mouse MG.** Tab 1 (Zebrafish): DARs in zebrafish MG. Tab 2 (Mouse): DARs in mouse MG. Normalized counts are given in Tab 1 starting from column Z, and in Tab 2 starting from column AB.

**Supplementary Data 23. Correlated and differentially accessible regions (cDARs) in zebrafish and mouse.** Tab 1 (Zebrafish): cDARs in zebrafish MG. Tab 2 (Mouse): cDARs in mouse MG. Normalized counts are given in Tab 1 starting from column Z, and in Tab 2 starting from column AB.

**Supplementary Data 24. List of correlated and differentially expressed genes identified by RNA-Seq and scRNA-Seq data analysis.** Tab 1 (Mouse): List of correlated and differentially expressed genes identified by RNA-Seq and scRNA-Seq data analysis in mouse. Tab 2 (Zebrafish): List of correlated and differentially expressed genes identified by RNA-Seq and scRNA-Seq data analysis in zebrafish. FPKM values and normalized expression values are given separately for RNA-Seq and scRNA-Seq samples.

**Supplementary Data 25. Gene list of ten modules in zebrafish and mouse MG.** Tab 1 (Zebrafish): Gene list of ten modules in zebrafish MG. Tab 2 (Mouse): Gene list of ten modules in mouse MG. Fractions of expressed cells and FPKM values are given separately for scRNA-Seq and RNA-Seq samples.

**Supplementary Data 26. Regulatory network of regulators and genes in zebrafish and mouse MG.** Tab 1 (Zebrafish): Regulatory network of regulators and genes in zebrafish MG. Tab 2 (Mouse): Regulatory network of regulators and genes in mouse MG. The last column contains information on TF binding sites in the target gene. Motif ID from TRANSFAC database. FOS, Footprint occupancy score.

**Supplementary Data 27. Significance of candidate regulators regulating gene modules in zebrafish and mouse MG.** Tab 1 (Zebrafish): Significance of candidate regulators regulating gene modules in zebrafish MG. Tab 2 (Mouse): Significance of candidate regulators regulating gene modules in mouse MG.

**Supplementary Data 28. Regulatory network of enriched regulators in zebrafish and mouse MG.** Tab 1 (Zebrafish): Regulatory network of enriched regulators in zebrafish MG. Tab 2 (Mouse): Regulatory network of enriched regulators in mouse MG. The last column contains information on TF binding sites in the target gene. Motif ID from TRANSFAC database. FOS, Footprint occupancy score.

**Supplementary Data 29. Features of enriched regulators in zebrafish and mouse MG.** Tab 1 (Zebrafish): Features of enriched regulators in zebrafish MG. Tab 2 (Mouse): Features of enriched regulators in mouse MG. Tab 3 (Zebrafish.vs.Mouse): Features of enriched regulators across species. Gene module/network indicates module/network consisting of candidate genes (including TFs). TF module/network means module/network only consisting of enriched TFs. Indegree is the number of regulations from TFs. Outdegree is the number of regulations targeting TFs. Degree of TFs regulatory network is normalized by dividing the total number of regulations. TF expression is given using fractions of expressed cells for scRNA-Seq and FPKM values for RNA-Seq samples.

**Supplementary Data 30. List of differentially expressed genes in MG following hmga1a knockdown or Yap inhibition.** zfHmgaLdSc36R1, standard control morphants in 36hr light damage. zfHmgaLdMp36R1, hmga1a morphants in 36hr light damage. zfYap1LdDm36R1, DMSO control in 36hr light damage. zfYap1LdVp36R1, Yap inhibition using verteporfin in 36hr light damage.

**Supplementary Data 31. List of cluster markers from flow-sorted 21 days after NFI TKO and NMDA+GF treatment in mouse.** Cluster markers were identified through the roc (receiver operating characteristic) approach.

## ACKNOWLEDGMENTS

We thank J. Nathans, A. Kolodkin, and W. Yap for their comments on the manuscript. We thank Berrnard Kulemeka and the Freimann Life Science Center staff for breeding and raising the zebrafish strains used in this study. We thank X. Zhou and J. Wang for curation of the RNA and ATAC-Seq data at St. Jude.

## Author contributions

D.R.H., S.B., J.Q., A.J.F., and J.A. conceived of the project and supervised the research and experimental design. T.H. generated all RNA-Seq and ATAC-Seq data, conducted all functional studies in mouse and analyzed scRNA on *Nfia/b/x*-deficient mouse retina and developing zebrafish retina. J.W. performed all analysis of RNA-Seq and ATAC-Seq data, developed IReNA, and identified gene regulatory networks in zebrafish and mouse. F.W. and G.W. analyzed data from developing zebrafish retina. P.B. and M.L. performed light damage and NMDA damage paradigms in zebrafish and the TNFa+RO4929097 co-injections and cell sorting to isolate MG and neuronal fractions for bulk RNA-Seq, single-cell RNA-Seq, and ATAC-Seq. P.B. and M.J. performed morpholino and verteporfin studies in zebrafish. L.Z., C.Z., C.S., S.Y., F.R., V.T., and T.P. all contributed to analysis of mouse functional data. C.S., N.S., and C.K. contributed to analysis of mouse gene expression data. L.J.T. and I.P. conducted functional analysis in chick.

## Data and code availability

All RNA-Seq and ATAC-Seq data are deposited in GEO (<u>GSE135406</u>). Bulk RNA-Seq, bulk ATAC-Seq and scRNA-Seq data can be queried interactively at https://proteinpaint.stjude.org/F/2019.retina.scRNA.html. Source code for data analysis are available from https://github.com/jiewwwang/Single-cell-retinal-regeneration.

---

**Fig. S1.**
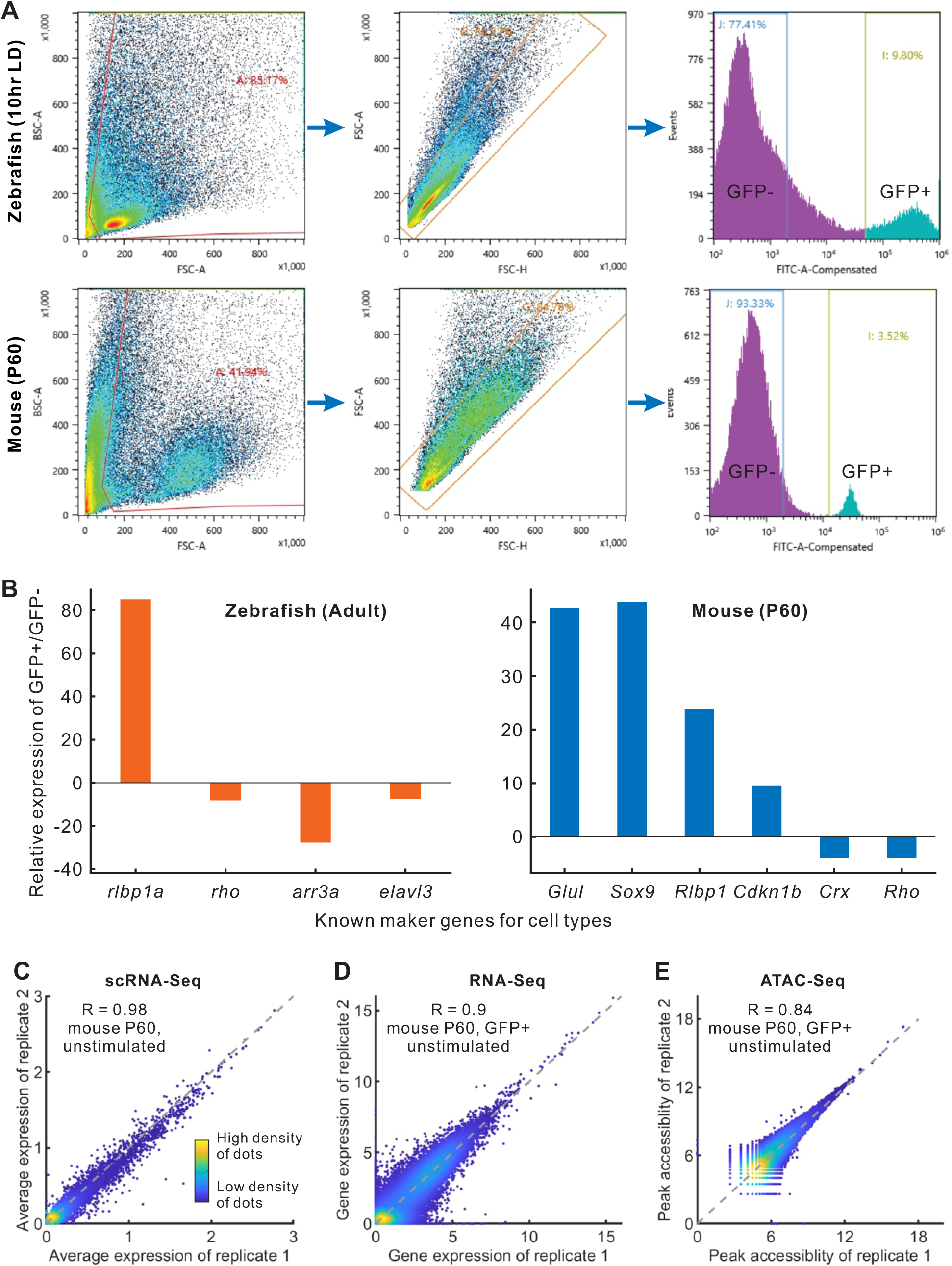
Quality of measured samples and data. **(A)**Fluorescence activated cell sorting (FACS) strategy for isolation of GFP+ (MG) and GFP- (non-MG) cells. **(B)** Expressions of marker genes measured by RT-qPCR from FACS in adult zebrafish and P60 mouse. **(C-E)** Correlation between two replicates from mouse P60 measured by scRNA-Seq, RNA-Seq and ATAC-Seq.

**Fig. S2.**
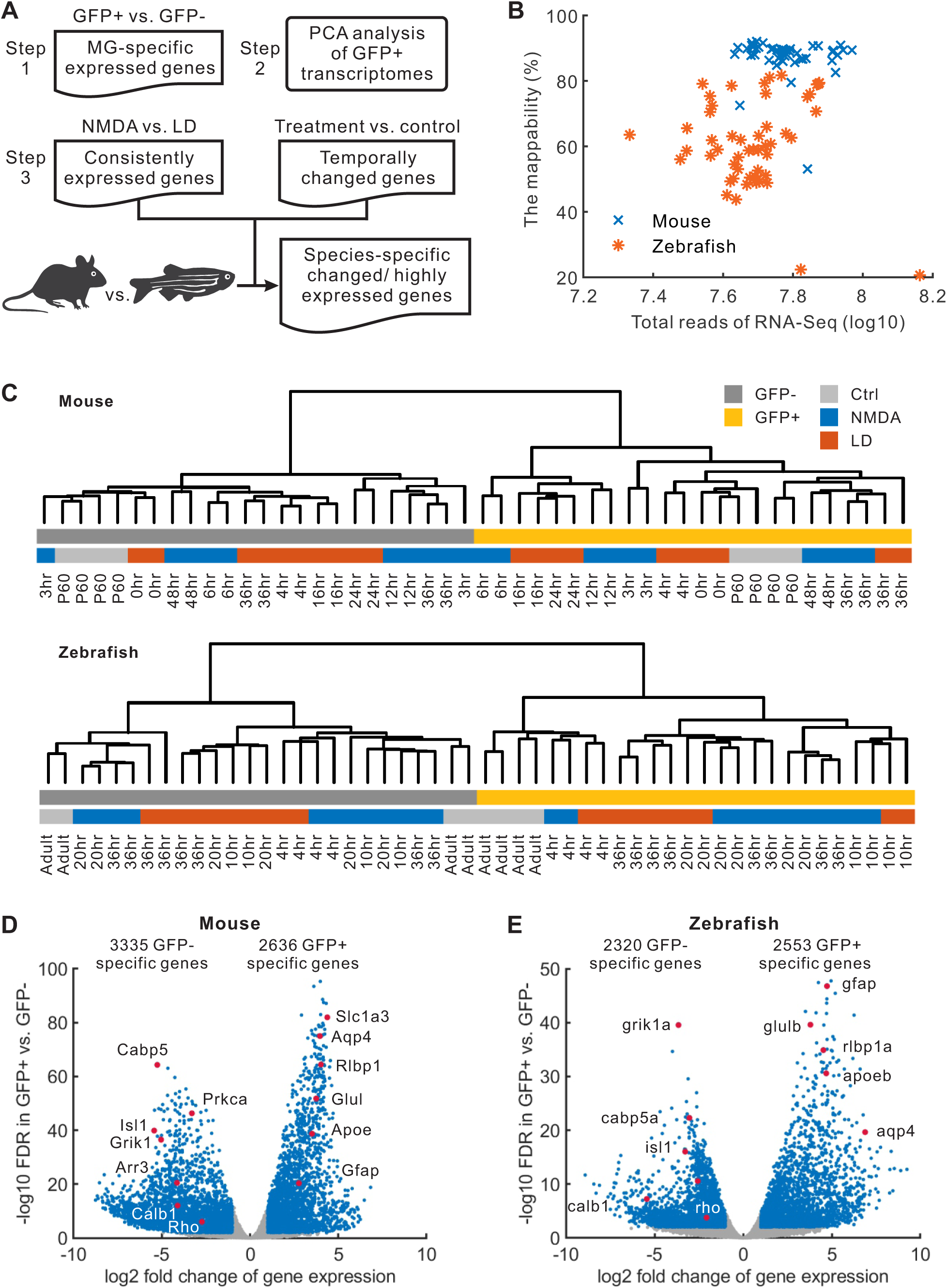
Comparison of GFP+ and GFP-samples. **(A)**Analysis workflow of RNA-Seq data. After identifying MG-specific expressed genes, we performed PCA (principal component analysis), and identified correlated and differentially expressed genes (DEGs) across multiple time points of NMDA and light damage (LD) models in mouse and zebrafish. **(B)** Mappability of RNA-Seq data. **(C)** Hierarchical clustering of GFP+ and GFP-samples based on expression profiles of all genes. **(D, E)** Volcano plot of DEGs between GFP+ and GFP-samples in zebrafish and mouse. Blue dots represent significantly DEGs (fold change > 2 and FDR < 0.01).

**Fig. S3.**
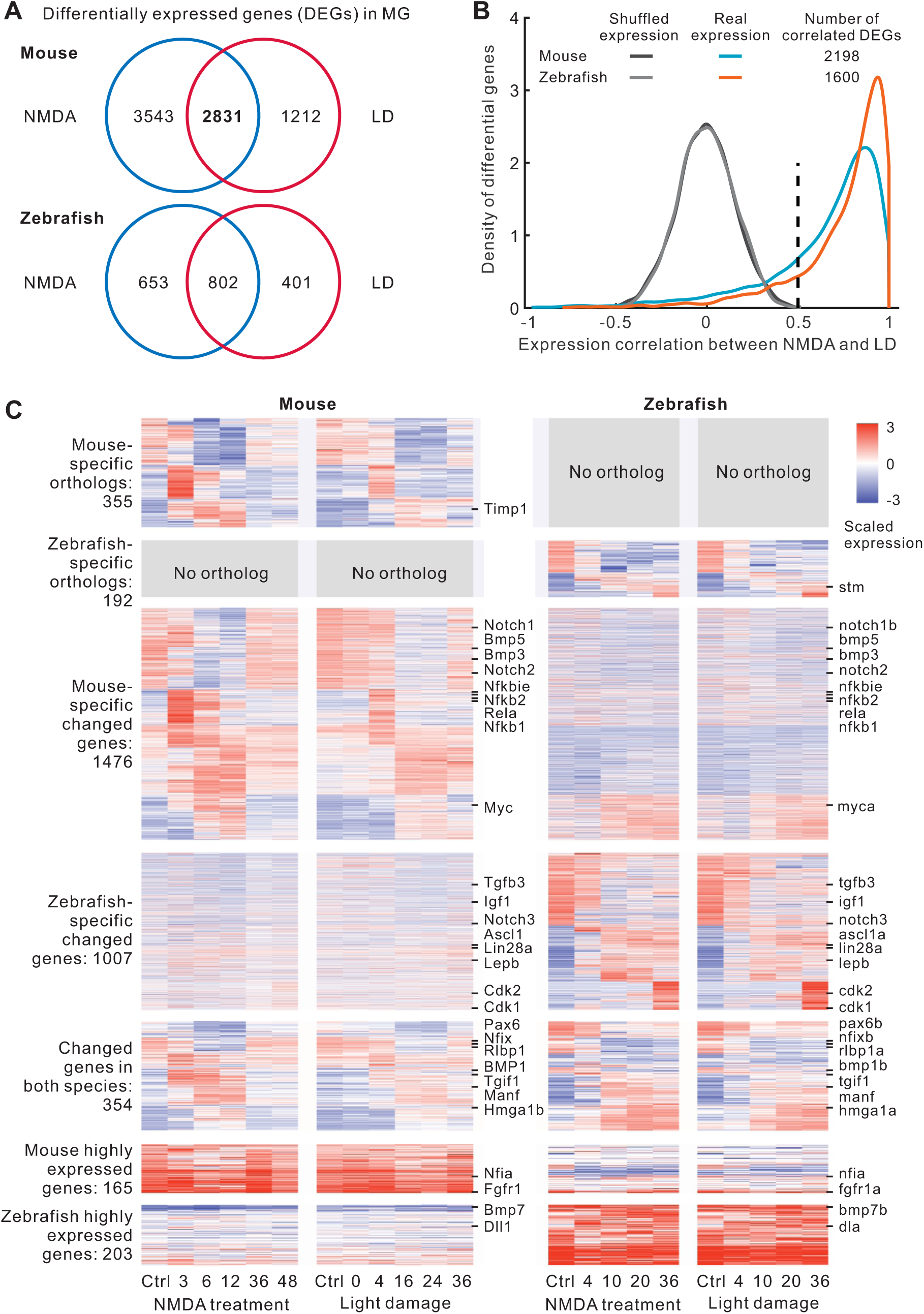
Identification of differentially expressed genes (DEGs) in mouse and zebrafish MG using bulk RNA-seq data after injury. **(A)**Venn diagram of the number of significantly DEGs following NMDA and light damage (LD) in mouse/zebrafish MG (fold change > 1.5 and FDR < 0.05). **(B)** Correlation of DEGs between two injury models. Shared DEGs in the mouse and all DEGs in the zebrafish were used to identify correlated DEGs (correlation > 0.5). **(C)** Heatmap of four broad categories of changed or highly expressed genes in each species. Scaled expression is shown for DEGs in the MG samples across different timepoints of injuries. Relative expression is shown for highly expressed genes in the MG samples as compared with MG-depleted samples in zebrafish and mouse.

**Fig. S4.**
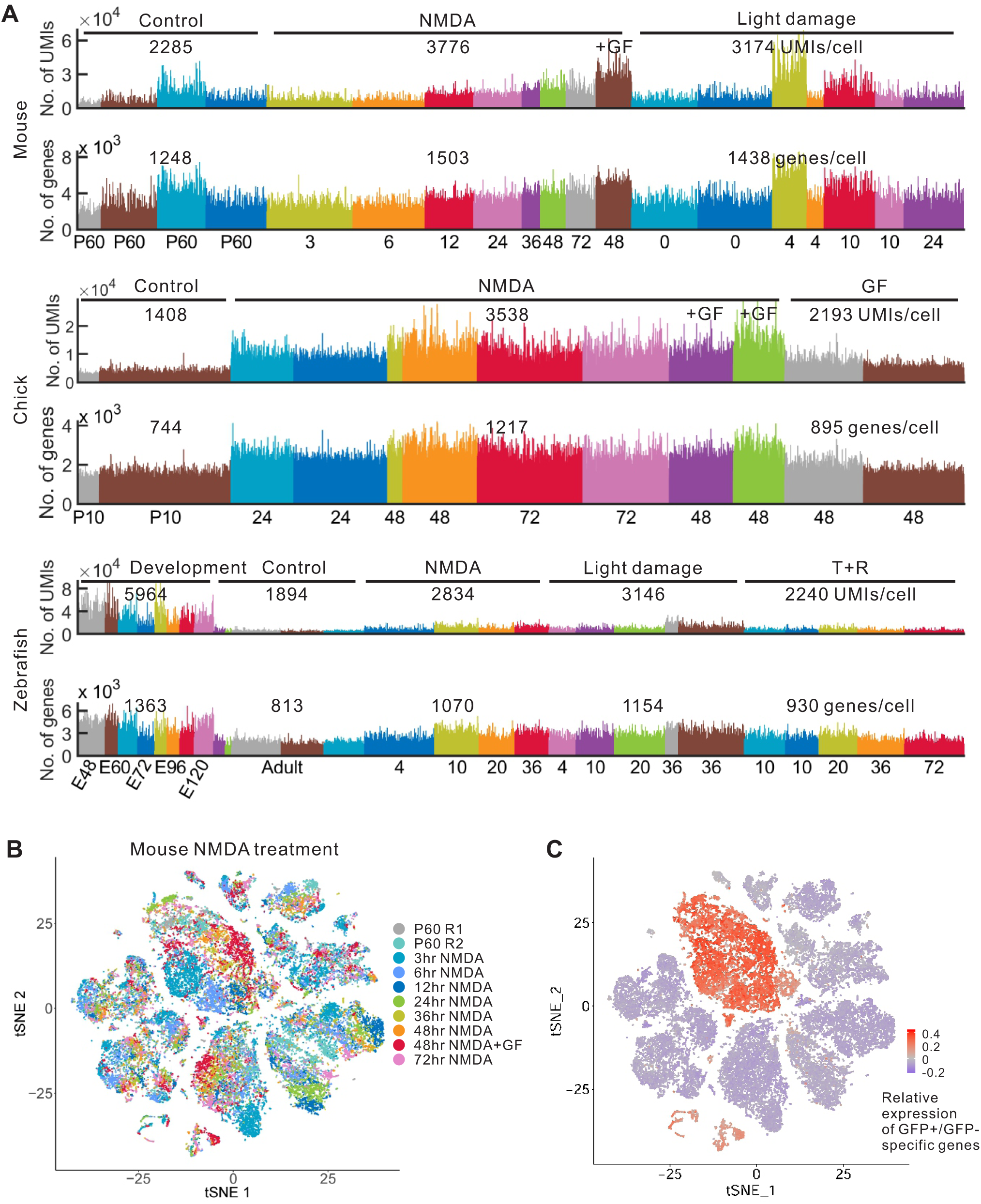
Quality of single-cell RNA-Seq data. **(A)**Number of genes and unique molecular identifiers (UMIs) in single cells in three species. Each bar represents one cell. GF, growth factor (insulin+FGF2). T+R, TNFα and RO4929097 treatment. **(B)** Clustering of mouse retinal cells from NMDA treatment. Cells are colored by samples. **(C)** Identification of mouse MG clusters using differentially expressed genes obtained from comparison of GFP+ and GFP-bulk RNA-Seq samples.

**Fig. S5.**
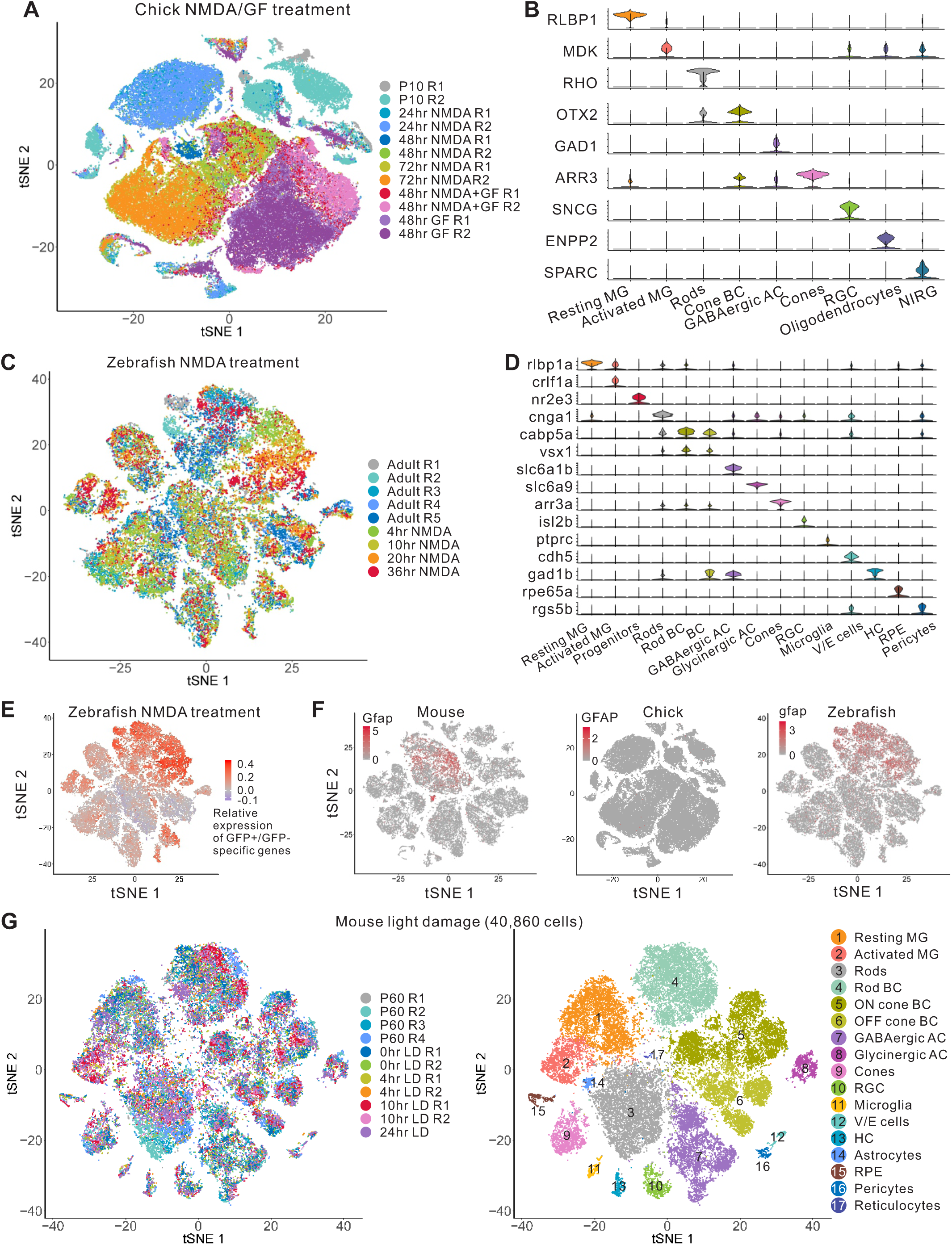
tSNE view of whole retinal single cells. **(A)**Aggregation of chick retinal single cell samples after NMDA and growth factor (GF) treatment. **(B)** Expression of cell type-specific markers in the chick. **(C)** tSNE view of zebrafish retinal single cells after NMDA treatment. **(D)** Expression of cell type-specific markers in zebrafish NMDA treatment samples. **(E)** Identification of zebrafish MG using differentially expressed genes obtained from comparison of GFP+ and GFP-bulk RNA-Seq samples. **(F)** Single-cell expression of *Gfap/GFAP/gfap* in mouse, chick and zebrafish after NMDA treatment. **(G)** tSNE view of mouse retinal cells from light damage samples. MG, Müller glia; BC, bipolar cell; AC, amacrine cell; RGC, retinal ganglion cell; HC, horizontal cell; NIRG, non-astrocytic inner retinal glia; V/E, vascular/endothelial cell; RPE, retinal pigment epithelium.

**Fig. S6.**
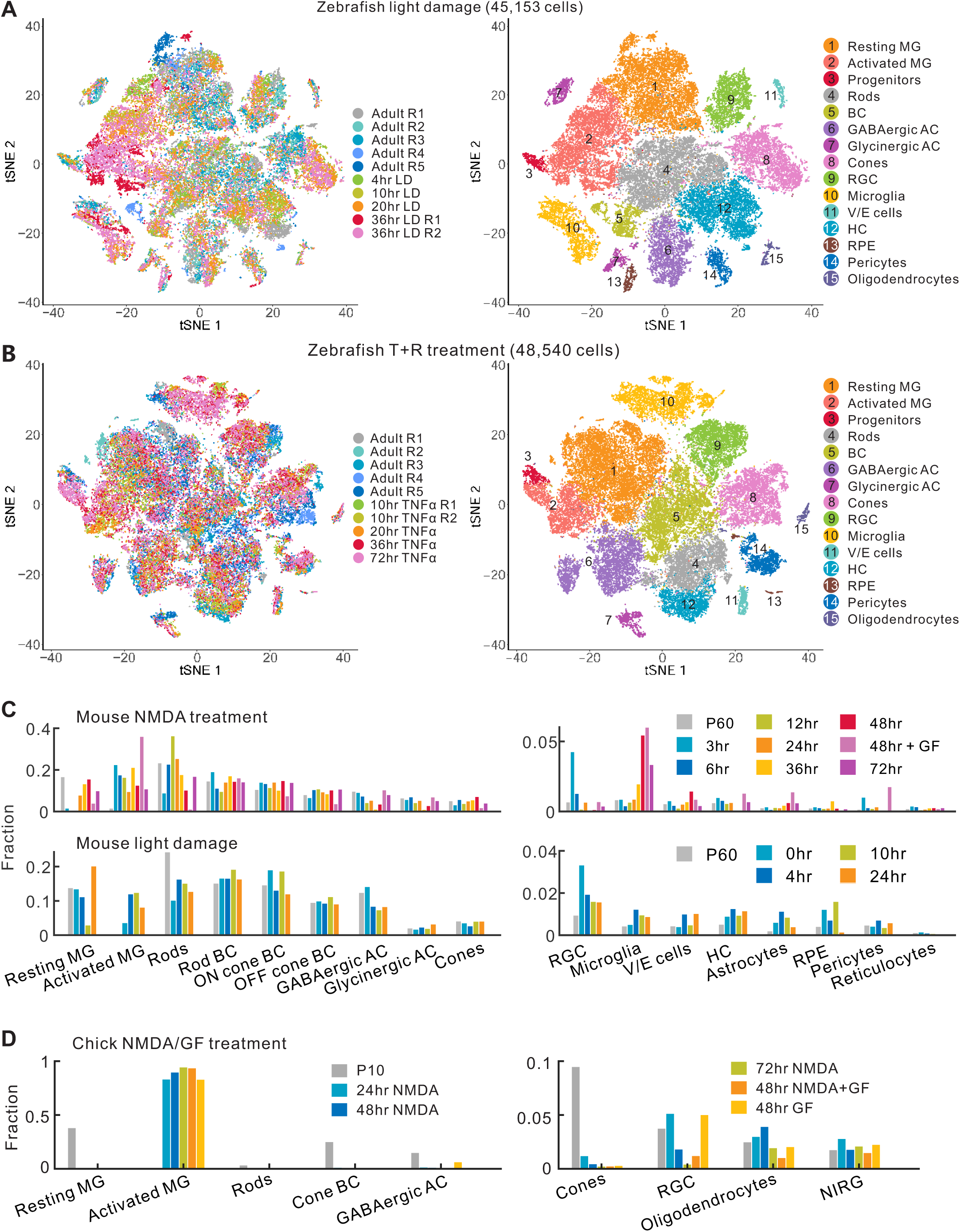
Clustering of zebrafish retinal single cells after light damage and TNFa+RO4929097 (T+R) treatment. **(A)**tSNE-view of zebrafish retinal single cells after light damage. **(B)** tSNE-view of zebrafish retinal single cells after T+R treatment. **(C)** Quantification of the relative abundance of mouse cell types following NMDA treatment and light damage. **(D)** Quantification of the relative abundance of chick cell types following NMDA/growth factor (GF) treatment. MG, Müller glia; BC, bipolar cell; AC, amacrine cell; RGC, retinal ganglion cell; HC, horizontal cell; V/E, vascular/endothelial cell; RPE, retinal pigment epithelium.

**Fig. S7.**
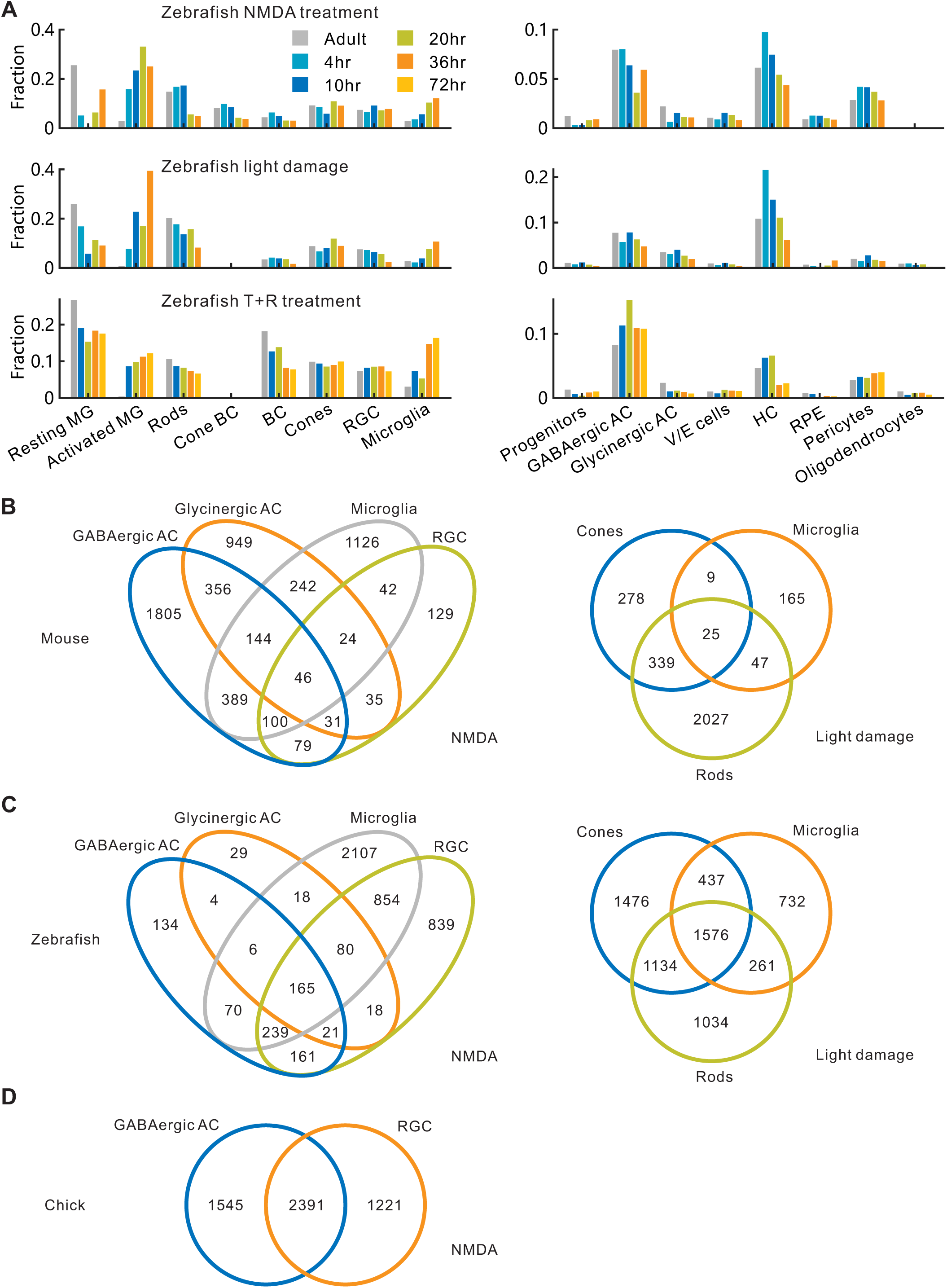
Comparison of retinal cell types across species. **(A)**Quantification of the relative abundance of zebrafish cell types following injuries or TNFa+RO4929097 (T+R) treatment. **(B-D)** Venn diagrams of pseudo-temporally changed genes in retinal cell types primarily targeted by NMDA or light damage in mouse (**B**), zebrafish (**C**) and chick (**D**). MG, Müller glia; BC, bipolar cell; AC, amacrine cell; RGC, retinal ganglion cell; HC, horizontal cell; V/E, vascular/endothelial cell; RPE, retinal pigment epithelium.

**Fig. S8.**
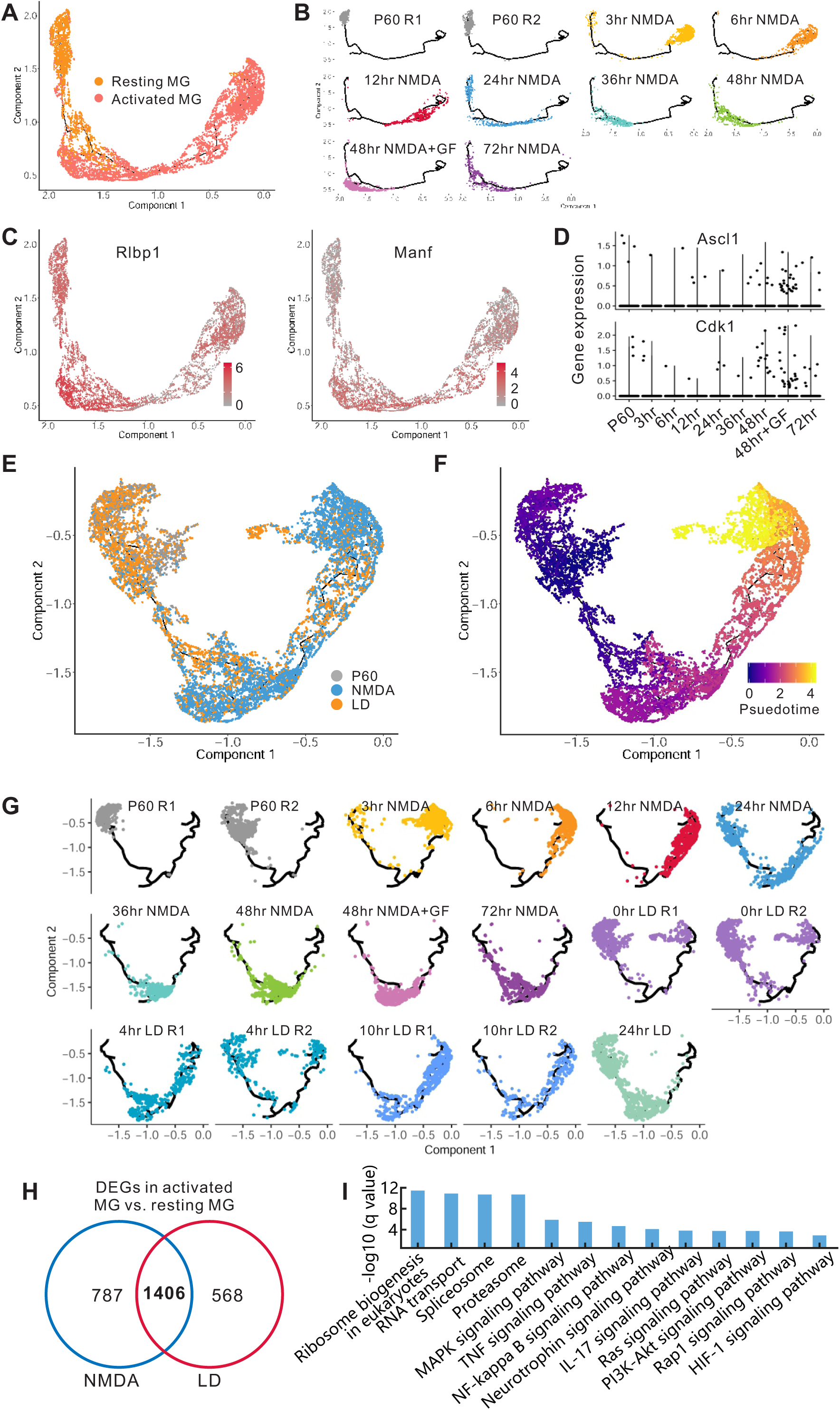
Single-cell trajectory and features of mouse MG. **(A)**Distribution of resting and activated MG in the single-cell trajectory. **(B)** Distribution of individual MG in the trajectory following NMDA treatment. **(C)** Expression of marker genes in mouse MG trajectory following NMDA treatment. **(D)** Expression of *Ascl1* and *Cdk1* in MG cells at real time points with NMDA treatment. GF, growth factor (insulin+FGF2). **(E, F)** Single-cell trajectory and pseudotime of mouse MG following NMDA treatment and light damage (LD). **(G)** Distribution of mouse MG from each sample in the trajectory. **(H)** Comparison of the number of differentially expressed genes (DEGs) from activated MG vs. resting MG between mouse NMDA and LD injuries. **(I)** Enriched functions of DEGs in activated MG vs. resting MG which are shared by both NMDA and light damages.

**Fig. S9.**
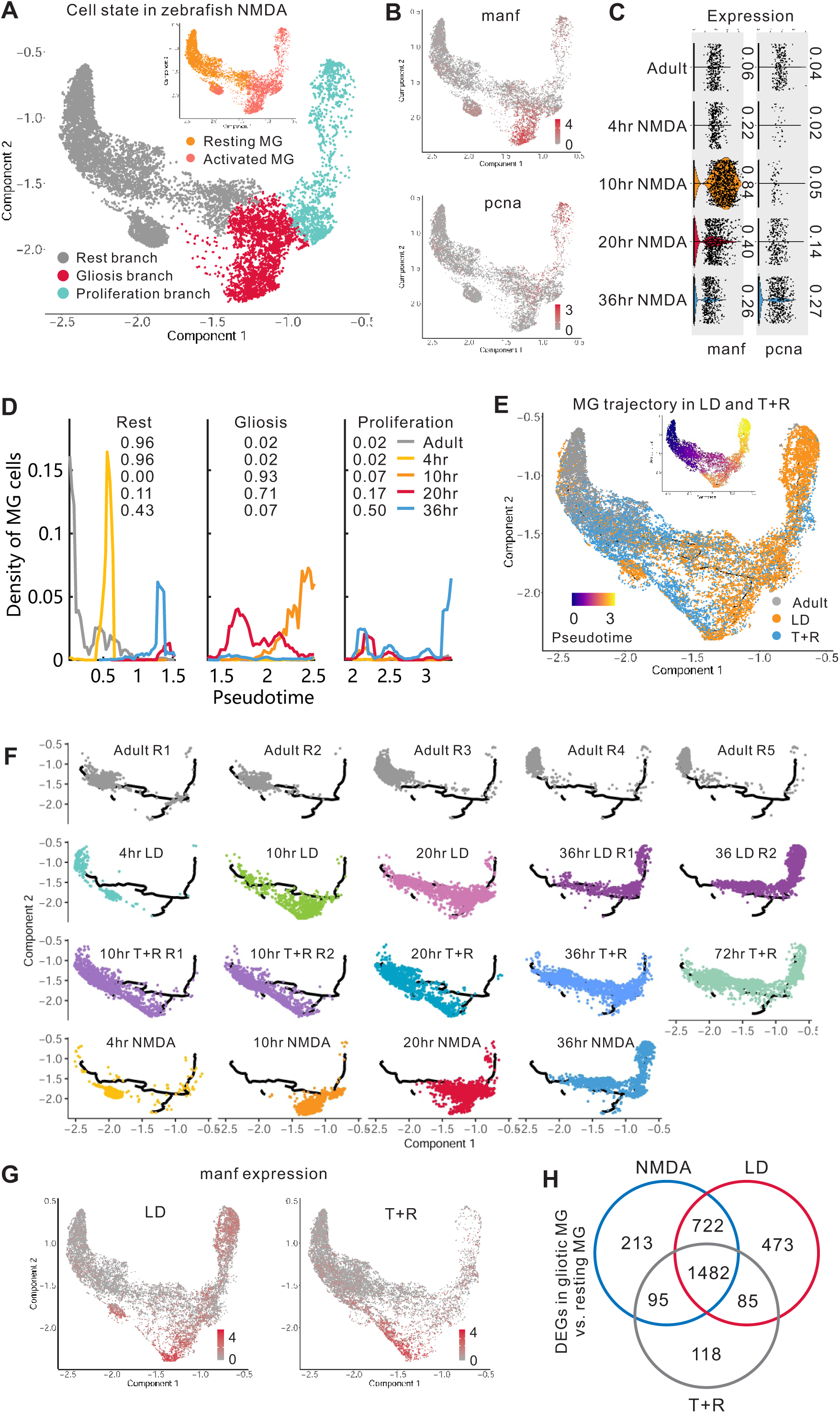
Single-cell trajectory and features of zebrafish MG. **(A)**Cell state of zebrafish MG following NMDA treatment. Inserted figure shows the distribution of resting and activated MG in the trajectory. **(B, C)**Expressions of gliotic (*manf*) and proliferation (*pcna*) markers in zebrafish in the single-cell trajectory (**B**) and in each time point of NMDA treatment (**C**). Fraction of expressed cells is shown. **(D)** The distribution density of zebrafish MG across pseudotime. **(E)** Single-cell trajectory of zebrafish MG following LD or T+R treatment. **(F)** Distribution of MG from each sample in the trajectory with LD or T+R treatment. **(G)** *manf* expression in MG following LD or T+R treatment. **(H)** Venn diagram of the number of differentially expressed genes (DEGs) from activated MG vs. resting MG between NMDA, LD and T+R treatments. LD, light damage; T+R, TNFa+RO4929097.

**Fig. S10.**
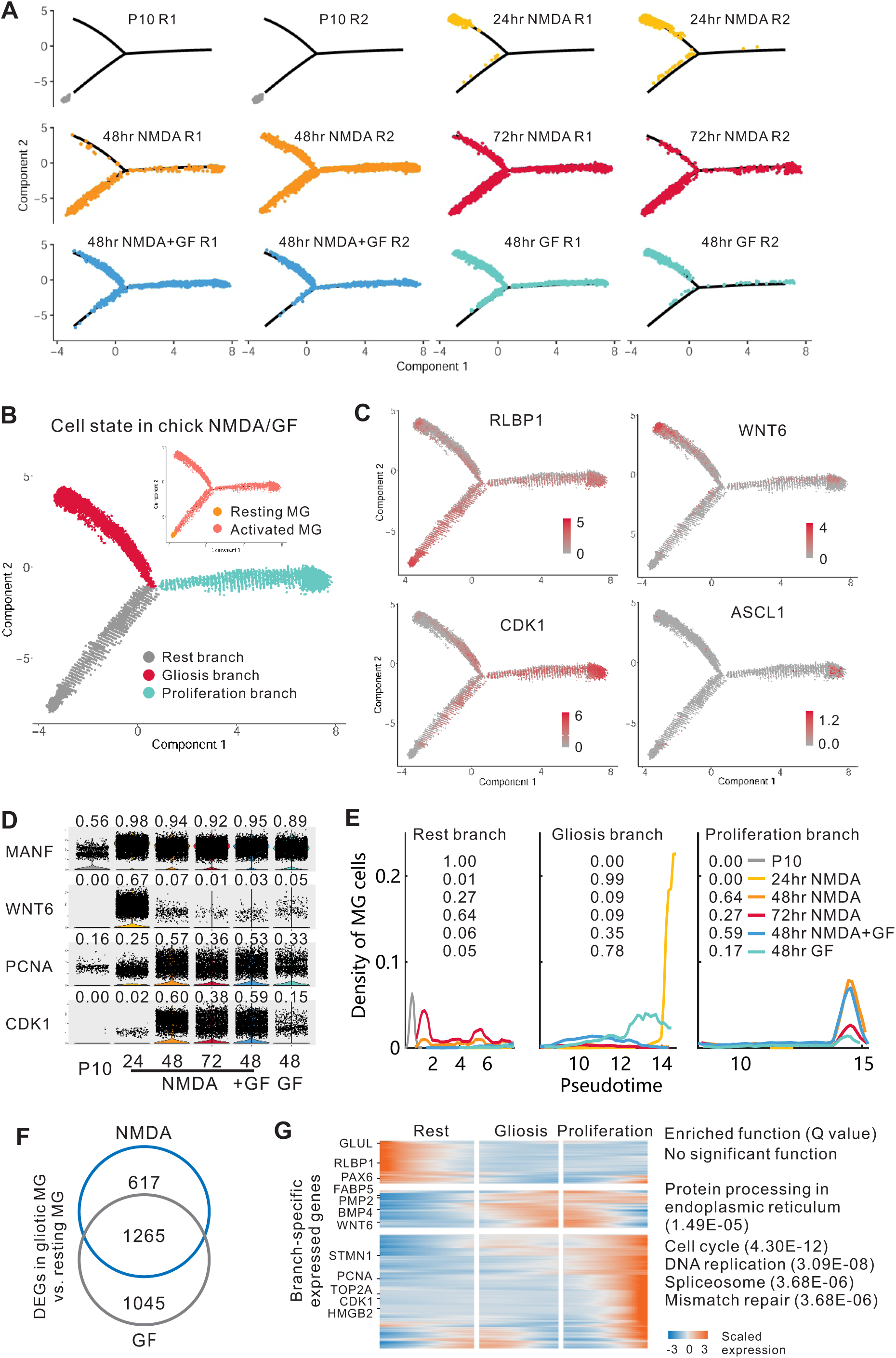
Single-cell trajectory and features of chick MG. **(A)**Distribution of chick MG from each sample in the trajectory. **(B)** Cell state of chick MG following NMDA /GF treatment. Inserted figure shows the distribution of resting and activated MG in the trajectory. **(C)** Expression of *RLBP1*, *WNT6*, *CDK1* and *ASCL1* in chick MG in the chick single-cell trajectory. **(D)** Expression of R*LBP1*, *WNT6*, *CDK1* and *ASCL1* in chick MG in each sample. Fraction of expressed cells is shown. **(E)** The distribution density of chick MG in the three branches across pseudotime. **(F)** Venn diagram of the number of differentially expressed genes (DEGs) from activated MG vs. resting chick MG between NMDA and GF treatment. **(G)** Heatmap and enriched functions of branch-specific expressed genes in chick MG. GF, growth factor (FGF2 + insulin) treatment.

**Fig. S11.**
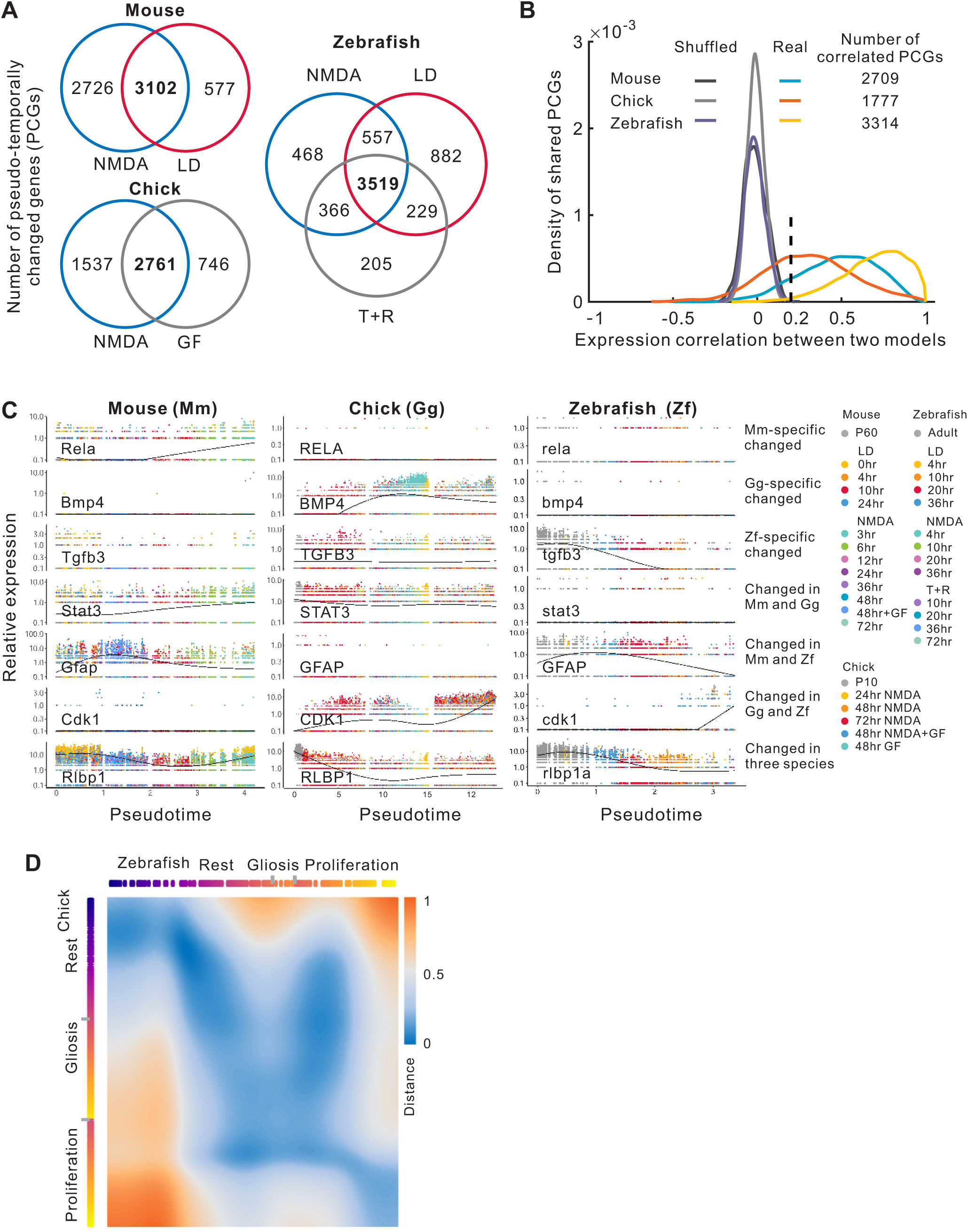
Comparison of pseudo-temporally changed genes (PCGs) from mouse, chick and zebrafish MG. **(A)**Venn diagram of the number of PCGs from different treatments in three species (fraction of expressed cells > 0.01, single-cell expression difference > 0.1 and q-value < 0.001). **(B)** Distribution of PCGs correlation between treatment models. Correlated PCGs(cPCGs) are refined from PCGs shared by all models in each species.**(C)** Examples of species-specific or shared cPCGs. **(D)** Alignment of two MG single-cell trajectories from chick and zebrafish. Colors of the branches in the trajectory indicate pseudotime. LD, light damage; T+R, TNFa+RO4929097 treatment; GF, growth factor (insulin+FGF2) treatment; Gg, Gallus gallus (chick.)

**Fig. S12.**
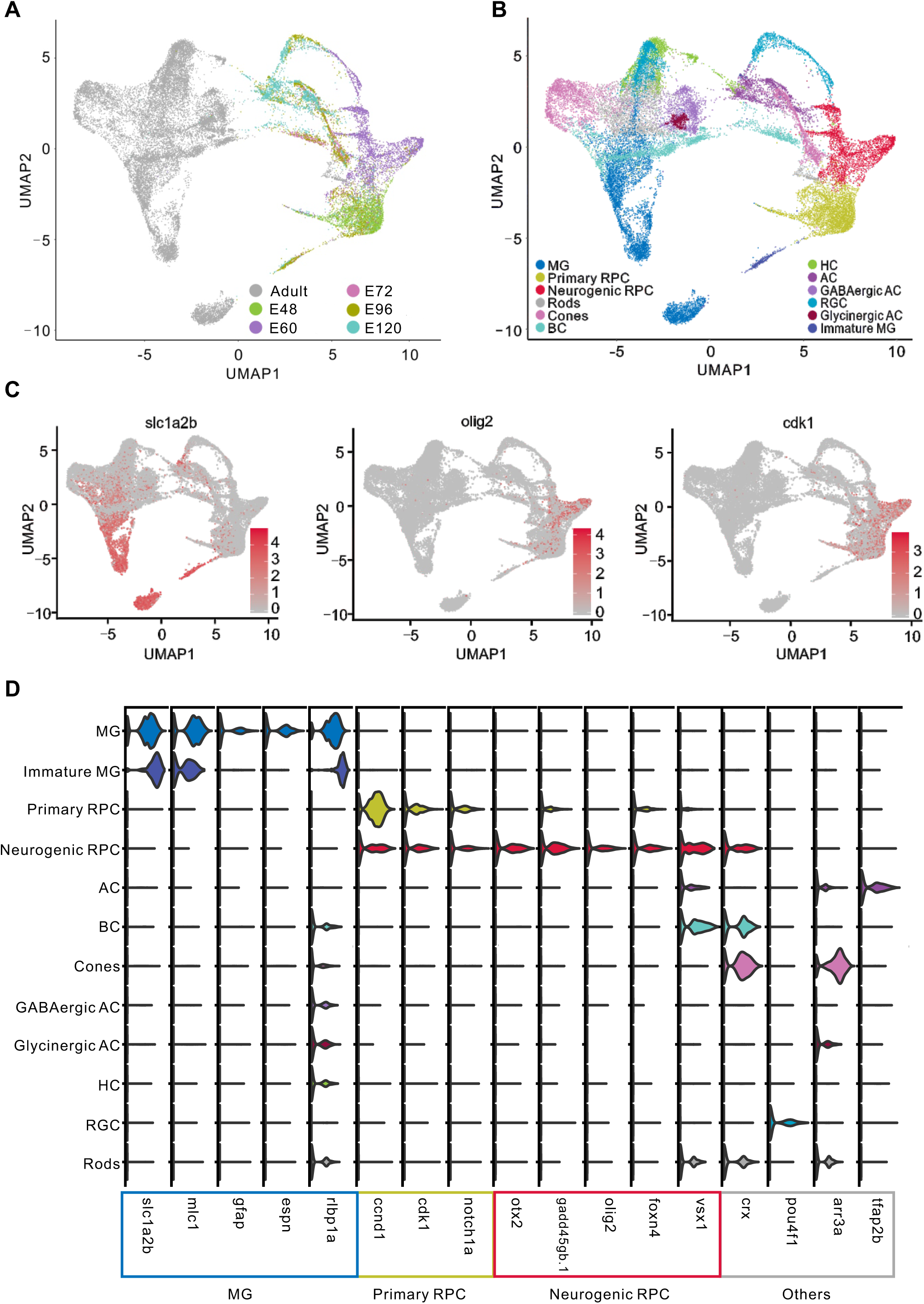
Clustering of MG cells from zebrafish development. **(A)**UMAP view of retinal samples collected at different timepoints during zebrafish development. **(B)** Clustering of retinal cell types during zebrafish development. **(C)** Single-cell expression of marker genes in the development of zebrafish RPC/MG. **(D)** Violin plot of the expression of known marker genes over the course of development of zebrafish RPC/MG. RPC, retinal progenitor cell; MG, Müller glia; BC, bipolar cell; AC, amacrine cell; RGC, retinal ganglion cell.

**Fig. S13.**
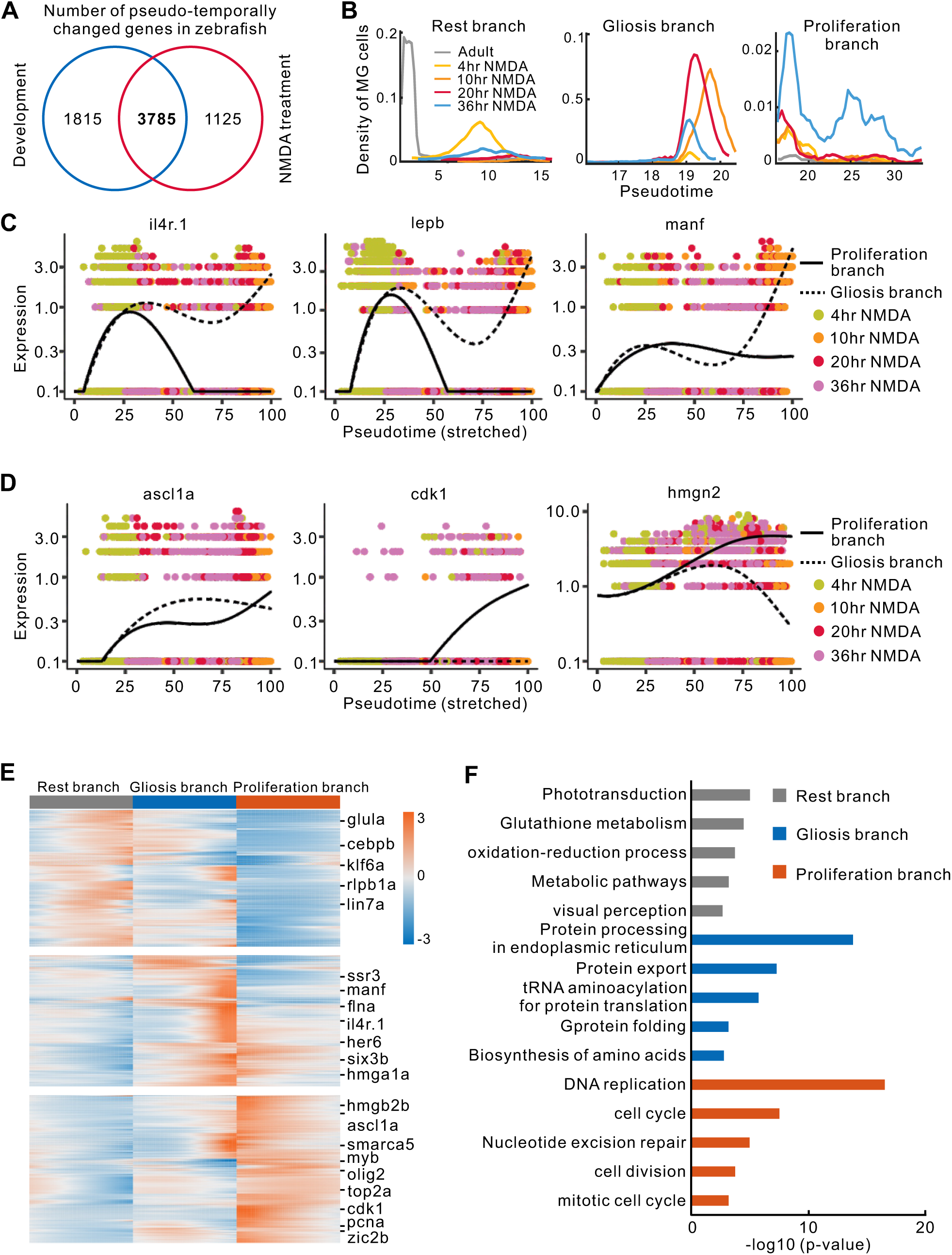
Analysis of branch-specific expressed genes in zebrafish MG following NMDA treatment. **(A)**Venn diagram of the number of pseudo-temporally changed genes in zebrafish MG during development and NMDA treatment. **(B)** The density of zebrafish MG across pseudotime after NMDA treatment. **(C)** Pseudo-temporal expression of genes enriched in the gliosis branch. **(D)** Pseudo-temporal expression of genes enriched in the proliferation branch. **(E)** Heatmap of genes specifically expressed in each of the three branches. **(F)** Top functions of genes enriched in three branches.

**Fig. S14.**
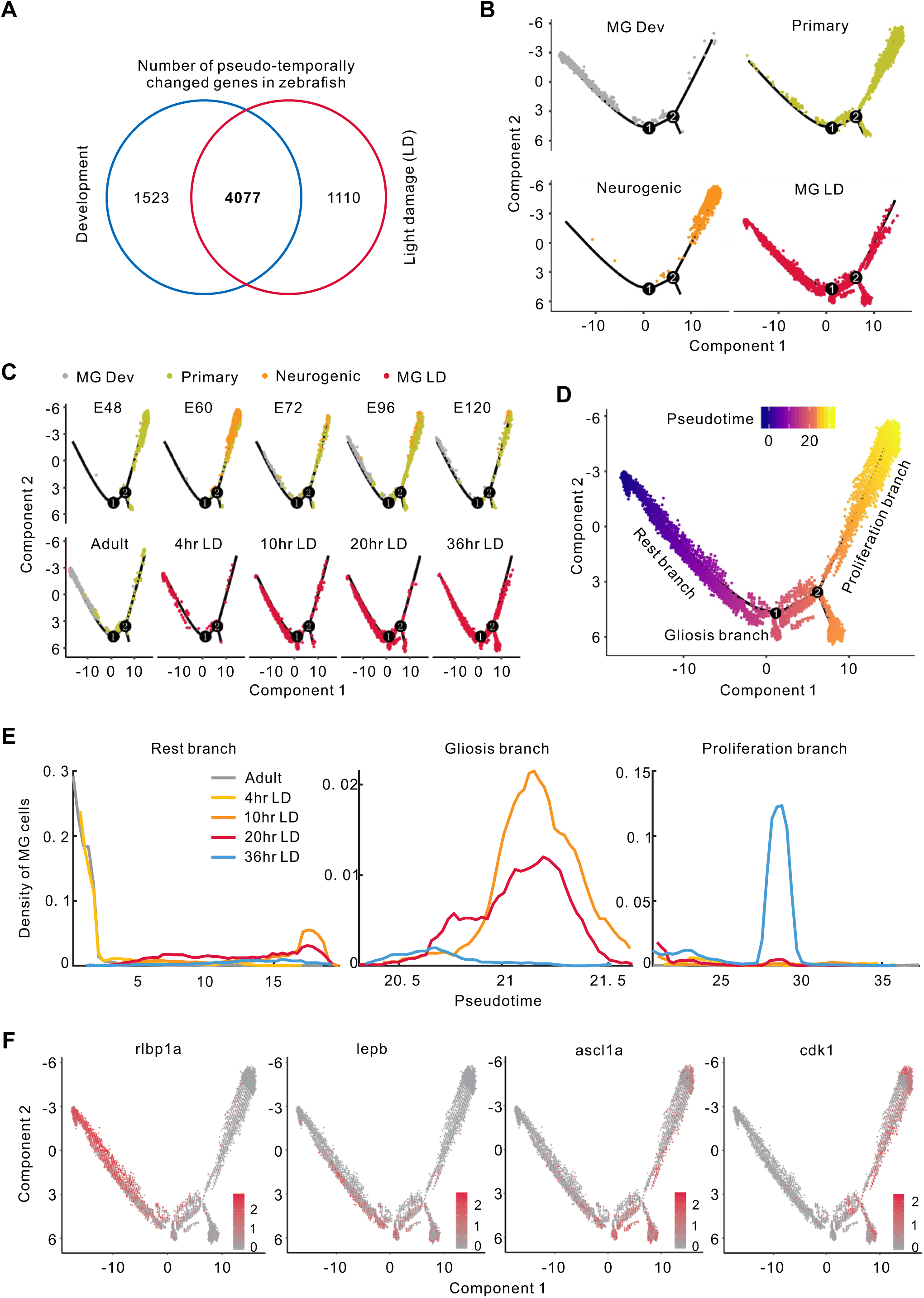
RPC/MG trajectory analysis for zebrafish development and light damage. **(A)**Venn diagram of the number of pseudo-temporally changed genes in MG during zebrafish development and light damage (LD). **(B)** RPC/MG trajectory from zebrafish development and light damage separated by cell types. **(C)** RPC/MG trajectory from zebrafish development and light damage separated by real time. **(D)** RPC/MG pseudotime analysis during zebrafish development and following light damage. **(E)** The density of MG across pseudotime of zebrafish light damage. **(F)** Single-cell expression of rest/gliosis/proliferation-related genes in light damage samples. RPC, retinal progenitor cell; MG, Müller glia.

**Fig. S15.**
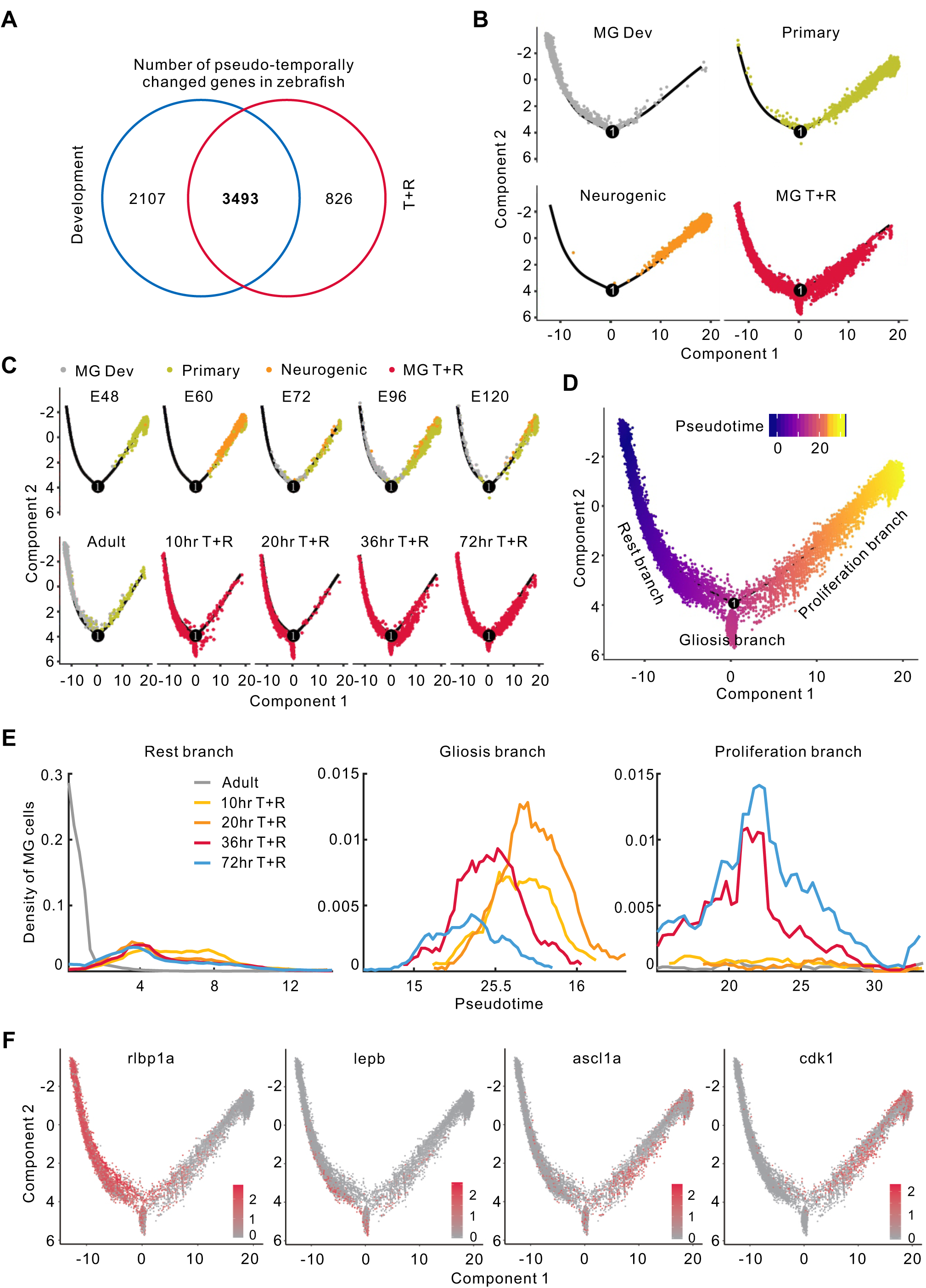
RPC/MG trajectory analysis for zebrafish development and TNFa+RO4929097 (T+R) treatment. **(A)**Venn diagram of the number of pseudo-temporally changed genes in MG during zebrafish development and T+R treatment. **(B)** RPC/MG trajectory from zebrafish development and T+R treatment separated by cell types. **(C)** RPC/MG trajectory of zebrafish development and T+R treatment separated by real time. **(D)** RPC/MG trajectory and pseudotime of zebrafish development and T+R treatment. **(E)** The density of MG across pseudotime of zebrafish T+R treatment. **(F)** Expression of rest/gliosis/ proliferation-related genes following T+R treatment. RPC, retinal progenitor cell; MG, Müller glia.

**Fig. S16.**
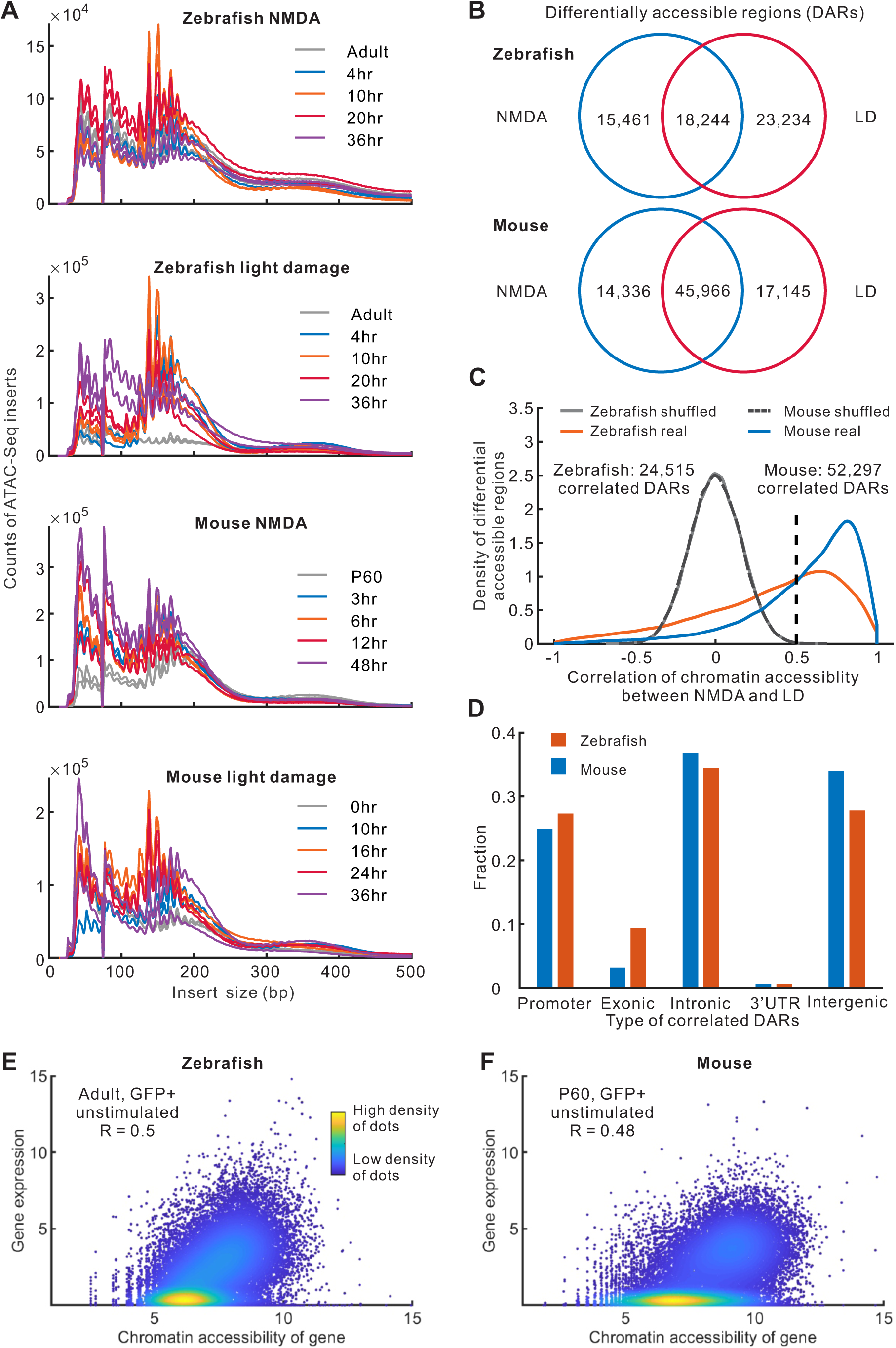
Identification of differentially accessible regions (DARs) from ATAC-Seq data. **(A)**Distribution of ATAC-Seq inserts. **(B)** Venn diagram of number of DARs in NMDA treatment and light damage (LD) of zebrafish and mouse. **(C)** Distribution of DAR correlations between NMDA and LD models. **(D)** Fraction of five types of correlated DARs. 3’UTR, three prime untranslated region. **(E, F)** Correlation of gene expression and corresponding chromatin accessibility in zebrafish (**E**) and mouse (**F**). Each dot represents one gene. The highest intensity of open chromatin regions was chosen for each gene.

**Fig. S17.**
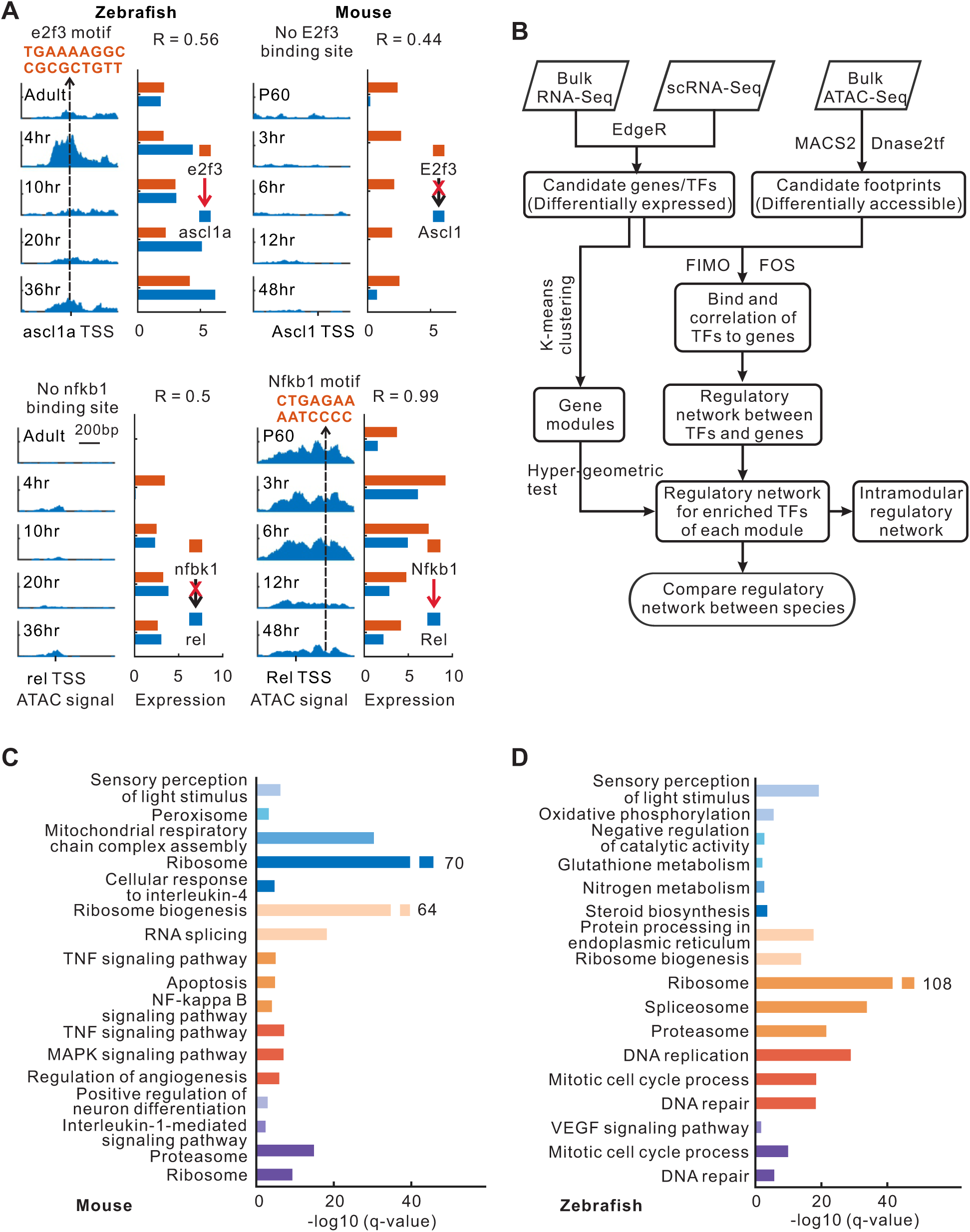
Changes in chromatin accessibility and gene expression. **(A)**Changes in chromatin accessibility and gene expression for two regulatory gene pairs of *e2f3/ascl1a* in zebrafish and *Nfkb1/Rel* in mouse. TSS, transcription start site. Expression correlation (R) of the gene pair was shown. **(B)** Workflow of Integrated Regulatory Network Analysis (IReNA) in integrating gene expression and chromatin accessibility to reconstruct the regulatory network (see Methods). Software EdgeR, MACS2, Dnase2tf and FIMO were used in IReNA. FOS, footprint occupancy score. TFs, transcription factors. **(C, D)** Most highly enriched functions of each gene module in zebrafish and mouse. Colors represent gene modules in Fig. 5G, H.

**Fig. S18.**
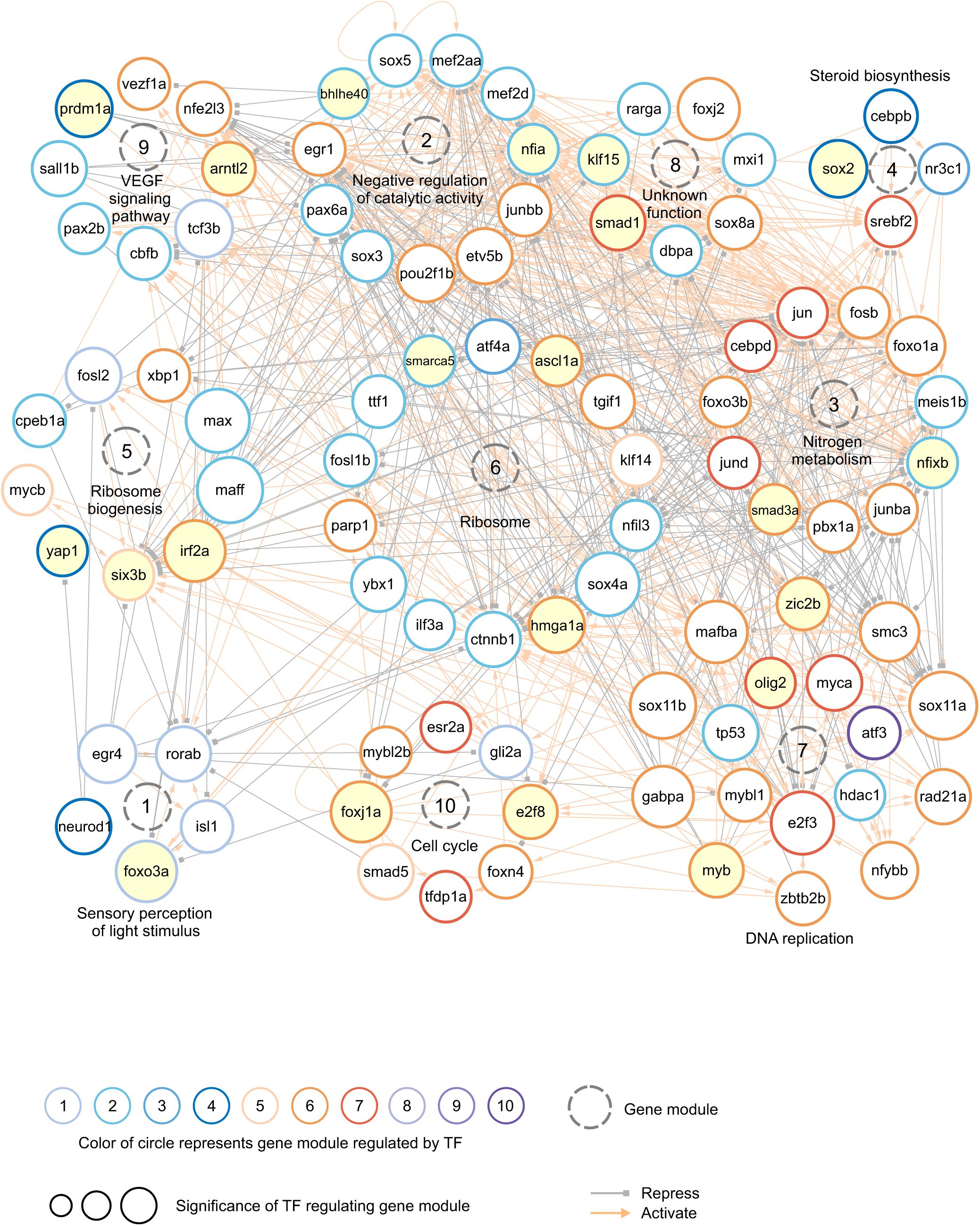
Regulatory network of enriched TFs in the zebrafish. All 97 enriched TFs are separated into ten modules based on the TF expression pattern from Fig. 5G. The color of the circle (TF) represents the module (Fig. 5G) that is most significantly regulated by TF. Circles filled in yellow indicate key regulators.

**Fig. S19.**
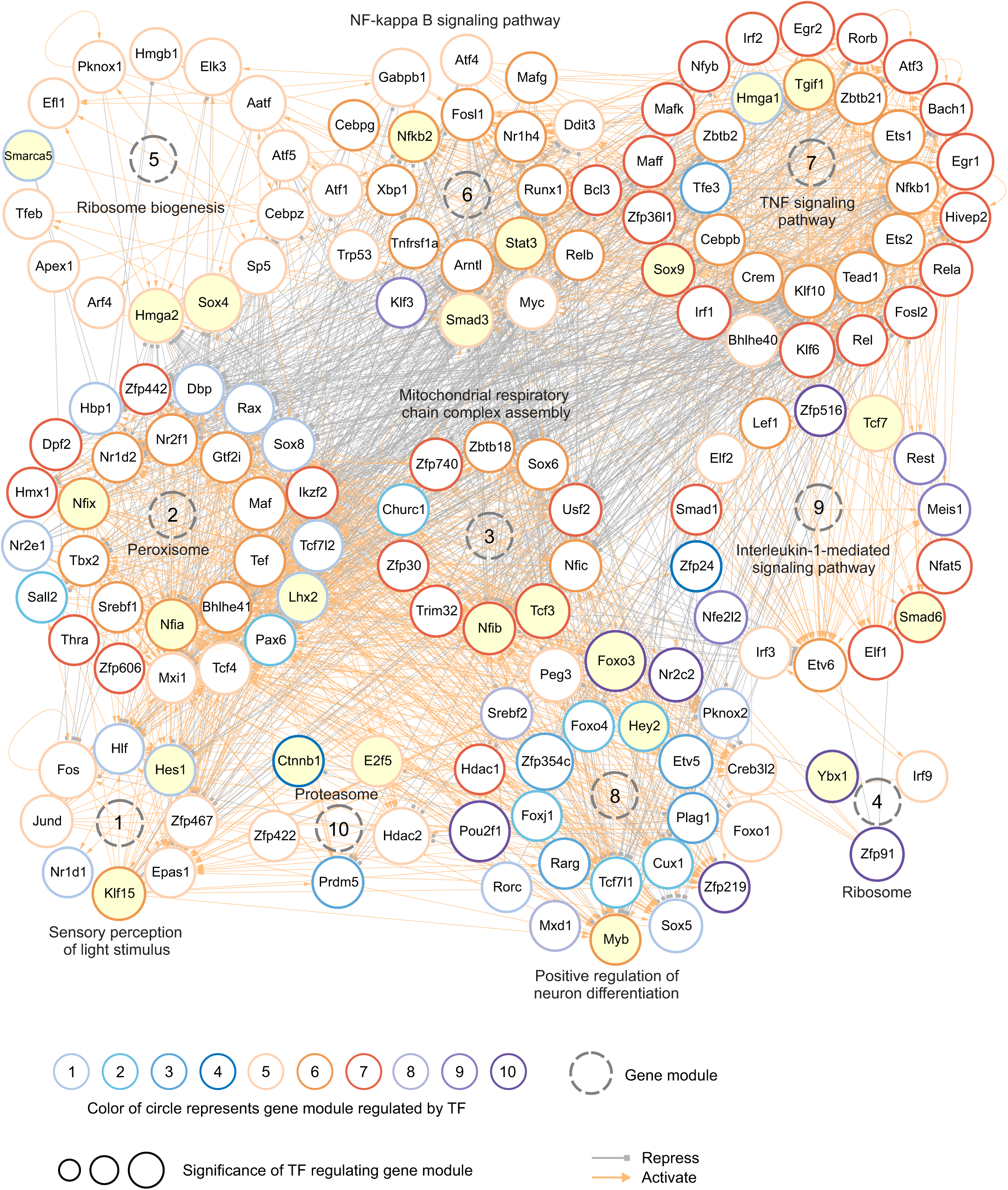
Regulatory network of enriched TFs in the mouse. All 156 enriched TFs are separated into ten modules based on the TF expression pattern from Fig. 5H. The color of the circle (TF) represents the module (Fig. 5H) that is most significantly regulated by TF. Circles filled in yellow indicate key regulators.

**Fig. S20.**
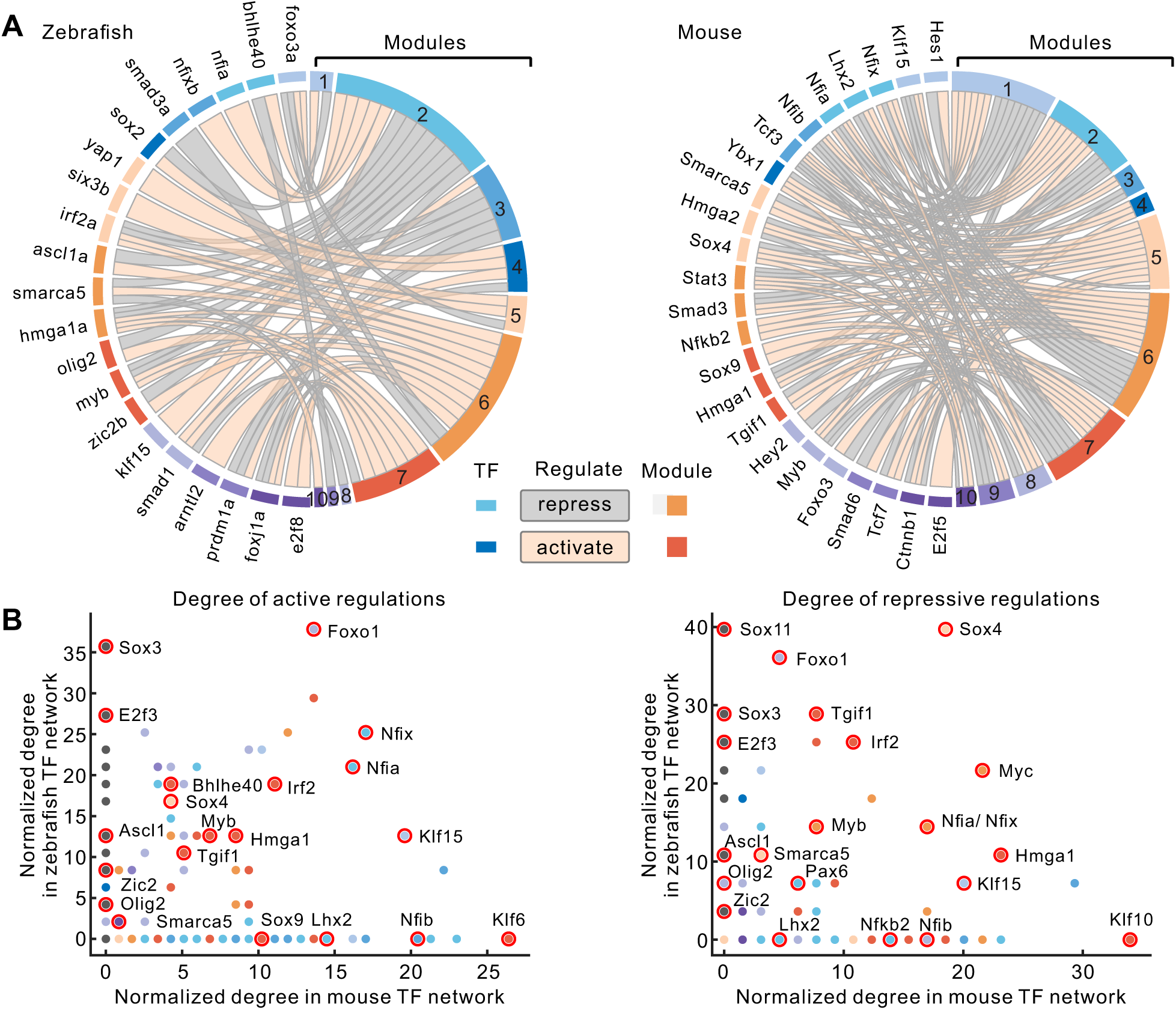
Features of enriched TFs. **(A)**Significant regulations between TFs and modules in zebrafish and mouse. Color of each TF indicates the module of TF assigned through k-means clustering in Fig. 5G, H. **(B)** Comparison of the regulation degree of enriched TFs between mouse and zebrafish. Colo r of dots represents the module of TFs in the mouse. Black dot indicates that the TF is not enriched in TF networks observed in the mouse. Dozens of key regulators are highlighted in red circles.

**Fig. S21.**
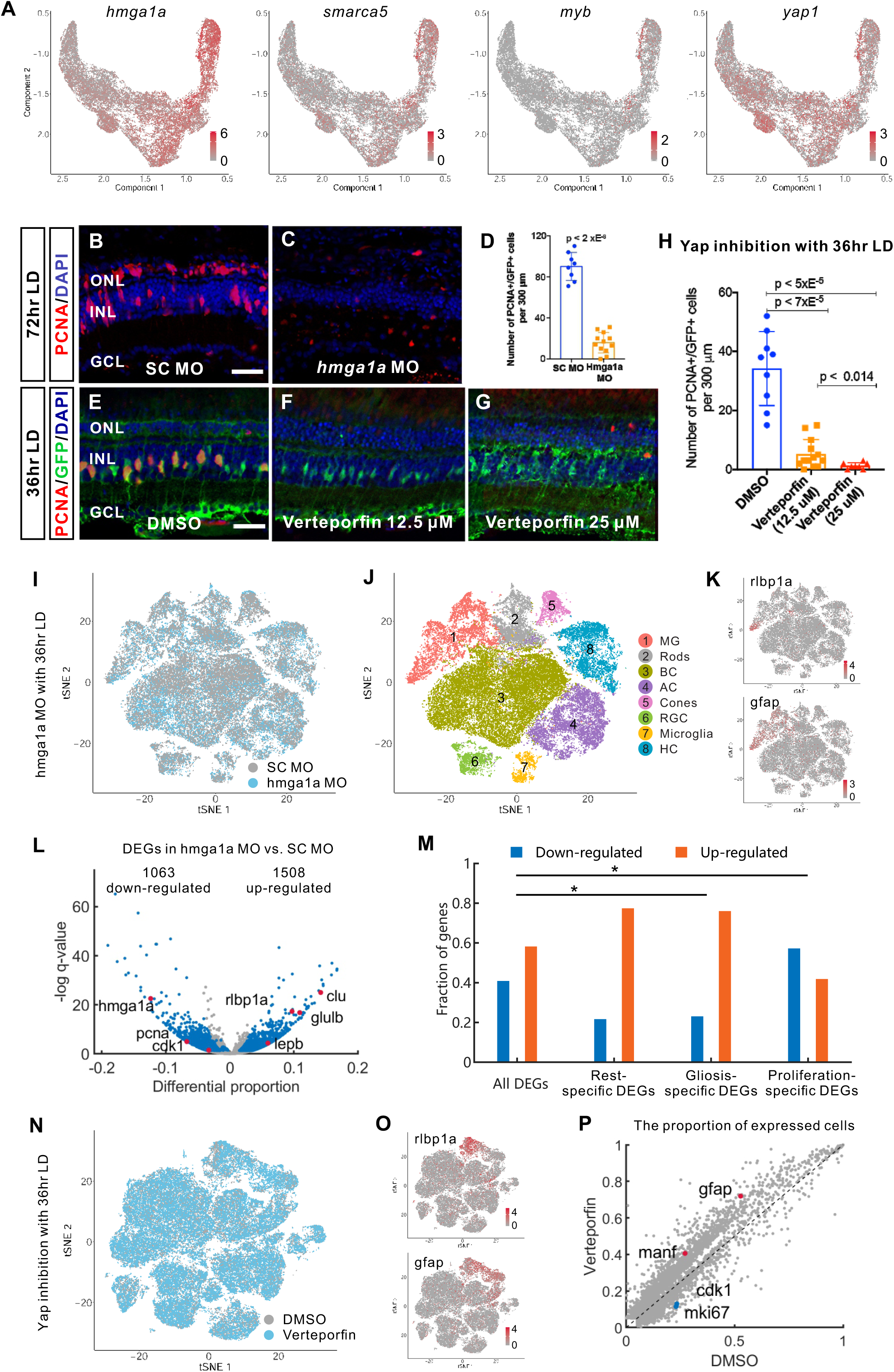
Candidate genes that regulate zebrafish MGPC formation in the damaged retinas. **(A)**Expression of candidate genes in zebrafish MG trajectory from NMDA, light damage and TNFa+RO4929097 (T+R) treatment. **(B, C)** Immunostaining for PCNA expression in standard control and *hmga1a* morphant retinas after 72 hrs of light treatment. **(D)** Quantification of the number of PCNA-positive INL cells in standard control and *hmga1a* morphant retinas at 72 hrs of light damage. **(E-G)** Immunostainings for PCNA and GFP expression in DMSO control and Yap inhibitor (verteporfin)-treated *albino*; *Tg[gfap:EGFP]nt11* retinas after 36 hrs of light damage. **(H)** Quantification of the number of PCNA and GFP colabeled INL cells in DMSO control and verteporfin-treated retinas after 36 hrs of light damage. Scale bars in panels **B** and **D** represent 20 μm. **(I, J)** t-SNE view of aggregated samples (**I**) and retinal cell types (**J**) in standard control and *hmga1a* morphant-treated retinas at 36 hrs of light damage. **(K)** Single-cell expression of MG-specific markers. **(L)** Volcano plot of differentially expressed genes (DEGs) in MG following *hmga1a* morpholino treatment as compared with the standard control morpholino. Differential expression was measured using the differential proportion of expressed cells. **(M)** Changes of branch-specific genes in MG following *hmga1a* morphant treatment. **(N)** t-SNE view of aggregated samples from DMSO and Yap inhibitor (verteporfin) treatments after 36 hrs of light damage. **(O)** Single-cell expression of MG-specific markers from DMSO and verteporfin-treated retinas. **(P)** Differentially expressed genes (DEGs) in MG following verteporfin treatment as compared with DMSO control.

**Fig. S22.**
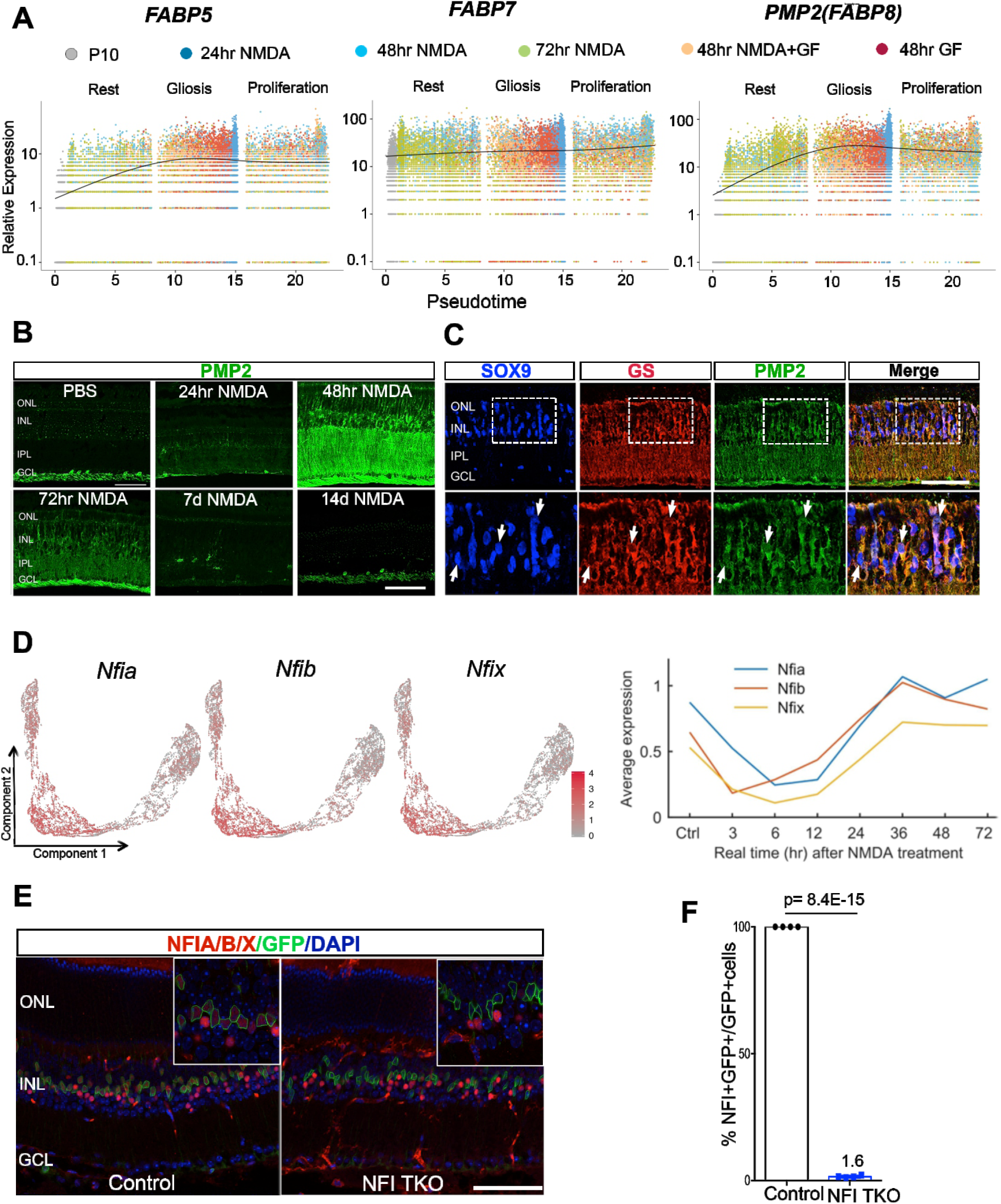
Expression of *FABP5, FABP7* and *PMP2 (FABP8)* was induced in chick MG after NMDA injury. **(A)**Pseudo-temporal expression of *FABP5, FABP7* and *PMP2* in the chick MG during NMDA damage or growth factor treatment. **(B)** Immunohistochemistry shows a transient upregulation of expression of PMP2 protein at 48 and 72 hr in chick retina after NMDA injury. **(C)** Increased expression of PMP2 in the chick MG as indicated by co-localization of PMP2 with SOX9 and GS proteins (white arrows). **(D)** Expression of *Nfia/b/x* in mouse MG in the single-cell trajectory and real time after NMDA injury. **(E, F)** Immunohistochemistry and quantification indicate a MG-specific deletion of *Nfia/b/x* in adult mouse (NFI TKO) retina as compared with the control. Scale bar, 50 ì m. GF, growth factor (FGF2+insulin).

**Fig. S23.**
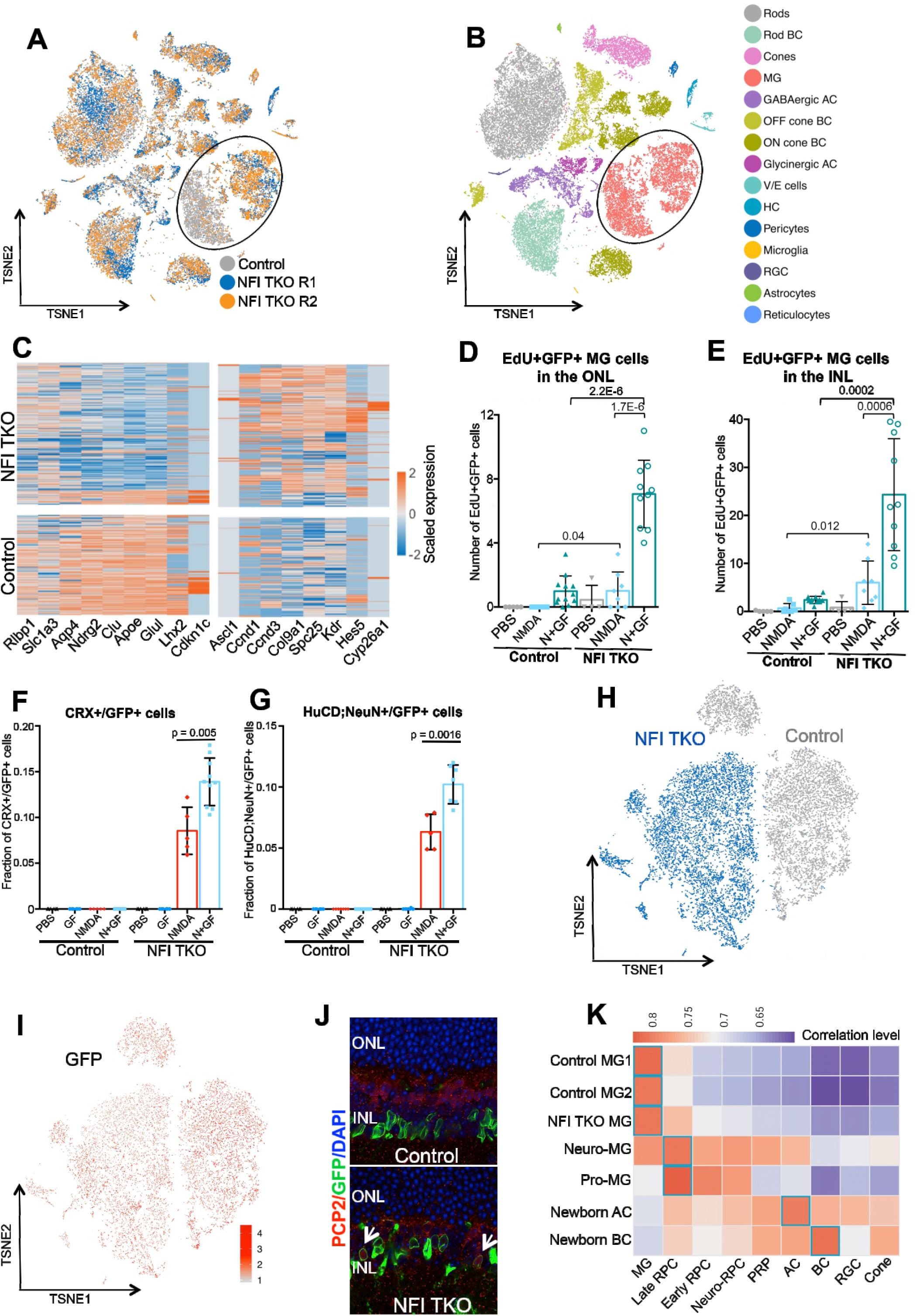
Loss of Nfia/b/x led to downregulation of MG markers, upregulation of cell cycle-related genes and generation of neurons from MG after injury. **(A)** tSNE view of single cells from the aggregation of two NFI TKO retinal samples and a control retinal sample. **(B)** tSNE view of cell types from the aggregation of two NFI TKO and one control retinal samples shows that MGs from NFI TKO samples are clearly distinct from the control sample. **(C)** Single-cell heatmap shows a set of differentially expressed genes (DEGs) of the MG from the two mouse genotypes. **(D, E)** Quantification of MG proliferation in the ONL and INL after 3 days of intravitreal injection of NMDA with and without growth factors. (**F**, **G**) Quantifications of CRX+ and HuCD;NeuN+ newborn neurons derived from MG in control and *Nfia/b/x* knockout retina at day 14 after injury. **(H)** tSNE view of single cells from the aggregation of NFI TKO and control GFP-positive retinal cells flowed-sorted at D21 after NMDA and growth factor injection. **(I)** GFP mRNA expression in scRNA-Seq data from flow-sorted cells. **(J)** Expression of PCP2 was confirmed by immunostaining in MG-derived bipolar cells (white arrows) from *Nfia/b/x-*deficient MG. **(K)** Comparison of MG-derived neurons to native retinal cells from the mouse (Clark, et al. 2019). Expression correlation between cell types was calculated using 9489 genes that have > 0.1 normalized expression in scRNA-Seq. For each type of cells from control and NFI TKO, the most similar cell types are highlighted in the cyan box. ONL, outer nuclear layer; INL, inner nuclear layer; GF, growth factor (FGF2 and Insulin); RPC, retinal progenitor cells; PRP, photoreceptor precursors; RGCs, retinal ganglion cells; BC, bipolar cells; AC amacrine cells.

